# Average miniature post-synaptic potential size is inversely proportional to membrane capacitance across neocortical pyramidal neurons of different sizes

**DOI:** 10.1101/2024.07.05.602190

**Authors:** Martynas Dervinis, Guy Major

## Abstract

In chemical synapses of the central nervous system (CNS), information is transmitted via the presynaptic release of a vesicle (or ‘quantum’) of neurotransmitter, which elicits a postsynaptic electrical response with an amplitude termed the ‘quantal size’. Measuring amplitudes of miniature postsynaptic currents (mPSCs) or potentials (mPSPs) at the cell soma is generally thought to offer a technically straightforward way to estimate quantal sizes, as each of these miniature responses (or minis) is generally thought to be elicited by the spontaneous release of a single neurotransmitter vesicle. However, in large highly-branched neurons, a somatically recorded mini is typically massively attenuated compared with at its input site, and a significant fraction are indistinguishable from (or cancelled out by) background noise fluctuations. Here, using a new software package (also) called ‘minis’, we describe a novel quantal analysis method that estimates the ‘electrical size’ of the synapse by comparing events detected in somatic recordings from the same neuron of (a) *real* minis and (b) background noise (with minis blocked pharmacologically) with *simulated* minis added by a genetic algorithm. The estimated minis’ distributions reveal a striking inverse dependence of mean excitatory mPSP amplitude on total cell membrane capacitance (proportional to cell size, or more exactly, extracellular membrane surface area) suggesting that, in rats at least, the average charge injected by single excitatory synapses (ca. 30 fC) is conserved across neocortical pyramidal neurons of very different sizes (across a more than three-fold range).

## Introduction

Synaptic transmission in the CNS occurs in ‘quanta’ (Forti et al., 1997; Hardingham et al., 2010; Isaacson and Walmsley, 1995; Jonas et al., 1993; Kraszewski and Grantyn, 1992; Kullmann and Nicoll, 1992; Kuno, 1964; Larkman et al., 1991; Sahara and Takahashi, 2001; Stern et al., 1992; Wall and Usowicz, 1998), usually thought to each correspond (for the most part at least) to the release of a single neurotransmitter vesicle (Heuser et al., 1979). The amplitude of the postsynaptic response to a quantum of transmitter, the quantal size, is a key parameter defining the efficacy of synaptic transmission (Katz, 1969; Vere-Jones, 1966) and, therefore, an important determinant of neural computation. For example, how many synapses, on average, does it take to fire a cell?

Quantal size measurements in the CNS are difficult, due to multi-synaptic contacts (multiple synapses per connection between a particular pair of neurons, and multiple release sites/postsynaptic densities/receptor ‘rafts’ for some synapses), limited accessibility, significant quantal variance (both within and between synapses of a given connection), temporal non-stationarity of synaptic properties (e.g. receptor desensitisation or internalisation, and depletion of glutamate from presynaptic vesicles), and low signal-to-noise ratios of somatic electrophysiological recordings of quantal events (Bekkers, 1994; Farsi et al., 2021; Korn and Faber, 1991). For these reasons classical quantal analysis as done at the neuromuscular junction often runs into problems in central synapses, where most of the minis do not ‘stand out’ from the noise. More recent methods such as noise deconvolution, Variance-Mean analysis, and optical quantal analysis have not fared much better (Biró et al., 2005; Clamann et al., 1991; Clements, 2003; Clements and Silver, 2000; Edwards et al., 1976; Enoki et al., 2009; Jack et al., 1981; Liu et al., 2022; Oertner et al., 2002; Peled and Isacoff, 2011; Redman, 1990; Reid and Clements, 1999; Silver, 2003; Yuste et al., 1999), or suffer from potential selection biases (e.g. paired recordings typically require cells close together whose synapses may be a non-representative sub-population, differing from the overall population of synapses).

A complementary method of assessing synaptic function is to measure properties of mPSCs. Miniature PSC amplitude measurements are often taken as quantal size estimates across the population of active synapses onto a neuron, as the mPSC (under voltage clamp) or mPSP (under voltage recording) likely corresponds (in most cases) to the response to a spontaneous release of a single neurotransmitter vesicle (Brown et al., 1979; del Castillo and Katz, 1954; Fatt and Katz, 1952; Isaacson and Walmsley, 1995; Wall and Usowicz, 1998) and, therefore, if performed accurately, can be used as the basis for constructing a relatively straight-forward quantal analysis method (of the population of synapses onto a given neuron). Measurements of mPSC amplitudes are routine in the literature. However, they are likely overestimates due to small amplitude events being difficult to detect and largely being missed or lost in the noise.

The use of voltage clamp can be a major reason why mPSCs ‘go missing’. Theoretical studies (Jonas et al., 1993; Major, 1993; Rall and Segev, 1985; Spruston et al., 1993) predicted, and experimental findings (Williams and Mitchell, 2008) confirmed that somatic voltage clamp fails to space clamp extended dendrites, and to capture all (or even most of) the current (thus charge) flowing in at electrically remote synapses (which instead spreads out over the entire cell membrane capacitance). Amplitude attenuation can also be caused if series resistance (R_ser_) is not adequately compensated (Major, 1993; Williams and Mitchell, 2008). These issues can be somewhat sidestepped by switching to voltage recording, at the cost of (a little) more temporal integration by the membrane (thus overlap between minis). Although some charge is inevitably going to escape by leaking across the dendritic membrane no matter the recording method, charge escape through the relatively low membrane conductance is a much slower process than charging of the neuronal membrane via the far larger axial cytoplasmic conductance (Jonas et al., 1993; Major, 1993). In fact, it is so slow, relatively, that just about all the synaptic current would be integrated onto the membrane capacitance and, thus, most of the synaptic charge is ‘captured’ by the voltage recording (current clamp), as the PSP peak amplitude, before then leaking out slowly via membrane ion channels.

However, during voltage *clamp*, most mPSCs go undetected because of the electrophysiology of the neuron itself. Distal minis leak some charge via the dendritic membrane, but a far more significant amplitude attenuation, 40-fold or more, is inflicted by charging the membrane capacitance of the entire dendritic arbour (Larkum et al., 2009; Major et al., 2013; Nevian et al., 2007; Stuart and Spruston, 1998; Williams and Mitchell, 2008; Williams and Stuart, 2002). Such amplitudes, at the soma (approximately 80 µV, on average, but as small as 30 µV, in layer 5 pyramidal neurons under *current* clamp (Nevian et al. 2007), detected via simultaneous dendritic whole-cell recordings) are often indistinguishable from noise fluctuations. Recording from thin dendrites close to synaptic input sites is technically extremely challenging – a tour de force beyond most labs. Therefore, a reliable yet widely accessible quantal analysis method, based on standard somatic whole-cell patch recording, would require taking account of the background noise in addition to measuring putative mPSPs (or mPSCs).

Here we describe such a method, based on recording excitatory mPSPs (mEPSPs) in the presence of background physiological noise (‘noise with minis’) then noise alone, from the same cell, after blocking mEPSPs pharmacologically. To detect mEPSP-like events in these recordings, we used a newly developed algorithm called ‘minis’ which is described in our companion article (Dervinis and Major, 2024). Because of small potentials being ‘overshadowed’ by larger minis, either by summating with them on rising phases, or having their amplitudes reduced on decaying phases (analysed and discussed in detail in our companion article (Dervinis and Major, 2024)), events detected in the ‘noise-alone’ recording cannot be used to remove the noise component in the ‘noise with minis’ recording simply by subtracting their two distributions. Instead, we adopted the approach of using simulated minis, superimposing them on the ‘noise-alone’ recording and varying their parameters until the amplitude and rise time distributions of events *detected* in the ‘noise with simulated minis’ voltage trace closely matches those detected in the ‘noise with real minis’ recording, with identical detection parameters. Therefore, we augmented our ‘minis’ software with a genetic optimisation algorithm (GA) that automatically adjusts amplitudes, rise times and frequencies (rates) of simulated minis until a statistically indistinguishable match is achieved. This novel quantal analysis method produced an unbiased ‘upper limit’ average quantal mEPSP amplitude for each neuron, which unexpectedly turned out to be inversely proportional to membrane capacitance. This in turn allowed us to obtain a population ‘average quantal charge’ estimate for excitatory neocortical synapses in the rat of approximately 31 pC. This was conserved across pyramidal neurons of all sizes in (somatosensory) cortical layers 2/3 and 5, across a more than 3-fold range. ‘Minis’ thus provides an experimentally straightforward ‘quantal analysis’ method (in the broadest sense) for detecting mPSPs (or indeed mPSCs) in somatic recordings, and (in aggregate) ‘separating’ the smaller of these events from similar-sized background noise fluctuations.

## Materials and Methods

### Animals and electrophysiology

All experimental procedures were carried out in accordance with the UK Animals (Scientific Procedures) Act 1986. The full description is provided in our companion article (Dervinis and Major, 2024). Coronal somatosensory neocortical slices obtained from 19- to 27-day old Wistar rats (RRID:RGD_13508588; n = 11) were used to obtain whole-cell voltage recordings (n = 14) of mEPSPs in the background of physiological (and other) noise (‘noise with minis’ condition), by superfusing artificial cerebrospinal fluid (aCSF) containing blockers of action potentials (APs) and inhibitory postsynaptic potentials (IPSPs). Excitatory postsynaptic potential (EPSP) blockers were then added (‘noise-alone’ condition, with all glutamatergic, GABAergic and AP-dependent synaptic potentials blocked, i.e. the vast majority of fast synaptic inputs in the neocortex), and background noise traces were recorded from each cell, while maintaining stability of the whole-cell recording (as judged by numerous criteria, below).

Passive membrane time constant, total membrane capacitance, input resistance, and bridge balance (R_ser_ compensation) error were estimated by injecting current pulses as described in (G. Major et al., 1993).

### Methodology

The goal of the study was to design an experimentally straightforward quantal analysis method based on standard somatic recordings of minis, as opposed to far more challenging recordings from sub-micron diameter input dendrites closer to (a few of) the synapses. Due to their low signal-to-noise ratio, *simulated* minis were used to infer the signal (mEPSP) component and to eliminate (in aggregate) the noise component in these recordings.

The experimental design of this study was similar to the one described in the preceding companion article (Dervinis and Major, 2024). As described in the previous section, whole-cell recordings were obtained where first mEPSPs were recorded in the background of physiological and other noise followed by blocking EPSPs pharmacologically to record ‘pure’ noise in the absence of minis. Simulated mEPSPs (smEPSPs) were then randomly added to the noise traces (Figure 1) and the ‘minis’ algorithm was applied to this ‘noise with simulated minis’ voltage trace. The resulting distribution of detected events (in terms of their amplitudes and 10-90% rise times) was compared to the equivalent distribution based on the ‘noise with real minis’ recording, detected with the same parameters (such as amplitude detection threshold, baseline duration and maximum rise time gap till each peak). The discrepancy between the two distributions was used to inform the GA, which would then adjust the distribution of smEPSPs and generate new ‘noise with simulated minis’ voltage traces and compare them again to the equivalent distribution detected with the same parameters from the ‘noise with real minis’ recording. This procedure was repeated multiple times (of the order of tens to hundreds of generations with 260 examples per generation; 10 examples per each parameter optimised by the GA) until the two distributions were closely matched, within the bounds of chance statistical errors (details below). At the end of this process the minis distribution used for simulating the closely matched recording was deemed to be the inference of the true minis distribution underlying the ‘noise with real minis’ recording (or at least compatible with it).

**Figure 1:**
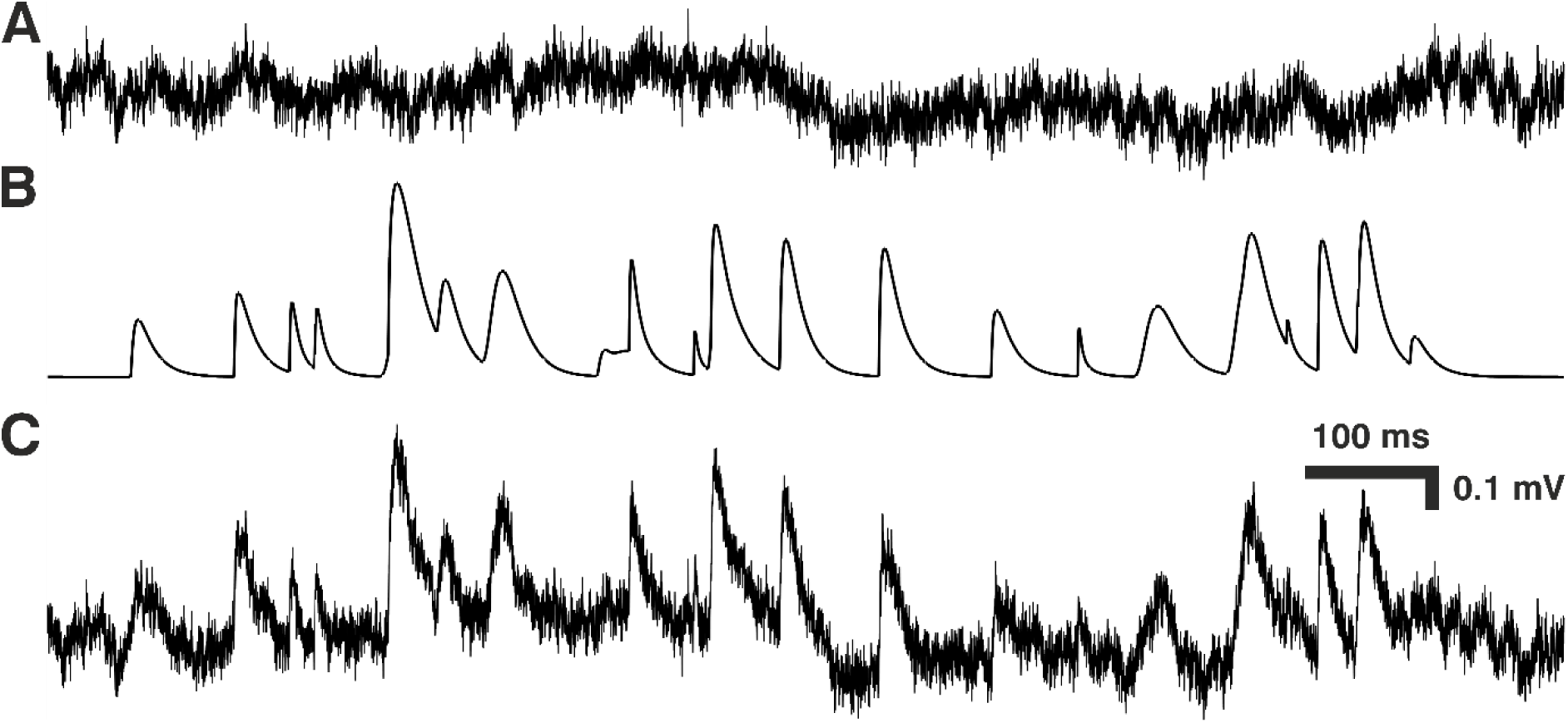
Constructing ‘noise with simulated minis’ voltage trace. (A) Segment of ‘noise-alone’ voltage recording (from cortical layer 2/3 pyramidal neuron p131a). (B) Simulated mEPSPs that were randomly added to the noise-alone trace in (A) to produce the ‘real noise with simulated mEPSPs’ voltage trace in (C).

### Detection of miniature postsynaptic potentials

Detection of real and simulated mEPSPs was carried out using our ‘minis’ software. The detailed description of this detection algorithm is provided in the companion article (Dervinis and Major, 2024). We used the same detection algorithm parameters as described in that article except for the amplitude range, which was set to 0.02-10 mV.

### Selection of recordings for fitting distributions of detected real minis-like events

Prior to carrying out any estimation of the distribution of minis, segments of ‘noise with real minis’ and ‘noise-alone’ recordings were carefully selected. Segments containing unusual amount of noise or V_m_ glitches were rejected. Parts of recording showing instability or drift of recording parameters (or excessive movement of the recording pipette relative to the soma) were also rejected, to ensure that only good quality data were used. Data analysis performed at this preprocessing/selection stage were built into our ‘minis’ software.

As part of the preprocessing stage, ‘minis’ produced seven graphs showing the evolution of various recording parameters across the entire duration of a single recording. An example (from the longest recording obtained) is provided in Figure 2, showing these graphs as individual panels. Figure 2A shows the evolution of V_m_ from the very beginning of the recording obtained in the regular aCSF with APs present (in this case) to the very end of the recording, where APs, mIPSPs and mEPSPs were all blocked for some time (‘noise-alone’). The data is shown after removing V_m_ segments containing responses to regularly injected current pulses and has a total duration of 8520 s. Throughout the entirety of this recording the baseline V_m_ varied between –70 and –67 mV and, thus remained relatively stable.

**Figure 2:**
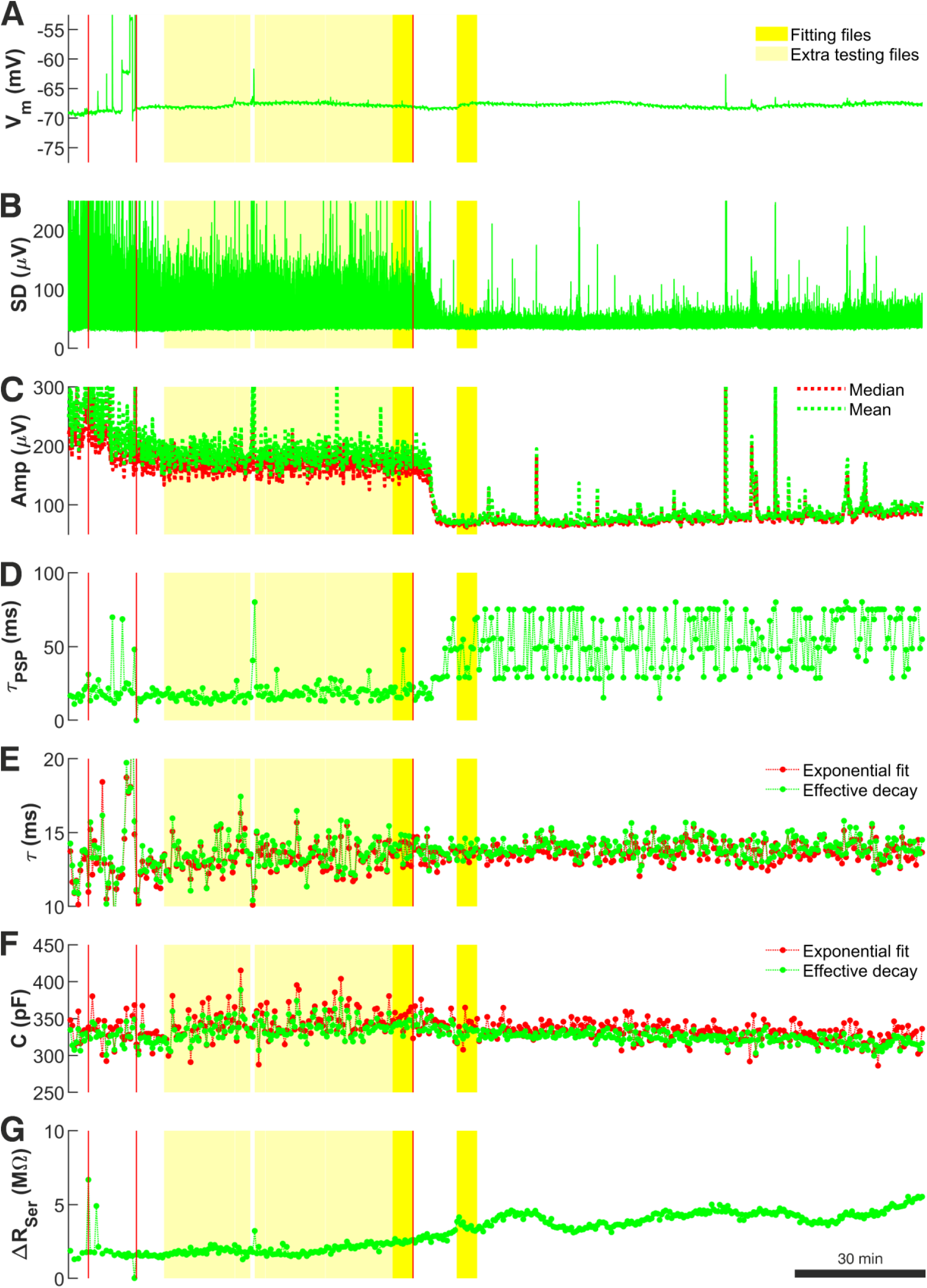
Recording quality measures used during the data preprocessing stage to select ’noise with minis’ and ’noise-alone’ recording sweeps for further data analysis for recording p103a (layer 5 ‘burster’ with tuft and thick trunk). (A) Baseline V_m_ across the entire recording session. Stimulation pulses were excluded from the display. The total recording duration (excluding pulse periods) was 8520 s. The first red vertical line marks the time when pharmacological blockade of APs was initiated. The second red vertical line marks the time when no more APs were observed in the recording. The third red vertical line marks the time when pharmacological blockade of mEPSPs was initiated. Bright yellow periods mark ‘noise with minis’ and ‘noise-alone’ recording segments selected as the main comparison files for the distribution fitting. Light yellow marks all the remaining good quality ‘noise with minis’ recording sweeps that were also included in the distribution fitting procedure for estimating target SAD, MAD, and ‘deviation’ scores, as well as statistical evaluation of fits (see Methods). (B) V_m_ standard deviation measured over consecutive 15 ms windows over the same recording session. (C) Evolution of mean and median amplitudes of the biggest top 10% of detected events (including minis and noise events), averaged over single sweeps (duration of each was 4 s). (D) Effective decay time constants (time to drop to 1/e of starting value ‘safely’ (8 ms) after the peak and upwardly convex part) of the same biggest 10% of detected events as in panel (C) averaged over consecutive 20-second-long (5-sweep-long) segments across the same session. Once minis were blocked pharmacologically, effective time constant of detected events, (now pure noise) increased, and its scatter, so this is another useful indication of when the block is complete. (E) Evolution of passive membrane time constant over the entire recording session. Measurements were averaged over consecutive 20-second-long (5-sweep-long) segments. Measurements based on exponential decay curve fit and on ‘effective time constant’ (time for V_m_ to drop to 1/e of a value just after change in current (after pipette capacitance artefact and fast charge redistribution components: starting at 4 ms after the end of the pulse). (F) Evolution of total cell membrane capacitance across the recording session. Measurements were averaged over consecutive 20-second-long (5-sweep-long) segments. Measurements based on exponential decay curve fit and on effective time constant (time taken for V_m_ drop to 1/e of an initial value of the slowest impulse response component). (G) Evolution of whole-cell pipette series resistance change (bridge balance error) across the recording session, calculated from any jumps at starts or ends of charging curves. Measurements were averaged over consecutive 4-second-long (5-sweep-long) segments.

There are three time markers that extend across all figure panels marking the time when the aSCF containing TTX, gabazine, and CGP55845A was infused to block APs and IPSPs (the first marker), the time when APs were observed to cease (the second marker), and the time when the aCSF containing NBQX and CPP was infused to block remaining EPSPs (the third marker). The recording segment between the second and the third markers constituted the ‘noise with minis’ recording, while the segment following the third marker *after* mEPSPs were eventually blocked constituted the ‘noise-alone’ recording. The darker yellow shaded regions that also extend across all panels of Figure 2 mark files (epochs) that were selected for subsequent minis distribution fitting.

The remaining panels of Figure 2 were used to guide data selection. The standard deviation (SD) of the V_m_ (Figure 2B) was useful in determining the transitions between different pharmacological conditions. Stable prolonged decreases in SD reliably indicated when APs or minis were effectively blocked. Sudden brief increases in SD flagged problematic recording segments containing higher levels of noise or V_m_ glitches. Gradual increases in SD suggested gradual deterioration of recording quality.

Similar information was also provided by the evolution of the biggest 10% of amplitudes of all detected minis-like events in a single sweep (Figure 2C). Prolonged drop in the amplitudes marked transitions between recording conditions, while sudden jumps were indicative of V_m_ instabilities. Finally, effective decay time constants of biggest 10% of detected events (Figure 2D) could be used as an indicator for the presence/absence of minis. Blocking minis resulted in time constants becoming large and fluctuating unstably, indicating that only noise fluctuations were being detected that did not possess the typical decay dynamics common to EPSPs.

Further insights into the recording quality over time were provided by V_m_ responses to brief current pulses. Passive membrane time constant measurements over time (Figure 2E) were expected to remain stable. Deviations from this could signal a change in passive membrane properties due to degradation in the recording quality. Total membrane capacitance (Figure 2F) was also expected to be stable over time. Change in the capacitance could indicate membrane tearing, patch-resealing or pipette movement or clogging, loss of membrane or part of the cell or some substantial uncontrolled destabilisation of the recording. Finally, change in input resistance (detected from a change in the bridge balance error; Figure 2G) was not permitted to exceed the balance error estimated offline at the beginning of the recorded data by more than two-fold. Violation of this rule of thumb indicated a deterioration in the recording quality potentially resulting from debris accumulating in the recording pipette, movement of the pipette tip or the recorded cell body, partial formation of a membrane ‘tube’ (similar to what would happen in the process of producing an outside-out patch by slowly retracting the pipette), or other reasons.

We obtained the seven longitudinal recording state/quality indicators described above for all our recordings. Multi-panelled graphs showing these indicators for each individual recording separately are provided in Supplementary Figures 1-13.

### Simulations

#### Simulating distributions of miniature excitatory postsynaptic potentials with a range of amplitudes and rise times

All simulations were carried out using our ‘minis’ software, developed in Matlab (Mathworks; RRID:SCR_001622), and performed on the Cardiff School of Biosciences’ Biocomputing Hub HPC/Cloud infrastructure. Simulation of minis-like events was based on an analytical solution to the passive cable equation for a transient current (Rall, 1977) using 100 lumped terms with a double exponential synaptic current with τ_1_= 0.15-3 ms (optimised in each instance/iteration) and τ_2_ = 2 ms (G. Major et al., 1993). The detailed description of how arbitrary shape PSPs were obtained is provided in our companion article (Dervinis and Major, 2024).

A pool of simulated events was generated combining four bivariate (two-dimensional: amplitude and 10-90% rise time) normal distributions. Six parameters described each distribution: amplitude mean μ_X_ and standard deviation σ_X_, 10-90% rise time mean μ_Y_ and standard deviation σ_Y_, a correlation (or rotation) factor ρ, and a (relative frequency or number) scale factor n:

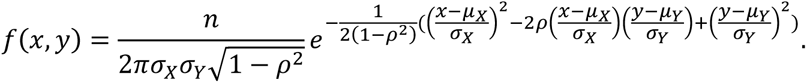

All distribution parameters were controlled by the GA.

#### Fitting distributions of detected real mini-like events

The Matlab ga() function was used to control 24 parameters describing Four (rotated) bivariate normal distributions (6 parameters per distribution). Additional controlled parameters were the synaptic rise time constant τ_1_ which ranged between 0.15 and 3 ms and the passive membrane time constant τ_m_. The range for the latter constant was taken to be between 0.9 times and 1.1 times the initial estimate based on the averaged waveform of all minis-like events detected in the ‘noise with real minis’ recording. Thus, there were 26 parameters in total, controlled by the GA.

The GA would start by generating multiple distributions of simulated minis or smEPSPs (i.e. ‘individuals’; we used 260 individuals per ‘generation’ with 10 individuals per parameter optimised by the GA; see a single run structure in Figure3A). Events would be drawn pseudo-randomly from a distribution and then simulated events of the chosen shapes (in terms of amplitude and 10-90% rise time) would be added to the ‘noise-alone’ recording at times that were also chosen pseudo-randomly (Figure 1). For generating the initial set of individuals (‘population’), the GA chose parameters pseudo-randomly within prescribed ranges. For each of the four bivariate normal distributions the parameter ranges were as follows: μ_X_ (-0.1-1 mV), σ_X_ (0.01-1 mV), μ_Y_ (-1-10 ms), σ_Y_ (0.25-10 ms), ρ (-1 to 1), and n (0-10,000 for the first distribution and –10,000 to 10,000 for the remaining three distributions; negative counts were used for ‘sculpting’ the base distribution). Any negative count bins within the resulting final distribution were zeroed out. Any events with negative rise times or amplitudes below the simulation threshold (varying non-negative value) were excluded.

Next, the GA assessed the ‘fitness’ of each individual in the generation by using the cost function (explained in the next subsection). The top 5% of the most fit individuals survived to the next generation without change, while individual ‘crossover’ (‘recombination’), genetic migration, and other GA parameters were set to default values of the Matlab ga() function. A new evolved generation of individuals was then created, which then served as the basis for yet another generation and so on. This process (a single run in Figure 3A) was repeated until an ‘individual’ was created with the lowest cost (highest fitness value) possible, or there was no improvement in the fitness for 100 consecutive generations, or the number of generations reached 1000 (Figure 3A).

**Figure 3:**
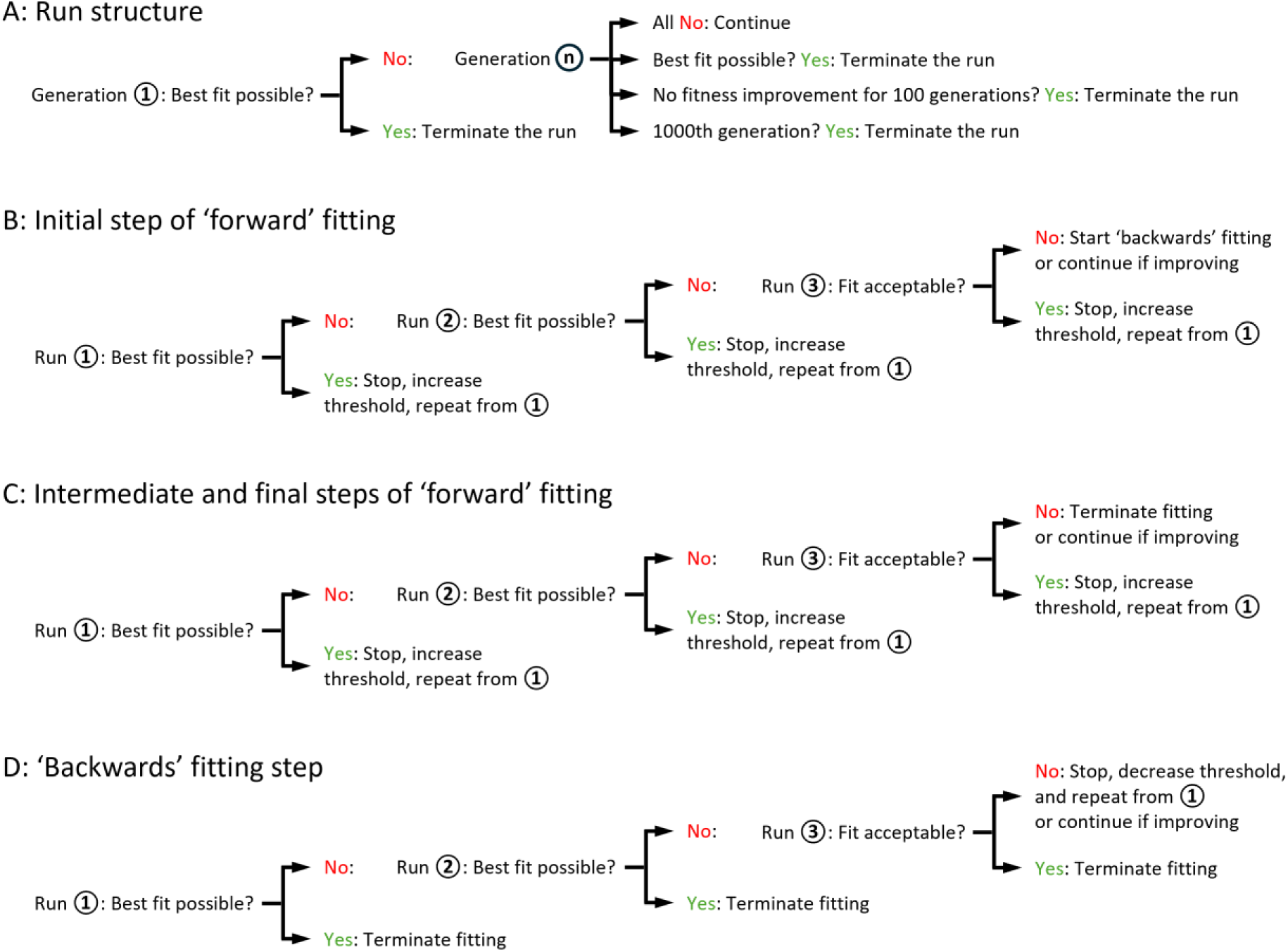
Schematic illustration of ‘minis’ genetic algorithm execution steps leading to the discovery of a minis source distribution underlying a distribution of minis-like events detected in a ‘noise with minis’ recording. (A) The structure of a single GA run. (B) The structure of the initial step of the ‘forward’ (‘upward’) fitting procedure where the lower threshold of the simulated minis amplitude is gradually raised. (C) The structure of intermediate and final steps of the ‘forward’ (‘upward’) fitting procedure. (D) The structure of the ‘backwards’ (‘downwards’) fitting procedure where the lower threshold of the simulated minis amplitude is gradually lowered.

**Figure 4:**
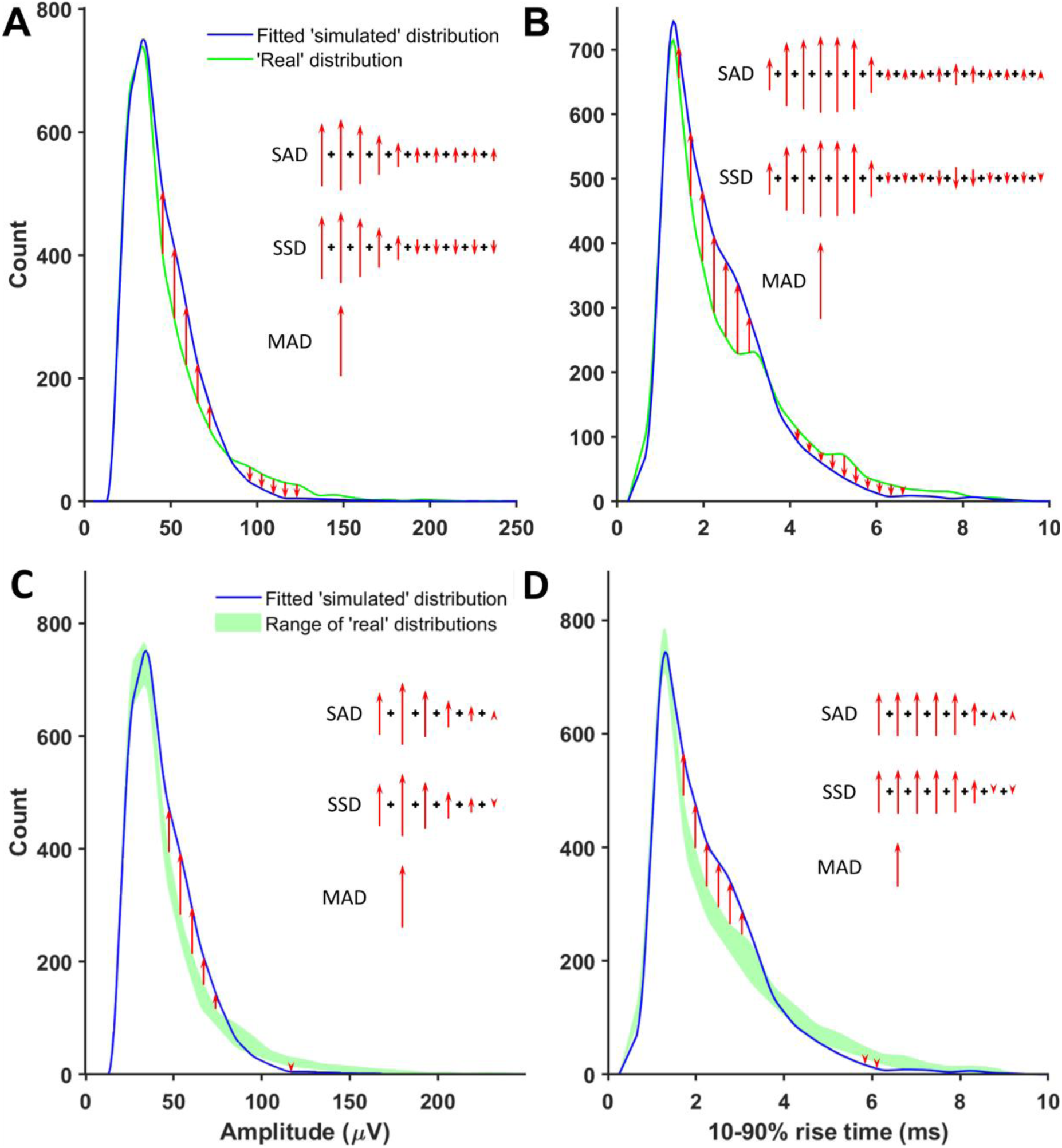
Non-shape components of the genetic algorithm’s cost function. (A) A schematic illustration of how amplitude SADs, MADs, and SSDs were calculated. Red arrows mark detected minis-like event count discrepancies between ‘simulated’ and ‘real’ ‘noise with minis’ distribution bins. (B) A schematic illustration of how 10-90% rise time SADs, MADs, and SSDs were calculated. (C) A schematic illustration of how amplitude ‘envelope deviation’ scores corresponding to SADs, MADs, and SSDs were calculated. Red arrows mark detected minis-like event count discrepancies between the ‘noise with simulated minis’ distribution and the envelope of all superimposed ‘noise with real minis’ distribution bins. (D) A schematic illustration of how 10-90% rise time ‘envelope deviation’ scores corresponding to SADs, MADs, and SSDs were calculated.

In cases where the GA terminated without converging on the highest fitness value, the distribution fitting procedure was reinitiated using the most fit individual as a seed (Runs 2 and 3 in Figure 3B). The population genetic diversity was reset by setting the GA-controlled parameter ranges to ±25% of the original ranges relative to the seed parameter values, to explore the space around the current best seed values. The re-running of distribution fitting was repeated until the GA converged to an ’apparently fittest possible individual’ (or the ‘best fit possible/achievable’; defined in the next subsection), or otherwise an ‘acceptable’ level of fitness was reached (also defined in the next subsection), or there was no improvement in fitness compared to the previous run for two consecutive runs while an ‘acceptable’ fit was not yet attained (Figures 3B and C). In the latter case, the entire distribution fitting procedure was reinitiated starting with a high simulated amplitude lower limit (e.g., 200 μV), with the limit being gradually lowered until the ‘best possible’ or ‘acceptable’ fit was attained (Figure 3D). This ‘backwards’ (‘downwards’) fitting procedure was required for several recordings having a large mean amplitude which, when we started with a low simulated amplitude lower limit, failed to converge to an acceptable fit.

If the most fit individual was within acceptable levels of fitness values, the lower threshold on smEPSP amplitude was increased by 10 μV (Figures 3B and C) (or lowered during the ‘backwards’ i.e. ‘downwards’ fitting procedure if no ‘acceptable’ fit was obtained; Figure 3D) and the entire fitting step was re-run. This was repeated until the GA stopped converging (or started converging during the ‘backwards’/’downwards’ fitting procedure) on a solution with acceptable fitness values.

The reason for attempting to push up the lower limit of the amplitudes of the smEPSPs is that it became obvious that GA-generated solutions were not unique. Given an acceptable fit, matching the histograms of events detected from the ‘noise with real minis’ recording can generally be achieved by using distributions with a larger number of smaller amplitude smEPSPs, which get ‘lost’ in the noise or summate with other smEPSPs, although the distribution of bigger smEPSPs is more tightly constrained. (Indeed, one can also often (but not always) achieve acceptable fits with *fewer* small amplitude smEPSPs (Supplementary Figure 14)). There is a large number of such distributions with ever-decreasing amplitudes of smEPSPs (in theory, all the way down to events resulting from brief single AMPA receptor (AMPAR) or NMDA receptor (NMDAR) channel sub-conductance state openings, injecting undetectably small amounts of charge – rare as these may be at ambient low background glutamate concentrations (Trussell and Fischbach, 1989) and resting membrane potential (Ascher and Nowak, 1988)). Therefore, the GA with the thresholding procedure described above has no guaranteed way of arriving at the *true* distribution of minis, only at a distribution giving an approximate *upper bound* on the *mean* (or median) mini amplitude – unless further biological constraints or assumptions are imposed (e.g. about background/extra-synaptic AMPAR or NMDAR channel activation or minimum numbers of glutamatergic ionotropic receptor channels per synapse). However, this should not be confused with the inability to ‘capture’ all (or most of) the postsynaptic charge, as is the case with the voltage clamp method of recording, applied to big, electrically distributed (non-isopotential) cells. This is rather an empirical observation of the fact that our method currently has a (poorly constrained) resolution limit on the lower end of the amplitude distribution of estimated minis, which may be further addressed (constrained) in future iterations.

#### Cost function

The goal of the minis distribution fitting procedure was to match ‘noise with simulated minis’ and ‘noise with real minis’ recordings as closely as possible, in terms of the distribution of amplitudes and 10-90% rise times of detected mini-like events in the two types of V_m_ recording traces (‘simulated’ vs. ‘real’ distributions), in the process ‘cancelling out’ any systematic errors in amplitudes or shapes introduced by the detection algorithm or the summation or superposition of events. This was achieved by minimising the cost function using the GA. The cost function was composed of four types of components: *shape constraints* on the *simulated* minis distribution, *sums* of *absolute* deviations (SADs) between ‘simulated’ and ‘real’ distributions, *sums* of *signed* deviations (SSDs) between ‘simulated’ and ‘real’ distributions, and *maximal absolute* deviations (MADs) between ‘simulated’ and ‘real’ distributions (Figures 4A and B). We avoided using the sum of squared errors (SSE) because it is sensitive to outliers (giving them undue weight).

We calculated the typical ranges of SADs and MADs for each of the 14 recordings by taking the individual ‘noise with real minis’ recording and dividing it into smaller files (range was 15-606 files), each the duration of a single recording sweep (20 s, typically). Each file in this set was subjected to the minis detection procedure resulting in a set of ‘real’ (detected mini-like events) distributions. These were then subtracted from each other, in all pairwise combinations, and the corresponding SAD (sum of absolute deviations) and MAD (maximum absolute deviation) scores were calculated for each of the possible n*(n-1)/2 file pairings (range 105 to 183315, corresponding to 15 to 606 available files, respectively). Six SAD- and six MAD-based scores were calculated based on: distributions of amplitudes of (1) *all* detected events, (2) the largest 50%, (3) the largest 10% and (4) the largest 2% of detected events, (5) distribution of all 10-90% rise times, and (6) a joint amplitude and 10-90% rise time distribution. Three different SSD (sum of signed deviations)-based scores were calculated based on the amplitudes of (1) the largest 50%, (2) the largest 10%, and (3) the largest 2% of detected events. These latter scores constrained high-amplitude (but low count) tails of ‘simulated’ distributions to have sufficient counts to avoid negative distribution deviations (‘simulated’ – ‘real’).

To ensure good fits to *all* portions of the histograms, and to prevent ‘dilution’ or ‘swamping’ effects on the high amplitude (but low event frequency) tail by the low amplitude, high frequency ‘hump’, the smallest ‘global’ percentile (same for every measure) was estimated, that guaranteed to include *half* of all single-sweep file pairings (real vs. real data) with *all* their 15 respective SAD, MAD, and SSD score values falling below this percentile threshold *simultaneously* (‘15-score combined 50^th^ centile’; Figure 5 and Tables 1 and 2). This centile estimate was used to determine cut-off values for all these 15 different scores, to construct the optimisation cost function. The ‘global’ centile value from single sweep file comparisons was then used with SAD and MAD scores from longer files during the GA- controlled fitting procedure (typically 5 recording sweep-long files to match the size of the ‘noise-alone’ recording files, 100 s typically).

**Figure 5:**
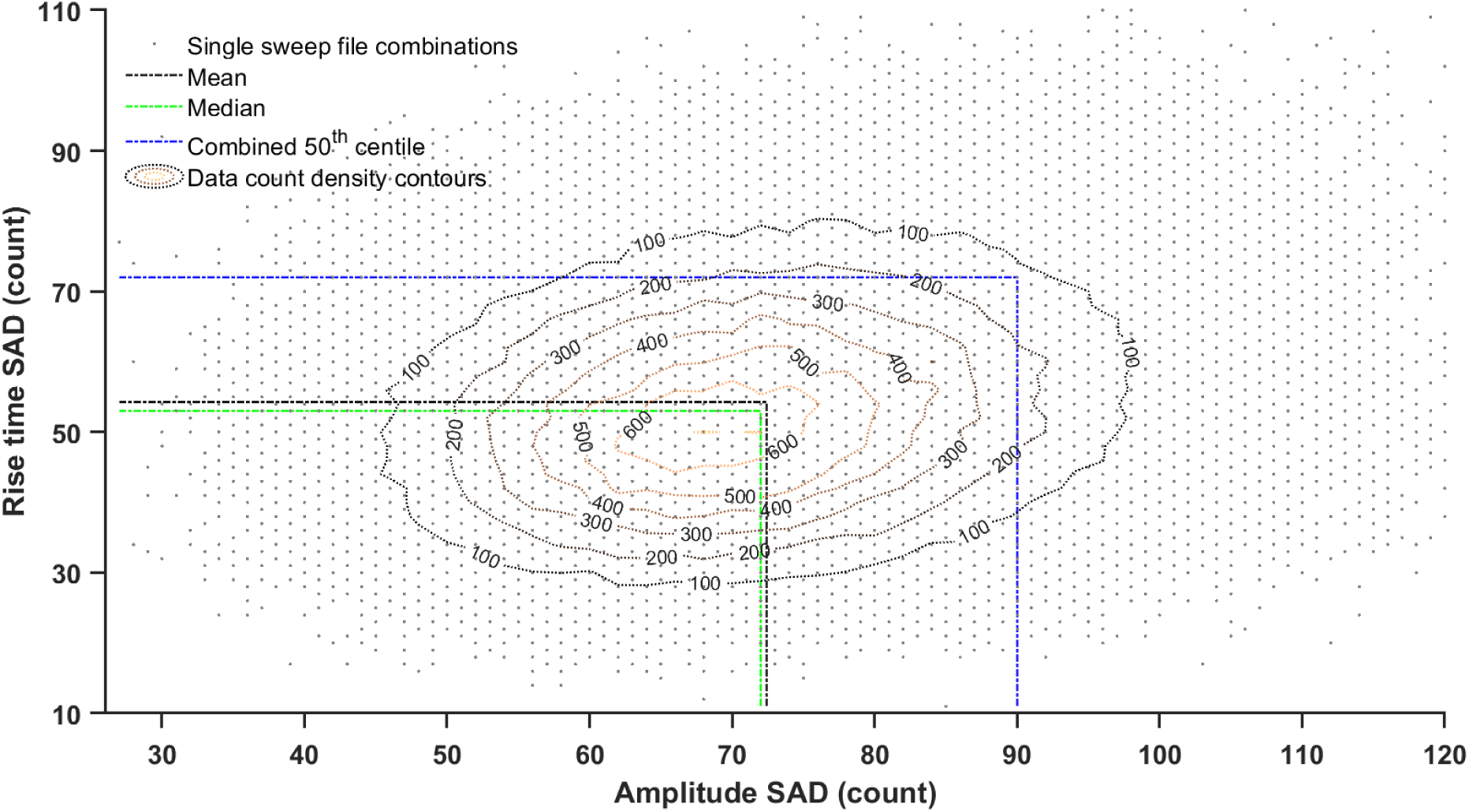
Amplitude and rise time scattergram (density plot) of sum of absolute deviation (SAD) scores between histograms from all possible pairs of ‘noise with minis’ single sweep files (4 s) from a layer 5 pyramidal neuron (p103a). The line marking ‘combined 50^th^ centile’ cutoff is based on 15 different SAD, SSD, and MAD scores as explained in Tables 1 and 2 (it extends to the remaining 13 dimensions not shown here). ‘15-score combined 50^th^ centile’ sets a lower (more exacting) threshold for accepting (‘steering’) distribution fits than the mean or the median would. This is justified because distribution fits need to satisfy all 15 SAD, SSD, and MAD scores *simultaneously* (e.g. to ensure that one of the tails of the distribution is not badly fit, as well as the remaining parts of the distribution).

**Table 1:** Failed hypothetical attempt to use 56^th^ centile as a basis for the combined ‘15-score 50^th^ centile’. The table shows a representative portion of a much bigger table illustrating a hypothetical scenario where the 15 SAD, SSD, and MAD scores are arranged in columns and rows corresponding to different two-file comparisons (many other comparisons are above and below those portrayed for the sake of the argument). SAD, SSD, and MAD scores are calculated by subtracting one file distribution from another (see the text for a detailed explanation). In this example, only *one* file comparison pair (green; 1/6 = 16.7%; row shaded in grey) satisfies the condition of having *all* its actual between-file SAD, SSD, and MAD values equal or below their respective 56^th^ centile (across all comparison pairs - a unique value applying all the way down each column) – i.e. *simultaneously* in *all* columns. Therefore, the 56^th^ percentile of each SAD, SSD, and MAD score would be too small to serve as the basis for the composite, joint or combined ‘15-score 50^th^ centile’: we need *half* the rows to be green, i.e. to pass the test.

| Amp<br>SAD<br>≤56<sup>th</sup><br>centile? | Top<br>50%<br>amp<br>SAD<br>≤56<sup>th</sup><br>centile? | Top<br>10%<br>amp<br>SAD<br>≤56<sup>th</sup><br>centile? | Top 2%<br>amp<br>SAD<br>≤56<sup>th</sup><br>centile? | Top<br>50%<br>amp<br>SSD<br>≤0? | Top<br>10%<br>amp<br>SSD<br>≤0? | Top 2%<br>amp<br>SSD<br>≤0? | RT SAD<br>≤56<sup>th</sup><br>centile? | Joint<br>SAD<br>≤56<sup>th</sup><br>centile? | Amp<br>MAD<br>≤56<sup>th</sup><br>centile? | Top<br>50%<br>amp<br>MAD<br>≤56<sup>th</sup><br>centile? | Top<br>10%<br>amp<br>MAD<br>≤56<sup>th</sup><br>centile? | Top 2%<br>amp<br>MAD<br>≤56<sup>th</sup><br>centile? | RT<br>MAD<br>≤56<sup>th</sup><br>centile? | Joint<br>MAD<br>≤56<sup>th</sup><br>centile? |
| --- | --- | --- | --- | --- | --- | --- | --- | --- | --- | --- | --- | --- | --- | --- |
| ... | ... | ... | ... | ... | ... | ... | ... | ... | ... | ... | ... | ... | ... | ... |
| Yes | Yes | Yes | Yes | No | No | No | Yes | Yes | Yes | Yes | Yes | Yes | Yes | Yes |
| Yes | Yes | Yes | Yes | Yes | Yes | Yes | Yes | Yes | Yes | Yes | Yes | Yes | Yes | Yes |
| Yes | Yes | Yes | Yes | No | No | No | Yes | Yes | Yes | Yes | Yes | Yes | Yes | Yes |
| No | No | No | No | Yes | No | No | Yes | Yes | Yes | No | No | No | Yes | Yes |
| No | Yes | No | No | No | Yes | Yes | Yes | Yes | Yes | Yes | No | No | Yes | No |
| Yes | Yes | Yes | Yes | No | No | No | No | Yes | Yes | Yes | Yes | Yes | No | Yes |
| ... | ... | ... | ... | ... | ... | ... | ... | ... | ... | ... | ... | ... | ... | ... |

**Table 2:** Using 90^th^ SAD centile as a basis for the combined ‘15-score 50^th^ centile’. The table shows another example where the 15 SAD, SSD, and MAD scores are arranged in columns and rows correspond to different two-file comparisons. SAD, SSD, and MAD scores are calculated by subtracting one file distribution from another (see the text for a detailed explanation). Now *three* of the six file pairings (*half* of them: green; rows shaded in grey) satisfy the condition of having *all* their between-file SAD, SSD, and MAD scores equal or below their respective 90^th^ centiles (across all comparison pairs - a unique number applying all the way down each column). Therefore, the 90^th^ percentile works as the basis for the combined ‘15-score 50^th^ centile’. Higher than 90 leads to more than half and lower than 90 leads to fewer than half of possible file pairs ‘passing’ all 15 tests simultaneously.

| Amp<br>SAD<br>≤90<sup>th</sup><br>centile? | Top<br>50%<br>amp<br>SAD<br>≤90<sup>th</sup><br>centile? | Top<br>10%<br>amp<br>SAD<br>≤90<sup>th</sup><br>centile? | Top 2%<br>amp<br>SAD<br>≤90<sup>th</sup><br>centile? | Top<br>50%<br>amp<br>SSD<br>≤0? | Top<br>10%<br>amp<br>SSD<br>≤0? | Top 2%<br>amp<br>SSD<br>≤0? | RT SAD<br>≤90<sup>th</sup><br>centile? | Joint<br>SAD<br>≤90<sup>th</sup><br>centile? | Amp<br>MAD<br>≤90<sup>th</sup><br>centile? | Top<br>50%<br>amp<br>MAD<br>≤90<sup>th</sup><br>centile? | Top<br>10%<br>amp<br>MAD<br>≤90<sup>th</sup><br>centile? | Top 2%<br>amp<br>MAD<br>≤90<sup>th</sup><br>centile? | RT<br>MAD<br>≤90<sup>th</sup><br>centile? | Joint<br>MAD<br>≤90<sup>th</sup><br>centile? |
| --- | --- | --- | --- | --- | --- | --- | --- | --- | --- | --- | --- | --- | --- | --- |
| ... | ... | ... | ... | ... | ... | ... | ... | ... | ... | ... | ... | ... | ... | ... |
| Yes | Yes | Yes | Yes | Yes | Yes | Yes | Yes | Yes | Yes | Yes | No | No | No | Yes |
| Yes | Yes | Yes | Yes | Yes | Yes | Yes | Yes | Yes | Yes | Yes | Yes | Yes | Yes | Yes |
| Yes | Yes | Yes | Yes | Yes | Yes | Yes | Yes | Yes | Yes | Yes | Yes | Yes | Yes | Yes |
| Yes | Yes | Yes | Yes | Yes | Yes | Yes | Yes | Yes | Yes | Yes | Yes | Yes | Yes | No |
| Yes | Yes | Yes | Yes | Yes | Yes | Yes | Yes | Yes | Yes | Yes | Yes | Yes | No | No |
| Yes | Yes | Yes | Yes | Yes | Yes | Yes | Yes | Yes | Yes | Yes | Yes | Yes | Yes | Yes |
| ... | ... | ... | ... | ... | ... | ... | ... | ... | ... | ... | ... | ... | ... | ... |

The cost function was constructed of 6 types of SAD scores and 6 types of MAD scores estimated using the ‘15-score combined 50^th^ centile’ procedure and 3 types of SSD scores. These 15 scores were used to evaluate how close the ‘simulated’ distribution was to the single selected ‘real’ distribution that was deemed to be of the best quality by a human observer (Figures 4A and B). Typically, it was a file close to the *end* of the ‘noise with minis’ recording epoch, therefore, being the most similar file, in terms of recording quality parameters, to the ‘noise-alone’ recording epoch (which was generally the *first* good-quality epoch safely *after* pharmacological blockade of neurotransmission was deemed complete). An additional corresponding 15 ‘envelope deviation’ scores (same as SADs, MADs, and SSDs but comparing to the envelope or lower and upper range limits of superimposed all available data distributions within that pharmacological epoch) were calculated assessing how much each ‘real’ or ‘simulated’ distribution deviated from the full set consisting of all the remaining ‘real’ distributions and the single ‘simulated’ distribution (Figures 4C and D). The ‘simulated’ distribution was expected to have lower deviation scores than the worst performing (most discrepant from the others) ‘real’ distribution (so the simulated distribution would not be picked as the ‘worst outlier’ by a double-blind observer).

‘Simulated’ distributions were further penalised if their underlying source ‘simulated minis’ distribution was non-unimodal, had a 10-90% rise time mean larger than 5 ms, or it violated a 10- 90% rise time skewness constraint that the mean 10-90% rise time of simulated minis should exceed the median at least by a quarter of the standard deviation, as seen for the real minis (with noise) distributions (range of 0.25 to 0.39 standard deviations). Finally, ‘simulated’ distributions were also penalised if their underlying ‘simulated minis’ amplitude distribution had a sharp cliff at its lower end. Hence, together with the four shape constraints there were 34 fitness components in total (15 + 15 + 4 = 34).

If all SAD, MAD, SSD, deviation scores, and shape constraints for comparing the ‘simulated’ distribution to the ‘real’ distributions were satisfied (≤ 15-score 50^th^ centile SAD and MAD cutoff values and ≥0 SSD values) at the end of the fitting procedure, the fit was deemed to be the ‘best possible’ (a ‘very good match’, by all criteria). Alternatively, the fit could also be deemed ’acceptable’ if all amplitude SAD, MAD, and SSD scores, as well as the distribution unimodality constraint were satisfied (excluding 10-90% rise time scores, all ‘envelope deviation’ scores, and the distribution skewness and ‘cliff’ constraints). The ‘best’ and ‘acceptable’ fits comprised the set of ‘good enough’ fits in terms of their GA-optimised ‘fitness’ which, however, did not guarantee a statistically acceptable fitness.

#### Final test for distribution acceptance or rejection

The notion of a ‘good enough’ fit is somewhat arbitrary, so far. Once the fitting procedure was completed, we set out to check whether the final fit was objectively acceptable using statistical tests. We divided the full available length of the ‘noise with real minis’ recording condition (epoch) into smaller files of the same length as the selected ‘noise-alone’ recording file and constructed a *set* of (independent) ‘real’ (‘noise with real minis’) distributions (histograms) by running the ‘minis’ detection algorithm on all these files (which we term the ‘real distribution set’). We then compared these individual ‘real’ distributions to each other, pairwise, by subtracting their marginal (‘1-D’) amplitude and 10-90% rise time distributions from each other, taking the absolute value of the differences, and summing over all bins (bin sizes of 10 µV and 0.5 ms; bin counts were normalised by the average bin count over all ‘real’ distributions to increase the weight of distribution tails). So we ended up with amplitude (Figure 6A) and 10-90% rise time (Figure 6B) sets of SAD scores for each distribution (individual file) within the ‘distribution set’ (the ‘real-real SAD score sets’; there is an amplitude and a rise time ‘real-real SAD score set’ for each distribution within the ‘distribution set’ corresponding to individual lines in Figures 6A and B). The number of SAD scores in each ‘real-real SAD score set’ corresponded to the number of possible comparisons between the distribution (file) of interest and the rest of the distributions (files) = n - 1 (given n different files). Finally, we selected the ‘worst’ (‘most discrepant’) and the ‘best’ (‘least discrepant’ compared to the others) ‘real’ reference distributions (files; the top and the bottom rows of SAD score matrices in Figures 6A and B, respectively), in terms of their median SAD scores (median over all SADs in the ‘real-real SAD score set’ for that distribution). Therefore, we were left with only two sets: The ‘worst real-real SAD set’ and ‘the best real-real SAD set’.

**Figure 6:**
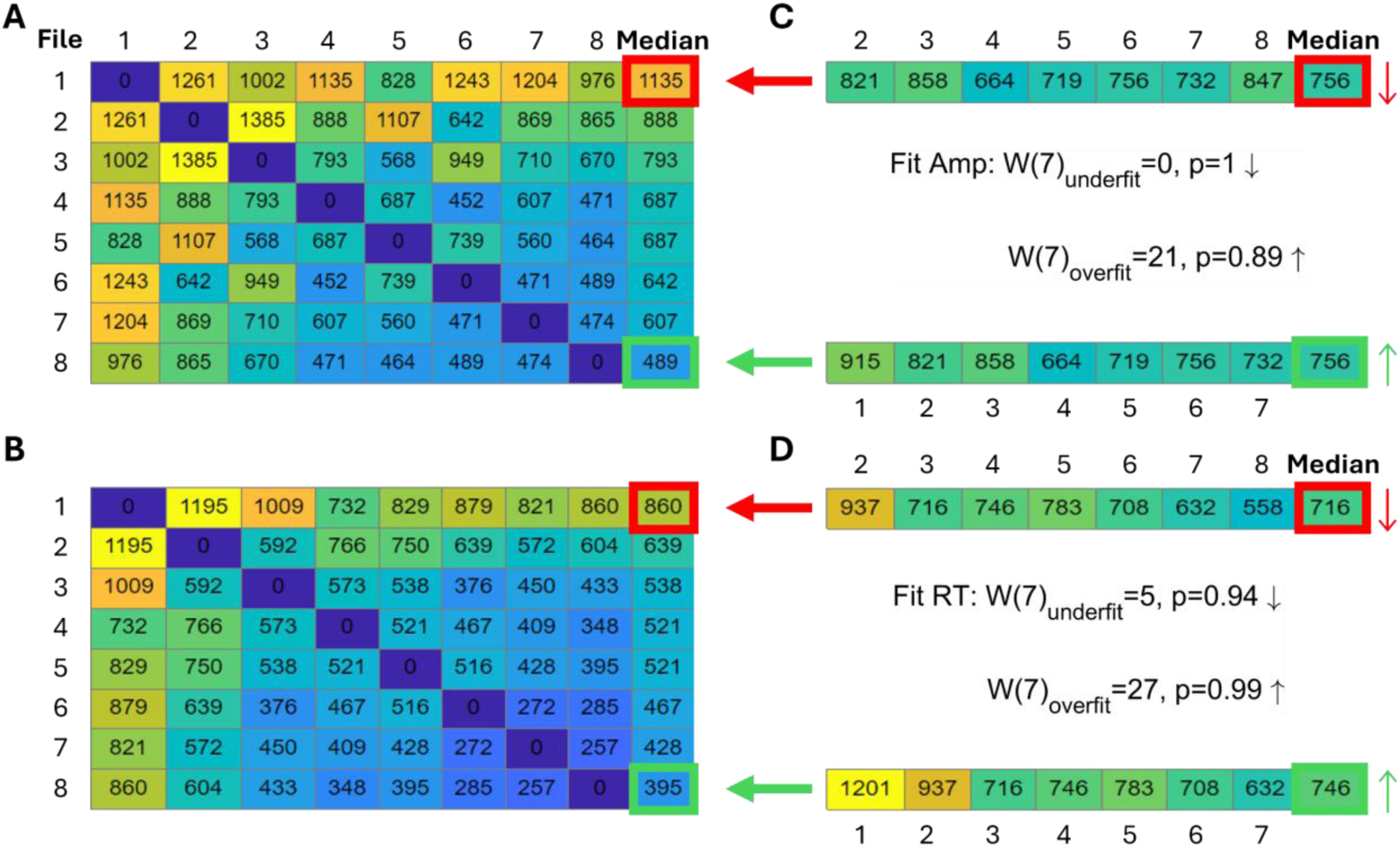
A schematic illustration of simulated minis distribution acceptance tests. (A) A matrix showing amplitude SADs for all ‘noise with real minis’ recording file comparisons. The last matrix column shows the median SAD score for all comparisons in individual matrix rows. The matrix cell with thick red borders indicates the ‘worst’-performing distribution, while the cell with thick green borders indicates the ‘best’-performing distribution. (B) A matrix showing 10-90% rise time SADs for all ‘noise with real minis’ recording file comparisons. (C) Two sets of SADs for comparisons of the ‘noise with simulated minis’ amplitude distribution and all non-reference ‘noise with real minis’ amplitude distributions. The top vector contains comparisons with the same ‘real’ distributions/files as the top matrix row in (A) as indicated by the thick red arrow (the reference ‘worst’ distribution was excluded from the comparison). The thin red arrow points in the downward direction consistent with the ‘simulated’ median (right) being lower than the ‘real’ ‘worst’-performing distribution’s median (left). The bottom vector contains comparisons with the same ‘real’ distributions/files as the bottom matrix row in (A) as indicated by the thick green arrow (the reference ‘best’ distribution was excluded from the comparison). The thin green arrow points in the upward direction consistent with the ‘simulated’ median (right) being higher than the ‘real’ ‘best’-performing distribution’s median (left). Results of the amplitude acceptance Wilcoxon signed-rank paired samples test for the simulated minis distribution are shown in the middle between the two SAD vectors. The top line tests whether the median of the SAD vector top set in (C) is larger than the median in the red-bordered cell of the matrix in (A) (W_underfit_; the number of files/distributions minus the reference distribution is indicated within brackets). The downward arrow indicates that the median of the SAD vector in (C) is smaller than the red-bordered median in (A). The bottom line tests whether the median of the SAD vector bottom set is smaller than the median in the green-bordered cell of the matrix in (A) (W_overfit_). The upward arrow indicates that the median of the SAD vector in (C) is larger than the green-bordered median in (A). (D) A set of SADs and acceptance tests as in (C) but for the 10-90% rise time distribution of simulated minis.

We also compared the ‘good enough so far’ (‘work-in-progress’) ‘noise with simulated minis’ distribution to all the non-reference ‘real’ (‘noise with *real* minis’) distributions. The comparison yielded another set of SAD scores: the ‘simulated-real SAD score set’ (Figures 6C and D). The median of this was then compared to the medians of the ‘worst real-real SAD score set’ and the ‘best real-real SAD score set’ using the non-parametric Wilcoxon signed-rank paired samples test. If the median score of the ‘simulated-real SAD score set’ was significantly larger than the median score of the ‘*worst* real-real SAD score set’, (i.e. if the distributions actually were identical, that discrepancy or more would only have occurred purely by chance less than 5% of the time; Bonferroni-corrected for a requirement to simultaneously satisfy both amplitude and 10-90% rise time tests), the ‘simulated’ distribution was deemed to be an inadequate fit and, was therefore rejected and the fitting procedure continued. (By contrast, if the median score of the ‘simulated-real SAD score set’ was significantly *smaller* than the median score of the ‘*best* real-real SAD score set’, the ‘simulated’ distribution was deemed likely to be an “excellent” fit; however, ‘passing’ this test was not strictly required and was used only for guidance). The ‘work-in-progress’ ‘good enough’ fit was ultimately accepted if the median of the ‘simulated-real SAD score set’ was *not* significantly larger than the median of the ‘worst (most discrepant) real-real SAD score set’. Intuitively, this corresponds (roughly) to a double-blind observer not being able to pick out that (simulated minis with real noise) distribution (histogram) from the (real minis with real noise) data distributions (histograms): it was not (obviously) the ‘most discrepant’ histogram.

### Statistical analyses

Parametric inferential statistics tests were used for incidence rates of minis (frequencies minis/s), mean amplitudes, and total cell membrane capacitances, as quantile-quantile (‘Q-Q’) plots indicated that these measures were normally distributed. Sums of absolute distribution bin deviations (SADs) were often not normally distributed as assessed by Q-Q plots, and therefore were tested using the corresponding non-parametric paired samples Wilcoxon signed-rank test.

### Data accessibility

All data analysed in this study are publicly available (Dervinis, 2024a, 2024b). The available data include ‘noise with minis’ and ‘noise-alone’ whole-cell patch membrane potential recordings that were used in computer simulations and minis’ distribution fitting, as well as GA distribution fitting results. All electrophysiological recordings were stored in Axon Binary File (ABF) format.

### Code accessibility

All analyses were carried out in Matlab and the analysis code is publicly available on GitHub (Dervinis, 2024c). The code is complete with instructions on how to reproduce figures reported in this study.

### Software accessibility

The present study reported the use of a novel quantal analysis method that is part of ‘minis’ software available on GitHub (Dervinis, 2024d).

## Results

### Estimating quantal size by direct subtraction of noise component

We have analysed ‘noise with minis’ recordings and ‘noise-alone’ recordings with the ‘minis’ detection algorithm. Detected mEPSP-like events had an amplitude and 10-90% rise time distributions that resembled log-normal or gamma distributions (Figure 7), left skewed with a long tail on the right. In the amplitude domain, the lower end of the distribution contains a mixture of real minis and noise fluctuations that were misidentified as real events (but also missing small minis cancelled out by noise, summating with larger minis, or failing to be detected due to being on a decay phase of a big preceding mini – the ‘overshadowing’ effect).

**Figure 7:**
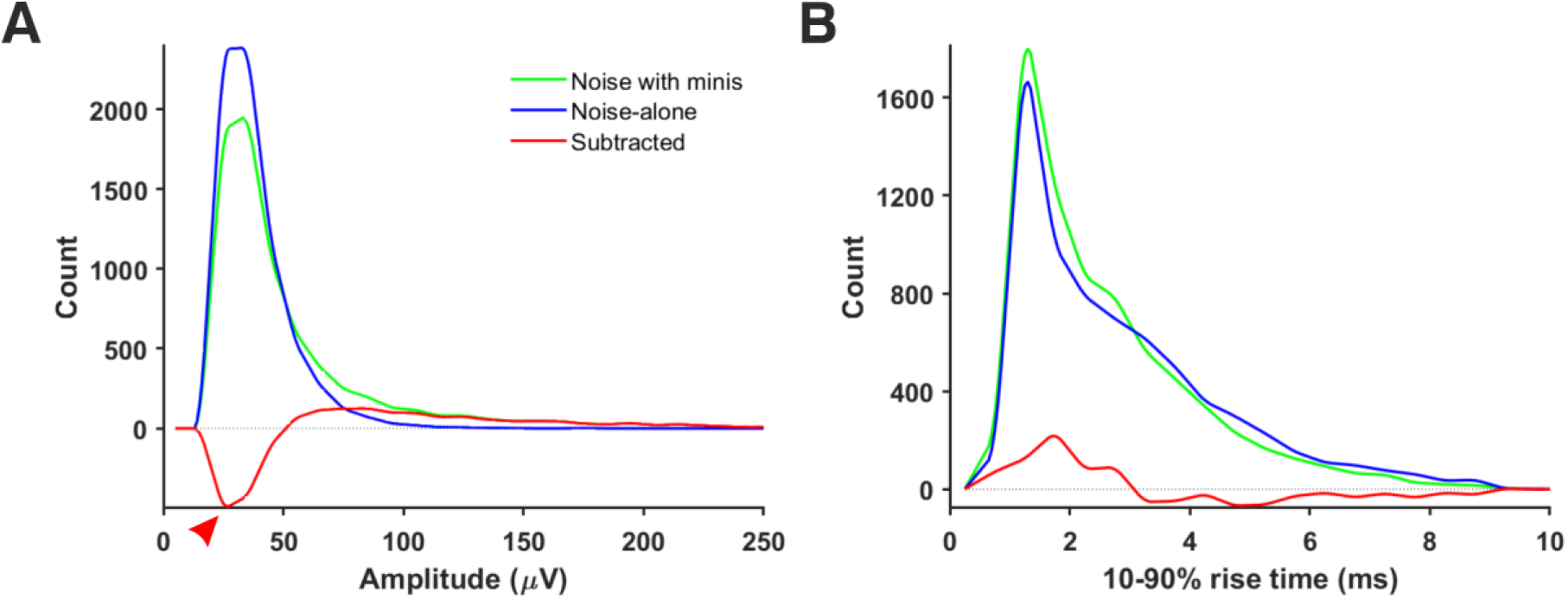
Amplitude and rise time distributions of detected mini-like events in ‘noise with minis’ and ’noise-alone’ recordings. (A) Amplitude distributions in a “thick” tufted neocortical layer 5 pyramidal neuron (p108b). Red arrowhead indicates negative bins after subtracting ‘noise-alone’ distribution from ‘noise with minis’ distribution. (B) 10-90% rise time distributions from the same neuron.

We initially tested whether the noise component (in ‘noise with minis’ recordings) could be reliably estimated based on the corresponding ‘noise-alone’ recording (where synaptic activity has been pharmacologically blocked), simply by directly subtracting the detected ‘noise-alone’ events distribution from the ‘noise with minis’ distribution, to yield an estimate of the pure minis’ component. Figure 7A shows both ‘noise with minis’ and ‘noise-alone’ detected event amplitude distributions superimposed. After subtracting the ‘noise-alone’ distribution from the ‘noise with minis’ distribution there appear negative low amplitude bins in the resulting distribution (indicated by an arrowhead in Figure 7A). These missing low amplitude events are caused by a greater number of small false positive mini-like detections in the noise-alone traces, and larger minis overshadowing smaller minis and small amplitude noise fluctuations in the ‘noise with minis’ recording (we have discussed this phenomenon in more detail in our companion article (Dervinis and Major, 2024)). Therefore, the direct subtraction approach can only provide a relatively crude ‘first order’, high-end upper limit on the mean of the mini amplitude distribution. Nevertheless, this over-estimate is more reliable than a highly subjective approach of simply estimating the mean amplitude based on events detected using a high amplitude threshold, or worse still, detected manually, by what could, perhaps uncharitably, be termed a “beauty contest” method (rather arduous, slow, subjective, and prone to human error/bias).

### Estimating quantal size by simulating excitatory postsynaptic potential source distributions

An alternative and more reliable way of estimating the quantal size in our recordings is to estimate the minis component directly using simulations. This relies on the ‘minis’ optimisation algorithm (GA) being able to closely match a ‘noise with simulated minis’ voltage trace (Figure 3) with the ‘noise with real minis’ recordings in terms of amplitude and 10-90% rise time distributions of minis- like events detected within these traces. We were able to successfully do that for all our recordings (see Figure 8 and Supplementary Figures 15-27). In the example shown in Figure 8, the overall amplitude SAD of the ‘simulated’ distribution was not significantly larger than the worst-performing ‘real’ distribution (unidirectional ‘underfit’ test: Is ‘simulated’ median SAD significantly > worst-performing ‘real’ median SAD? Wilcoxon signed-rank paired samples test statistic W (n = 16) = 105, p = 0.058 (Bonferroni-corrected due to comparing once on amplitudes and once on rise times); a high W statistic indicates a high absolute distance between the two medians, irrespective of its sign/direction, while p < 0.025 would indicate fit rejection, due to a fit SAD score being larger than the worst SAD score than would be expected by chance alone, at that significance level; downward arrow indicates ‘simulated’ median SAD < ‘real’ median SAD relationship in Figure 8) and was not outperforming the best ‘real’ distribution (‘simulated’ median SAD < best-performing ‘real’ median SAD Wilcoxon signed-rank paired samples unidirectional ‘overfit’ test W(16) = 135, p = 1; upward arrow indicates ‘simulated’ median SAD > ‘real’ median SAD relationship in Figure 8). Similarly, the ‘simulated’ 10-90% rise time distribution was also neither an underfit (‘simulated’ median SAD > worst-performing ‘real’ median SAD unidirectional ‘underfit’ test W(16) = 0, p = 1) nor an overfit (‘simulated’ median SAD < best-performing ‘real’ median SAD unidirectional ‘overfit’ test W(16) = 117, p = 1).

**Figure 8:**
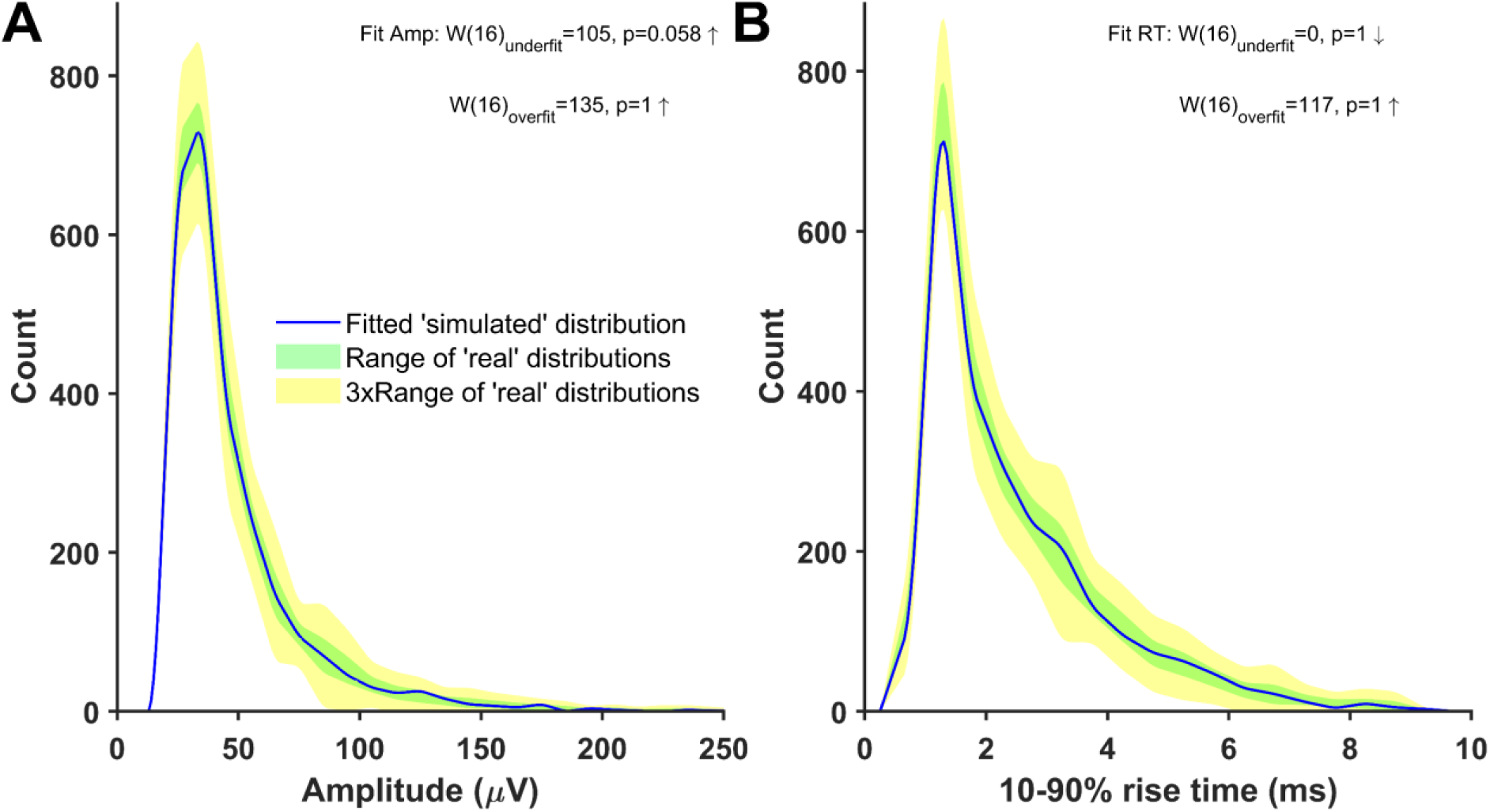
Distributions of mini-like events detected in a ’noise with simulated minis’ voltage trace reasonably closely matching ‘noise with real minis‘ recordings from the same cell, p106b (neocortical layer 5). Parameters of simulated minis were controlled by the Genetic Algorithm. Up arrows indicate ‘simulated’ SAD > ‘real’ SAD; down arrows ‘simulated’ SAD < ‘real’. The underfit test involves a comparison with the ‘worst-performing’ ‘real’ SAD (most discrepant from the others), whereas the overfit (‘excellent fit’) test involves a comparison with the ‘best-performing’ ‘real’ SAD (least discrepant from the others). With the underfit test (top), a fit SAD worse (bigger) than the ‘real’ SAD (up arrow), and p <0.05 would mean the fit is rejected (not the case here: both amplitude and rise time fits (A and B) were accepted). (See Methods and Figure 6 for more details). (A) Amplitude and (B) 10-90% rise time distributions (blue) of mini-like events detected in a ’noise with simulated minis’ voltage trace, compared with ‘envelopes’ or bands (shaded) of all available ‘real’ data distributions from the neuron (taking the most extreme low and high values at each point across all the available data files recorded with both APs and IPSPs blocked).

All simulations produced ‘good enough’ matches showing amplitude and 10-90% rise time distributions of detected minis-like events being within the range of (and thus indistinguishable from) amplitude and 10-90% rise time count values measured in sets of ‘real event’ distributions (Supplementary Figures 15-27). None of the fits performed significantly worse than the worst-performing ‘real’ distribution, qualifying them as ‘good enough’ (accepted) fits. All resulting minis source distributions are displayed in Figure 9 and Supplementary Figures 28-40. An example in Figure 9 shows that the ‘real minis’ source distribution likely has a log-normal or gamma-like shape in both amplitude and rise time domains.

**Figure 9:**
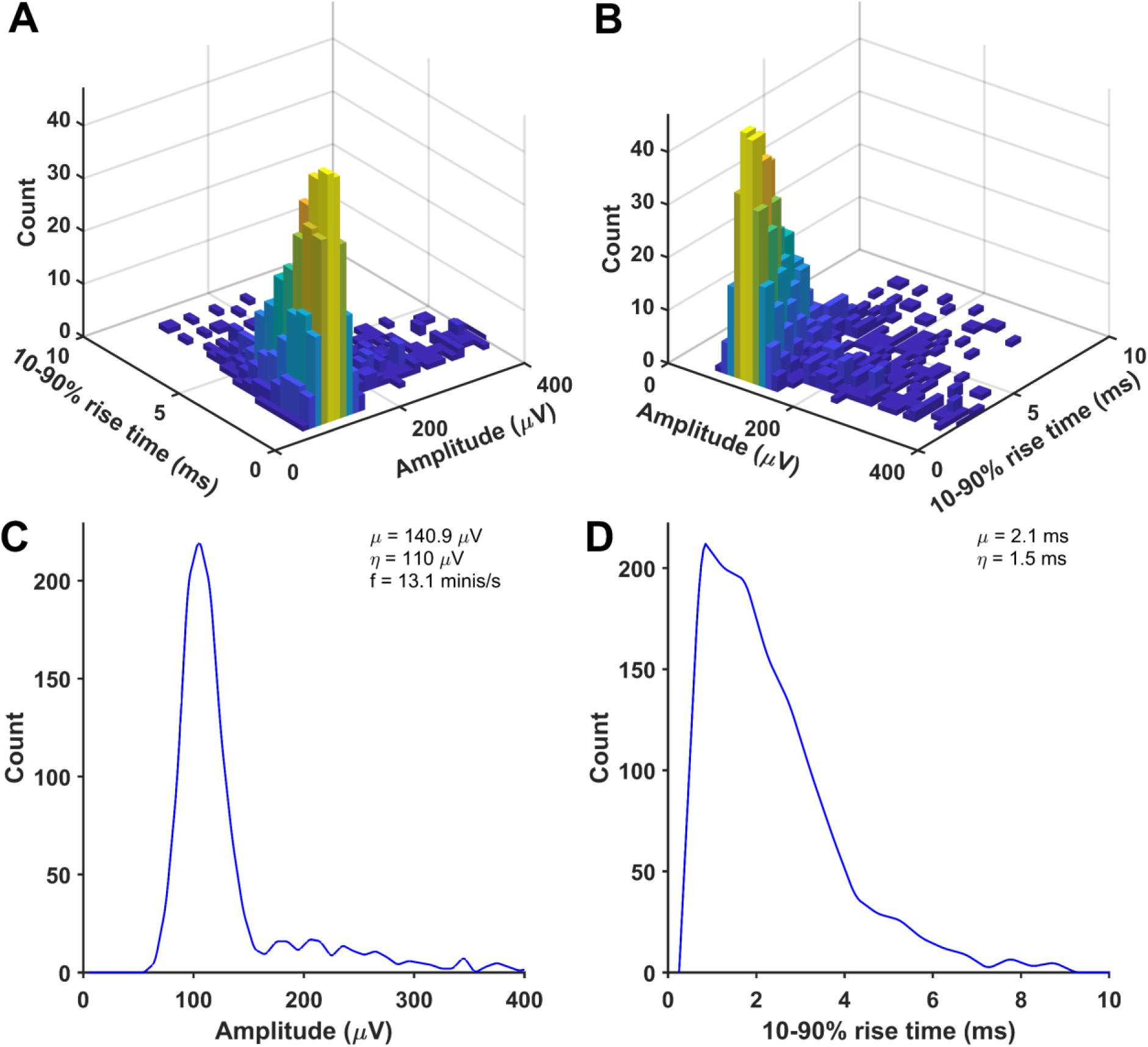
An example of a distribution of randomly selected simulated minis that was used to produce a close match between the distributions of detected mini-like events in a ’noise with simulated minis’ voltage trace (with a 60 µV lower limit on simulated amplitudes) and a ’noise with real minis’ recording, for a neocortical layer 2/3 cell (p131c). (A) and (B) Joint amplitude and 10-90% rise time distribution of simulated minis, viewed from different angles. (C) and (D) Corresponding ‘marginal’ (one-dimensional) amplitude and 10-90% rise time distributions, projected onto the axis indicated.

Incidence rates (frequencies of minis per second) estimated based on simulated minis source distributions ranged roughly between 10 and 100 minis/s (a mean of 27.3 ± 2.4 minis/s and a range of 6.8 to 58.1 minis/s; Figure 10). This estimate is an order of magnitude higher than the one based on the literature (3.6 minis/s; (Dervinis and Major, 2024)) and significantly higher than the one based on the direct subtraction method (15.9 ± 0.9 minis/s; t(13) = 3.51, p = 0.004) and also higher than the estimate based on amplitudes higher than a (deliberately chosen) unrealistically high detection threshold of 0.8 mV (14.3 ± 0.9 minis/s; t(13) = 3.69, p = 0.003). It is precisely within the range predicted in our companion article (Dervinis and Major, 2024). However, this range is possibly still somewhat of an underestimate, as mini source distributions were often generated using a lower limit on simulated amplitudes to constrain the distribution non-uniqueness, as discussed in the Methods subsection on the distribution fitting (also see Supplementary Figure 14).

**Figure 10:**
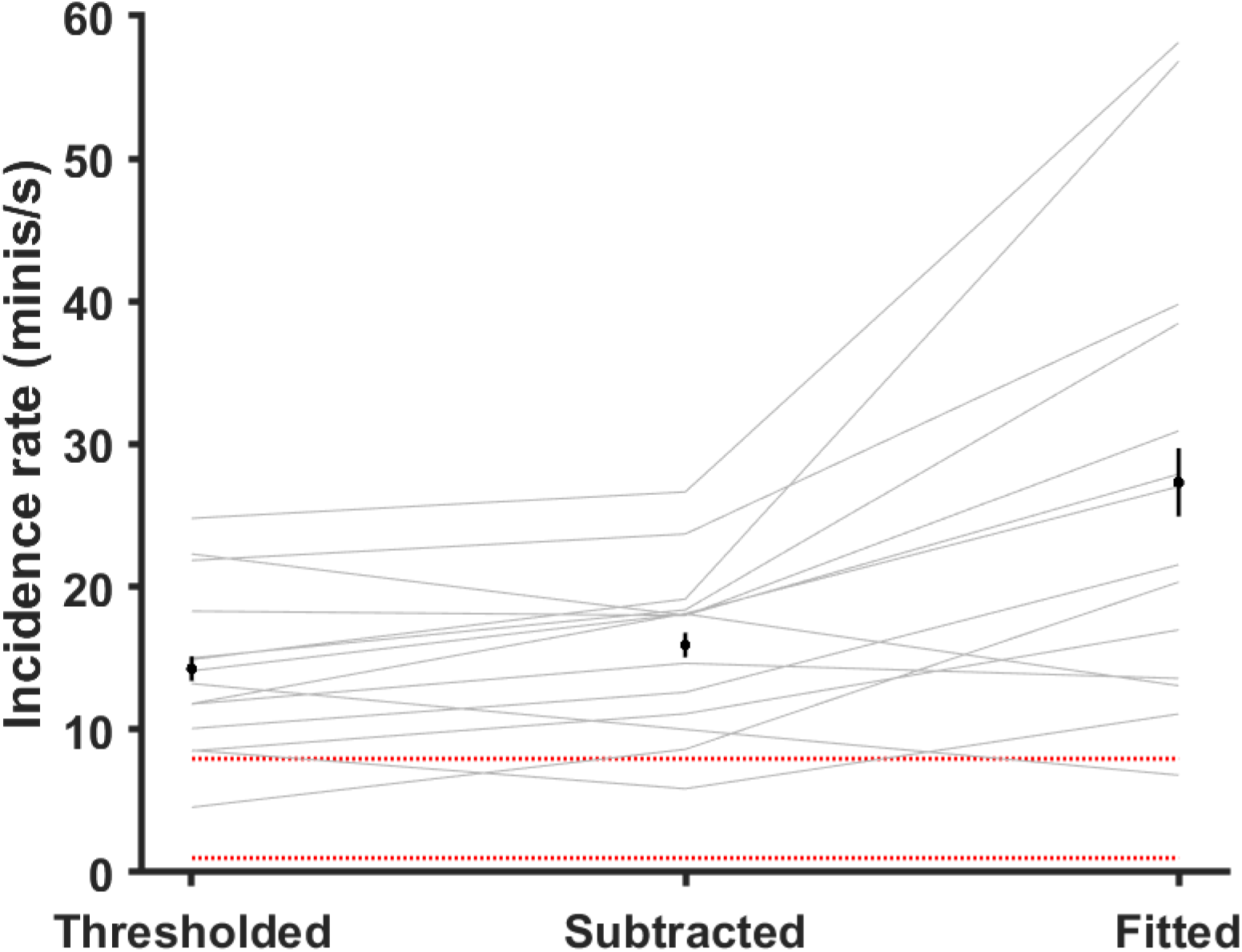
Estimated mEPSP incidence rates (‘frequencies’) using various methods (grey lines are individual cells). ‘Thresholded’ method: an ‘unreasonably high’ 80 µV detection threshold was applied to all 14 ‘noise with real minis’ recordings so only larger events were included. ‘Subtracted’ method: the distribution of mini-like events in the ‘noise-alone’ recording was subtracted from the distribution of mini-like events in the ‘noise with real minis’ recording and the incidence rate estimate was based on the positive bins of the resulting distribution. ‘Fitted’ method: the simulated minis distribution was used to estimate a lower bound on the mEPSP incidence rate (*lowest* acceptable rate statistically consistent with the real experimental data). Black markers and black vertical lines correspond to means and 95% confidence intervals. Red dotted lines mark the range of incidence rates typically reported in the literature; discussed in the companion article (Dervinis and Major, 2024).

Mean amplitudes of minis derived using simulations ranged between 53 and 182.9 µV (overall mean of 114.1 ± 5.5 μV, layer 2/3 mean of 133.1 ± 9.9 μV, and layer 5 mean of 95.1 ± 11.2 μV; Figure 11A). When compared to capacitance estimates (or 1/capacitance), there was a strong (very tight) correlation between each cell’s mean mEPSP amplitude and the inverse of its capacitance (which is proportional to cell size, assuming capacitance per unit area is more or less a biological constant for each cell class/type of membrane/density of channels with charged gates moving within the membrane (Gentet et al., 2000); r = 0.96, p = 7.1x10^-8^, R^2^ = 0.92). That is, the larger the cell, the smaller the amplitude of its somatically detected minis and vice-versa. This relationship is predicted by the theory of capacitors (C = Q/V, where C stands for capacitance, Q stands for charge, and V stands for voltage across the cell’s membrane), and it is a direct demonstration of an attenuating effect of cell’s capacitance on the amplitude of somatically detected minis. If a direct line was drawn through the data and the origin of the axes in Figure 11A, the slope of this line would provide a quantal size estimate for excitatory synapses in the CNS in terms of ‘injected’ charge (Q = ΔV/Δ(1/C)) which is approximately equal to 31.3 fC, for these data. The fact that the least square fitted line to the data (y = 30.6x + 2.76), with a negligibly small intercept term (2.76), roughly coincided with the line through the origin fitted to the data (y = 31.3x) indicated that no obvious ‘proactive’ mechanism *varying with cell size* was in place to compensate for the attenuation of the average minis’ amplitude due to the size of the cell.

**Figure 11:**
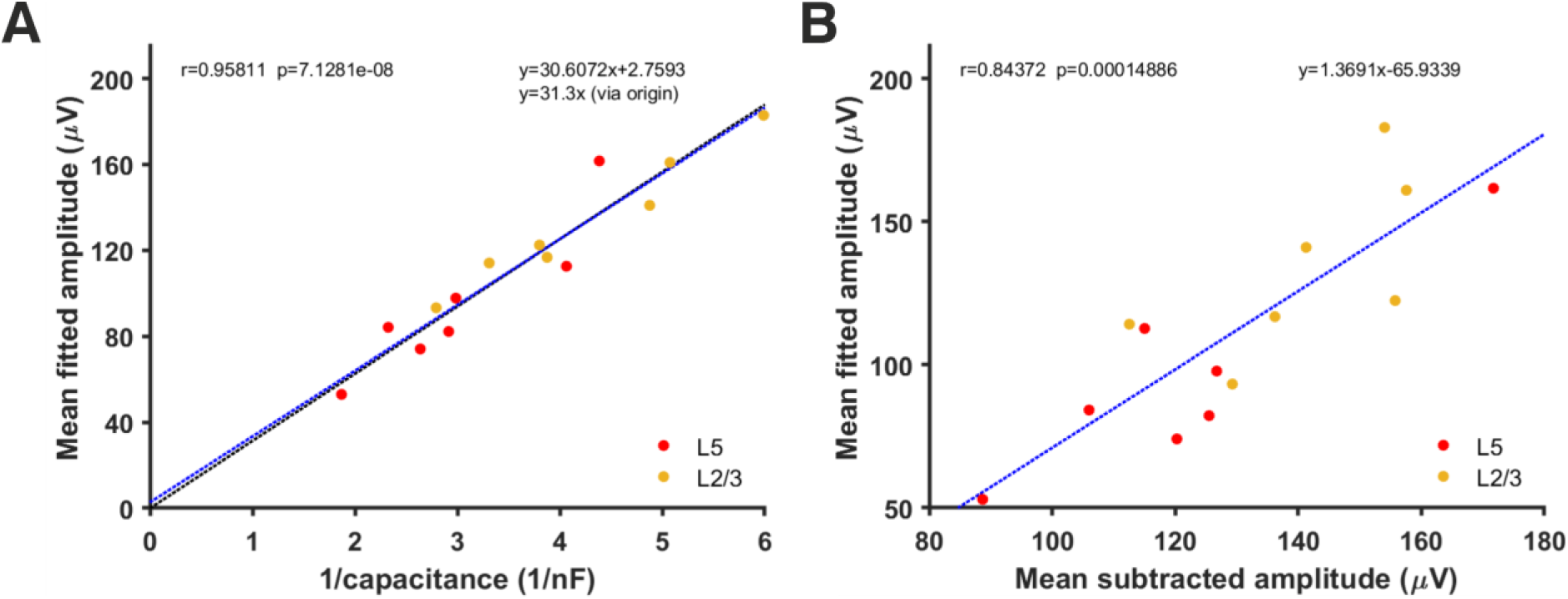
Relationship between each cell’s inverse total membrane capacitance and its estimated mean mEPSP amplitude using simulations and fitting of minis distributions. (A) Correlation between mean mini amplitudes from different neurons, as derived using the simulated minis distribution fitting method and the inverse of each cell’s total capacitance. The blue line is the least square fit to the data while the (nearly indistinguishable) black line is the fit to the data that passes through the origin. (B) Correlation between the mean mEPSP amplitude of each neuron estimated from ‘fitting’ versus ‘subtracting’ (the corresponding noise histogram, as in Figure 7).

Another notable observation was that the mean amplitude estimates derived either by noise subtraction or by minis source distribution simulation were highly correlated (r = 0.84, p = 1.49x10^-4^, R^2^ = 0.71 Figure 11B). The equation of the line (y = 1.37x – 65.9) fitted to this data provides a possible “shortcut” method to approximate the mean amplitude of fitted minis’ source distribution by inserting the mean estimated using ‘naïve’ noise subtraction directly into this equation.

### Generic miniature excitatory postsynaptic potential distribution in neocortex

To construct a generic mEPSP distribution, we normalised amplitudes of simulated minis’ source distributions shown in Figure 9 and Supplementary Figures 28-40 by dividing them by their respective means, taking the total average, and then scaling the amplitudes to the overall mean (114.1 ± 5.5 μV). The resulting combined (amplitude and 10-90% rise time) distribution is shown in Figures 12A and B. The corresponding marginal amplitude and 10-90% rise time distributions are shown in Figures 12C and D, respectively. Both marginal distributions have left-skewed log-normal-like or gamma-like distributions. This may be useful in neural network simulations, or to potentially speed up (and constrain) fitting of simulated minis on noise to real minis with noise, using the GA (e.g., limit the source distribution to one 2-D ‘rotated’ log-normal or gamma distribution).

**Figure 12:**
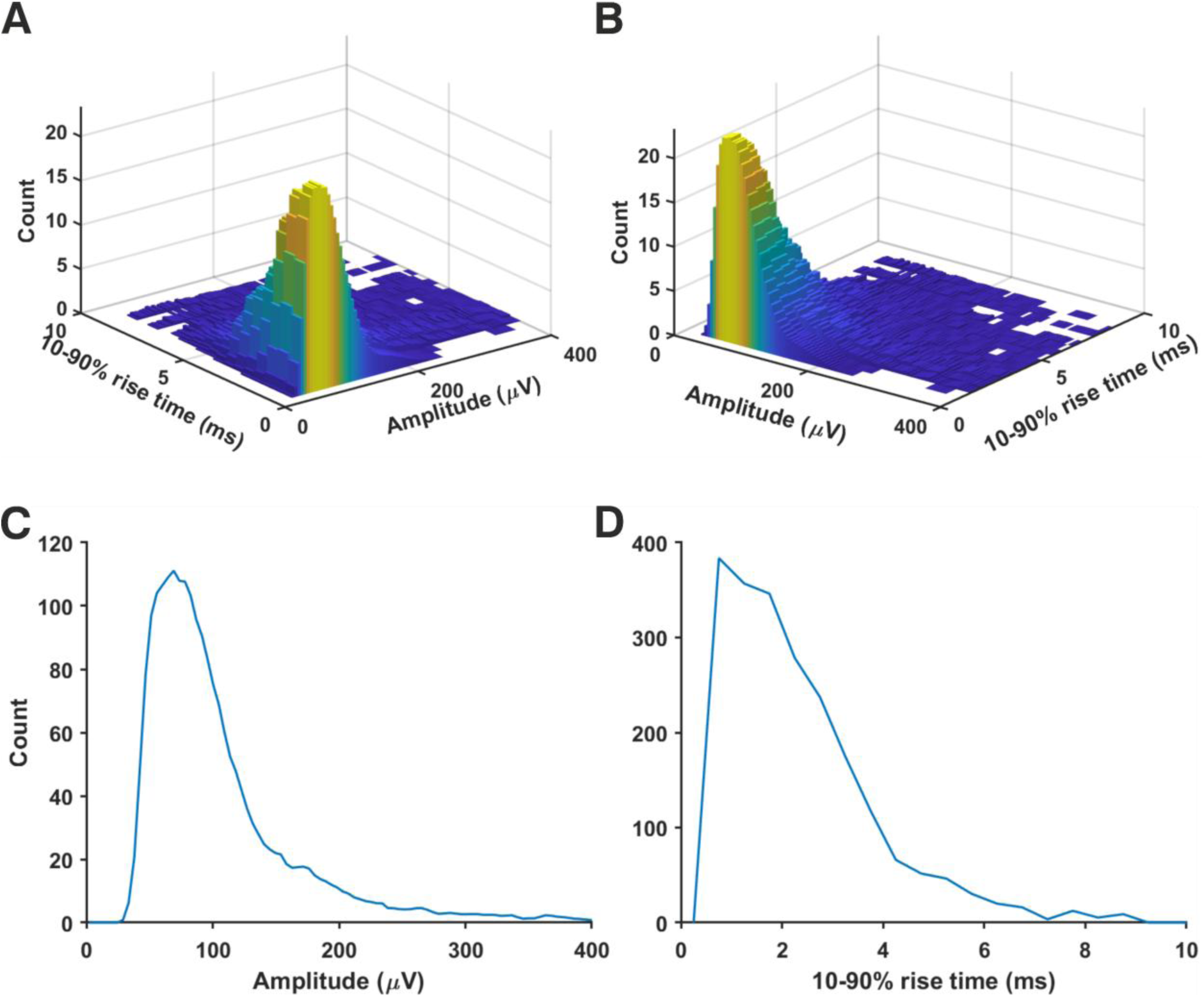
Generic ‘averaged’ mEPSP distribution from rat (somatosensory) neocortical pyramidal neurons. Distributions resulting from all 14 recordings were averaged. All mini amplitudes were first normalised by the means of their respective source distributions, then multiplied by the average of the average amplitudes (114.1 ± 5.5 μV) across all the neurons analysed. (A) A joint amplitude and 10-90% rise time distribution of simulated minis. (B) The same distribution but viewed from a different angle. (C) ‘Marginal’ (1-dimensional) amplitude distribution projected from the above distribution onto the amplitude axis. (D) ‘Marginal’ 10-90% rise time distribution.

## Discussion

We report two major scientific advances. First, a novel quantal analysis method based on sequential somatic recording, from a given neuron, of a) mEPSPs (after blockade of action potential-mediated neurotransmission, and all inhibitory neurotransmission, including mIPSPs) on a background of other physiological noise, and b) subsequent recordings from the same cell of noise alone, after additional pharmacological blockade of mEPSPs (all glutamatergic ionotropic excitatory neurotransmission). The latter type of recording provides a means to estimate the properties of the noise component which can then be used to take into account noise fluctuations in the combined minis and noise recording. However, this was done not by a direct subtraction of the histogram of amplitudes (and/or rise times) detected from the noise-alone recordings, but rather by the simulation of the underlying minis component and repeatedly superimposing (at random) the waveforms of simulated minis onto the noise trace, varying source distribution parameters until the histograms detected from the ‘noise with simulated minis’ voltage trace closely matched those detected from the ‘noise with real minis’ recording. The key tool for achieving this match was a genetic algorithm that optimised the parameters of the 2-dimensional simulated minis distribution in terms of amplitudes and 10-90% rise times (and rate, i.e. minis per second), simultaneously, to best fit the experimental data distributions.

The second major advance was the application of our novel quantal analysis method, that successfully measured the mean ‘electric charge injected’ size of excitatory synapses of cortical pyramidal cells, which was shown to be approximately 31.3 fC, across the entire sample of neocortical pyramidal neurons recorded. This number was inferred from the slope of the line which tightly fitted mean mEPSP amplitude vs. 1/total cell capacitance, in accordance with the theory of capacitors. The fact that this line had an intercept at the origin of the axes indicated that there was no biological compensation (other than *number* of synapses) for the amplitude attenuation of minis due to cell size (i.e., total capacitance). Indeed, cell capacitance played a major role affecting the *voltage* amplitude of mEPSPs, resulting in a large proportion of mEPSPs being otherwise indistinguishable from noise (as independently found in numerous other studies, e.g. (Major et al., 2013; Nevian et al., 2007), summarised in (Major et al., 2013)). The number of these ‘missed’ minis was so high that the estimated minis’ incidence rate (frequency) was in the order of 10-100 minis/s: an order of magnitude higher than typically reported in the synaptic physiology literature (with the notable exception of (Nevian et al., 2007), which predicted this result, by simultaneous dual somatic and basal dendritic whole-cell patch recordings of mEPSPs, with dendritic detection, and simultaneous somatic measurement). Finally, both the amplitude and 10-90% rise time distributions of minis had a left-skewed log-normal- or gamma-like shape.

These findings have some major implications. First, the method, as used for estimating the mEPSP distributions in cortical layers 2/3 and 5 of cells of various size, and then using these distributions to arrive at the electrical size of excitatory synapses, serves as a proof of principle that it can be successfully used to estimate PSP distributions of various types in the CNS. The next logical step of this method is to extend its application to the estimation of amplitude and rise time distributions of mIPSPs generated by different types of inhibitory synapses and, as a result, their ‘electrical sizes’. Moreover, the ability to obtain relatively accurate ‘quantal’ size estimates unobscured by physiological noise is important for constraining synaptic parameters in physiologically realistic simulations of neural activity. For example, now we should be able to better estimate how many active synapses are needed to generate an NMDA spike within a single dendritic segment (Major et al., 2013). The importance of these number estimates is not restricted to the domain of physiologically realistic simulations but also extends to more abstract computations that use NMDA spike-like mechanisms as coincidence detectors in the latest cortically inspired/constrained models of artificial intelligence (Hawkins et al., 2019). Accurate estimates of synaptic transmission parameters thus have very wide-reaching ramifications.

These findings also draw attention to some possible methodological issues in the field of synaptic physiology. As discussed in our companion article (Dervinis and Major, 2024), the reported incidence rates (‘frequencies’) for mEPSCs are generally well below 10 minis/s. Even if one doubts the incidence rates we arrived with simulations, in the range of 10-100 minis/s, the rates given by ‘naïve’ direct subtraction of the distribution of events detected in the ‘noise-alone’ recording from the distribution based on the ‘noise with minis’ recording are still within 5.9 to 26.7 minis/s, which we know to be an underestimate. Therefore, reported mini incidence rates below 10 minis/s are almost certainly underestimates. This may have multiple causes, including lower slice temperatures in the majority of studies (we used physiological, ca. 37 °C), and a significant temperature dependence of synaptic release, as well as differences in slice health and slicing angle (which can affect branch and local circuit ‘amputations’). Another big cause could be the (inappropriate) use of voltage clamp, especially if the R_ser_ is not compensated fully, as this would shrink and low-pass filter mPSCs, obscuring the smaller ones. As discussed earlier, these estimates could be improved by switching to voltage *recording* instead of using voltage clamp (Major, 1993; Rall and Segev, 1985; Spruston et al., 1993; Williams and Mitchell, 2008), but at the price of somewhat slower decay times of minis (Major, 1993), thus more temporal overlap/integration and consequent difficulties detecting peaks. But more importantly, a strong association between cell capacitance (i.e. surface area) and the extent to which somatically recorded minis’ amplitudes are attenuated, indicates that special care must be taken to account for the effect of background noise fluctuations on the amplitude measurements - as we have done here. This association is yet more evidence supporting experimental observations of massive mini amplitude attenuation as the PSP spreads from its dendritic source location to the soma, and beyond (Larkum et al., 2009; Major et al., 2013; Nevian et al., 2007; Stuart and Spruston, 1998; Williams and Mitchell, 2008; Williams and Stuart, 2002), especially in cells with large dendritic (or axonal) arbours, hence large surface areas and overall membrane capacitance. The apparent conservation of average synaptic charge may correspond to the observation that dendritic spines and post-synaptic densities (the specialised regions opposite release sites, containing high densities of synaptic receptors, with a substantial fraction tethered to the cytoskeleton) are not grossly different in size between different classes of pyramidal neurons in the neocortex, although differences exist between cortical layers, cortical areas, and, particularly, species (Yuste, 2010).

It is also worth noting that understanding the link between cell capacitance and somatically recorded average mini amplitudes, as well as knowing actual shapes of mini distributions in terms of their amplitudes and rise times, allows us to generate artificial data where the ‘ground truth’ is known. Such labelled data could perhaps be used to train artificial neural networks (ANNs) which could then be used to develop ‘next generation’ quantal analysis methods allowing detection of minis directly in mixed ‘noise with minis’ recordings. We are not the first to suggest the application of ANNs to this issue (see (Pircher et al., 2022) and (Wang et al., 2024)). However, we are not aware of attempts to use realistic mini distributions for this purpose.

Even though the novel ‘quantal analysis’ method we present here is an important advance, it nevertheless has limitations. One of them is the need for (relatively) long duration recordings using different pharmacological agents at various stages. However, the somatic recording set up is not as complicated and high-skill as an alternative multisite 2-photon targeted dendritic recording set up would be (Nevian et al., 2007), a heroic technique which has not to our knowledge been fully reproduced in other labs. Experiments themselves could be further streamlined, as we did in some cases, by pre-blocking action potentials before obtaining the whole-cell recordings and using high solution perfusion rates. This is particularly useful for smaller cells which are more prone to osmotic overloading or shrinkage and resulting recording instability after a few tens of minutes (the pipette solution cannot exactly reproduce the cytoplasm, as it contains no proteins or other macromolecules, and is missing many common metabolites). Moreover, the distribution fitting procedure involving the GA can be lengthy and computationally intensive. However, the ‘minis’ software is parallelised at the PSP detection level, as well as at the GA optimisation level, making it particularly suitable to be executed on multi-core computing clusters. Although our ‘minis’ software combined with two-stage recordings offers unprecedented control for the background physiological noise, provided recording stability is very good throughout, we understand that many researchers may not be persuaded to adopt it themselves. The possibility of misleading fits if the recording noise became substantially worse between ’noise with minis’ and ‘noise-alone’ recording phases needs to be vigilantly excluded. Nevertheless, our ‘minis’ software should still be useful for detecting mini-like events without recording independent noise, or fitting simulated to real distributions, if there is some other credible cut-off or separation method that can be used to distinguish between small or partially cancelled minis and mini-like noise. We anticipate, indeed, that, realistically, this scenario is likely to be its most popular use.

Some researchers may be critical of a method that does not simply rely on a direct measurement of minis but rather on an approximation via indirect means, using simulations. This is a valid criticism that could (perhaps) be addressed in the future by using a method of repeatedly subjecting a voltage recording to the minis detection procedure with an increasingly lower threshold while subtracting detected events from the voltage recording, or perhaps compensating for the ‘exponential-like’ decay phases of recent, temporally integrated, detected mini-like events. Some combination of these approaches could potentially recover most of the smaller minis that are ‘overshadowed’ by the larger ones (the so called ‘negative bins’ problem illustrated in Figure 7A), providing the waveforms were reasonably stereotyped in shape (not the case under voltage clamp, in neurons with large or extended or highly-branched dendritic arbours (Major, 1993; Jonas et al., 1993).

Furthermore, other researchers may question the ability to pharmacologically isolate the background physiological noise. Our ‘noise with minis’ recordings may contain random openings of individual AMPAR or NMDAR channels (possibly extra-synaptic), although background glutamate levels are likely too low to significantly open AMPAR channels (Trussell and Fischbach, 1989), and NMDAR channels would mostly be closed by Mg^2+^ block at resting membrane potential (Ascher and Nowak, 1988); it is also possible (in principle), but unlikely in practice, that our ‘noise-alone’ recordings may have failed to fully eliminate all AMPAR and NMDAR channel openings in the presence of quantal release of neurotransmitter (we used high AMPAR and NMDAR blocker doses, well above IC50s’ (Lehmann et al., 1987; Randle et al., 1992) and rapid slice perfusion, to achieve quick wash-in times, and checked for stationarity of the blockade by comparing multiple measures from consecutive traces: see Supplementary Figures 1-13). Background/physiological/electrical noise itself may also be drifting between two such (hypothetical) conditions, with the presence or absence of mEPSP pharmacological blockers. Occasionally, we rejected recordings (or epochs) with suspicious or obvious instabilities. These factors are unlikely to have affected the results significantly, due to the small amplitudes of any remaining membrane potential variations from these potential (but unlikely) causes in the fitted cells. Two largely independent analyses of the same data by the two authors separately yielded similar results and conclusions (albeit with somewhat different data selection, sweep lengths, and tests of goodness of fits: the ‘first pass’, not presented here, relied more on ‘double-blind’ visual inspection of multiple real minis with noise data distributions from the same cell superimposed on the simulated minis with noise fit distribution: if the latter ‘stood out obviously from the crowd’, it was rejected).

Finally, researchers may be reluctant to switch from the more commonly used voltage clamp recording method to unclamped voltage recording and, thus, would question whether our method captures postsynaptic currents in their full totality without distortions. We accept that there are experimental settings where the use of the voltage clamp may be preferred, like recording from small electrically compact granule cells in the cerebellum. ‘minis’ is not limited only to applications involving postsynaptic potentials and can equally well be used in detecting and analysing postsynaptic *currents* (indeed we have done so for several of the cells reported here, unpublished observations). However, as we argued throughout this article, the benefits of unclamped voltage recording in large cells are striking. Unlike voltage clamp, voltage recording offers a near-full capture of postsynaptic charge in these cells (when estimated from the PSP peak amplitude). Current leak through the membrane is relatively low, to first order, during the fast EPSP rising phase (i.e. during rapid axial charge redistribution; Major et al. 1993a), and recording of ‘noise with minis’ and ‘noise-alone’ combined with optimised simulations offers an unprecedented ability to estimate the actual minis population distribution and the upper limit on their average amplitudes. When care is taken to block other synaptic currents (as we did), then (in these neurons, at least) any boosting/sculpting/shunting impact of other active membrane channels can be ignored, reasonably safely, to first order, at the resting V_m_, as somatic (and dendritic) pulse responses scale relatively linearly within ca. 10 mV transients in this V_m_ range (Nevian et al., 2007). Voltage *recording* is also more resilient to small changes in R_ser_ and the compensation thereof, which under voltage *clamp* could substantially affect noise. Under voltage clamp, the noise recorded during the minis epoch may be different than during the noise-alone epoch, if (say) R_ser_ were to creep up a little. In part, this is because the clamp currents are directly proportional to compensated series conductance (Guy Major et al., 1993), so under voltage clamp, small changes in R_ser_ could differentially affect minis (with ‘their’ noise) and noise recorded later, following any non-stationarities.

Despite some of the limitations discussed above, the method and its application to neocortical excitatory connections presented in this article provide a useful advance in the field of synaptic physiology. Logical next steps could include the application of this novel quantal analysis method to establish the properties of other types of excitatory synapse, and various types of inhibitory synapse, in both neocortex and hippocampus, and in other species, and to investigate whether average synaptic ‘size’ (injected charge) is more widely conserved within different cell classes.

## Acknowledgements

Cardiff University (GM, MD), The Medical Research Council of the UK (MD PhD scholarship). Neural activity simulations and minis’ distribution fitting were performed on the Cardiff School of Biosciences’ Biocomputing Hub HPC/Cloud infrastructure.

## Abbreviations

ABF: Axon Binary File
aCSF: artificial cerebrospinal fluid
AMPA: α-amino-3-hydroxy-5-methyl-4-isoxazolepropionic acid
AMPAR: α-amino-3-hydroxy-5-methyl-4-isoxazolepropionic acid receptor
ANN: artificial neural network
AP: action potential
CGP55845A: selective competitive antagonist of GABA_B_ receptors (thus slow K^+^ dependent IPSPs)
CPP: (*RS*)-3-(2-carboxypiperazin-4-yl)-propyl-1-phosphonic acid (potent selective competitive glutamate NMDA receptor antagonist, thus depolarization-dependent, slow component of EPSPs)
EPSC: excitatory postsynaptic current
EPSP: excitatory postsynaptic potential
GA: genetic algorithm
GABA: gamma-aminobutyric acid (inhibitory neurotransmitter)
IC50: concentration blocking half of the receptors
IPSC: inhibitory post-synaptic current
IPSP: inhibitory post-synaptic potential
MAD: maximum absolute deviation
mEPSC: miniature excitatory post-synaptic current
mEPSP: miniature excitatory post-synaptic potential
mIPSP: miniature inhibitory post-synaptic potential
mPSC: miniature post-synaptic current
mPSP: miniature post-synaptic potential
NBQX: 2,3-dioxo-6-nitro-1,2,3,4-tetrahydrobenzo[*f*]quinoxaline-7-sulfonamide (potent selective competitive antagonist of AMPA (and kainate) glutamate receptors, thus fast component of EPSPs)
NMDA: N-methyl-D-aspartate
NMDAR: N-methyl-D-aspartate receptor
R_ser_: series resistance
SAD: sum of absolute deviations
SD: standard deviation
smEPSP: simulated mEPSP
SSD: sum of signed deviations
SSE: sum of squared errors
TTX: tetrodotoxin (potent selective competitive antagonist of Na^+^ channels, thus action potentials)
V_m_: transmembrane potential

## Supplementary Figures

**Supplementary Figure 1:**
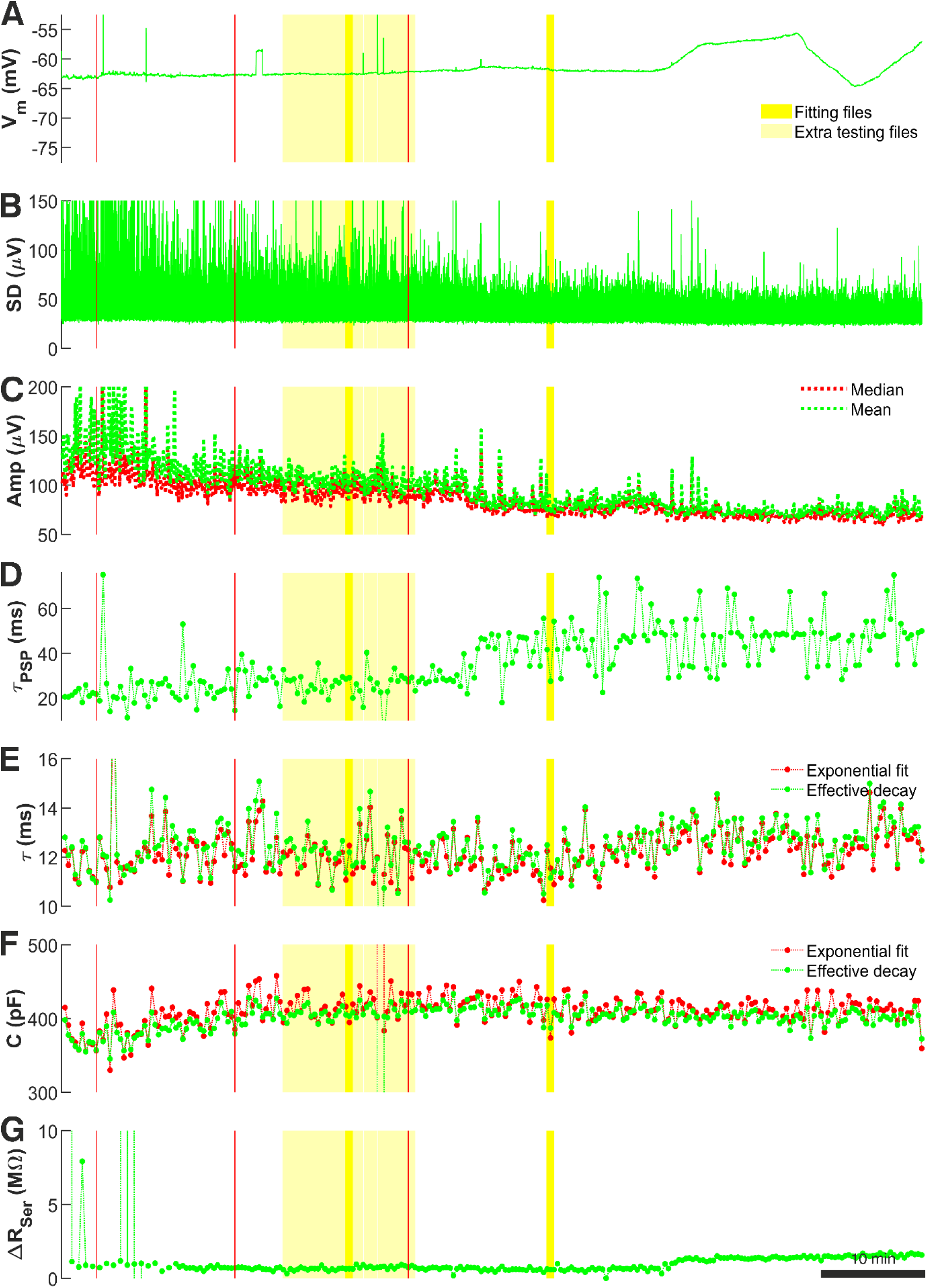
Cell p106b (layer 5). Recording quality measures used to select ’noise with minis’ and ’noise-alone’ sweeps. (A) Baseline V_m_ across entire recording (excluding stimulation pulse periods); duration 4960 s. Red lines and shaded yellow regions and Panels B-G as for Figure 2.

**Supplementary Figure 2:**
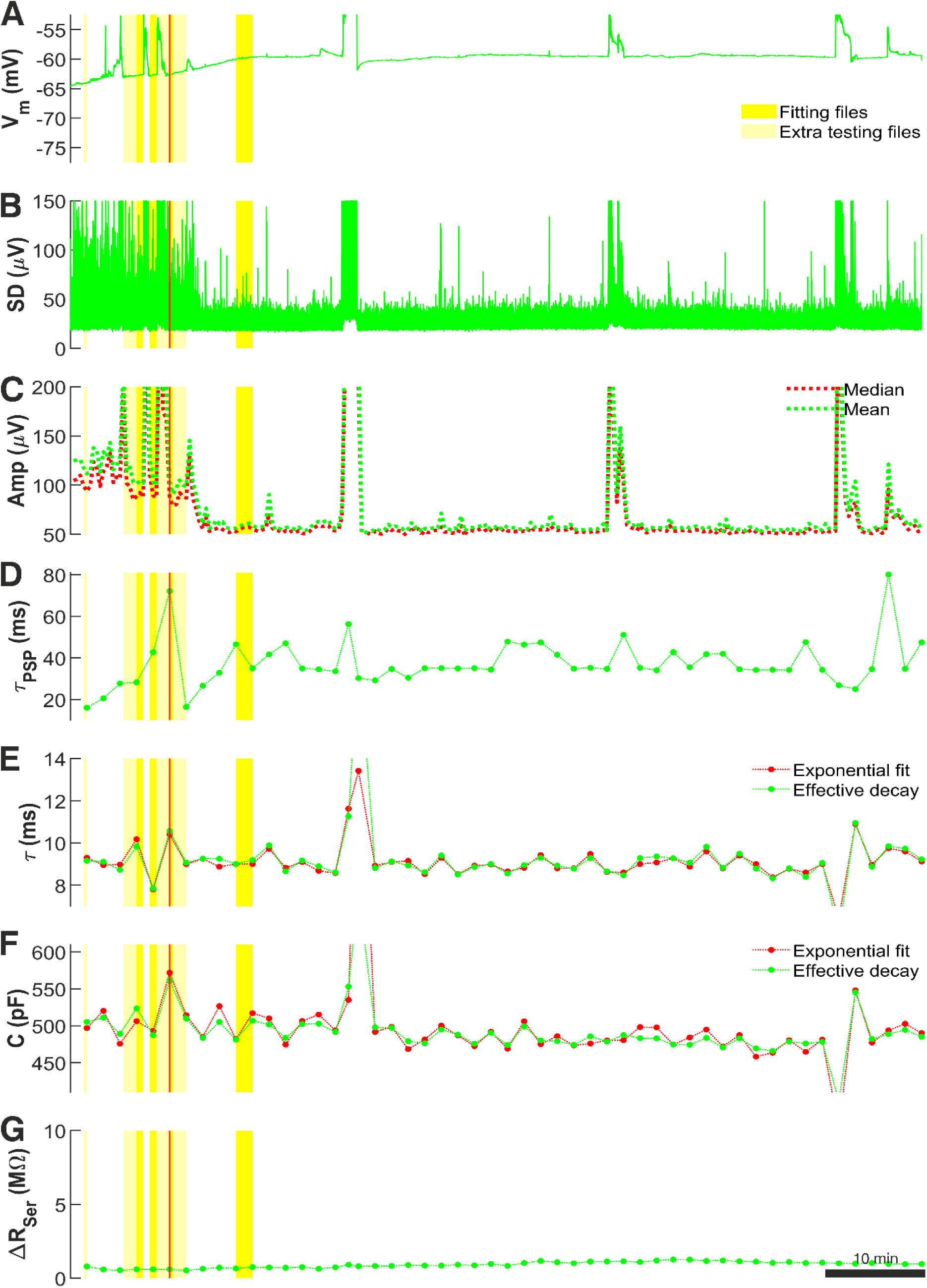
Cell p108a (layer 5). Recording quality measures used to select ’noise with minis’ and ’noise-alone’ sweeps. Panels A-G as for Figure 1 except for data being averaged over 100-second-long windows (5 recording sweeps of 20 s each) in panels D-G. The figure shows only part of the recording following the blockade of APs with only a single red vertical time marker visible indicating the time when pharmacological EPSP blockers were infused. The total duration of the recording shown in the figure was 5140 s. Some episodes with transient recording instabilities (noise glitches) were removed from the ‘noise with minis’ epoch (white gaps).

**Supplementary Figure 3:**
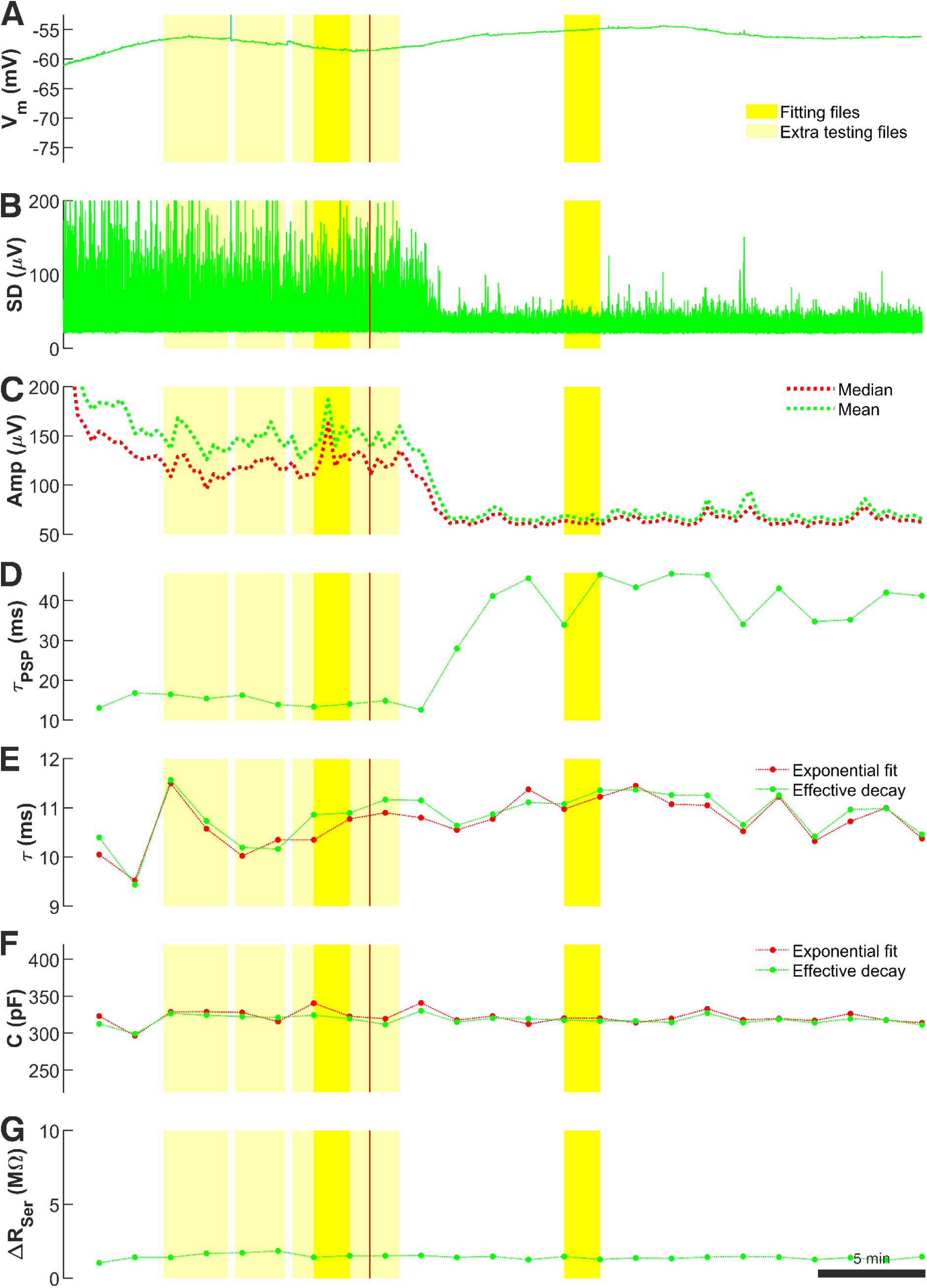
Cell p108b (layer 5). Recording quality measures used to select ’noise with minis’ and ’noise-alone’ sweeps. Panels A-G as for Figure 2 and Supplementary Figure 2. Data were averaged over 100-second-long windows (5 recording sweeps of 20 s each) in panels D-G; duration 2400 s.

**Supplementary Figure 4:**
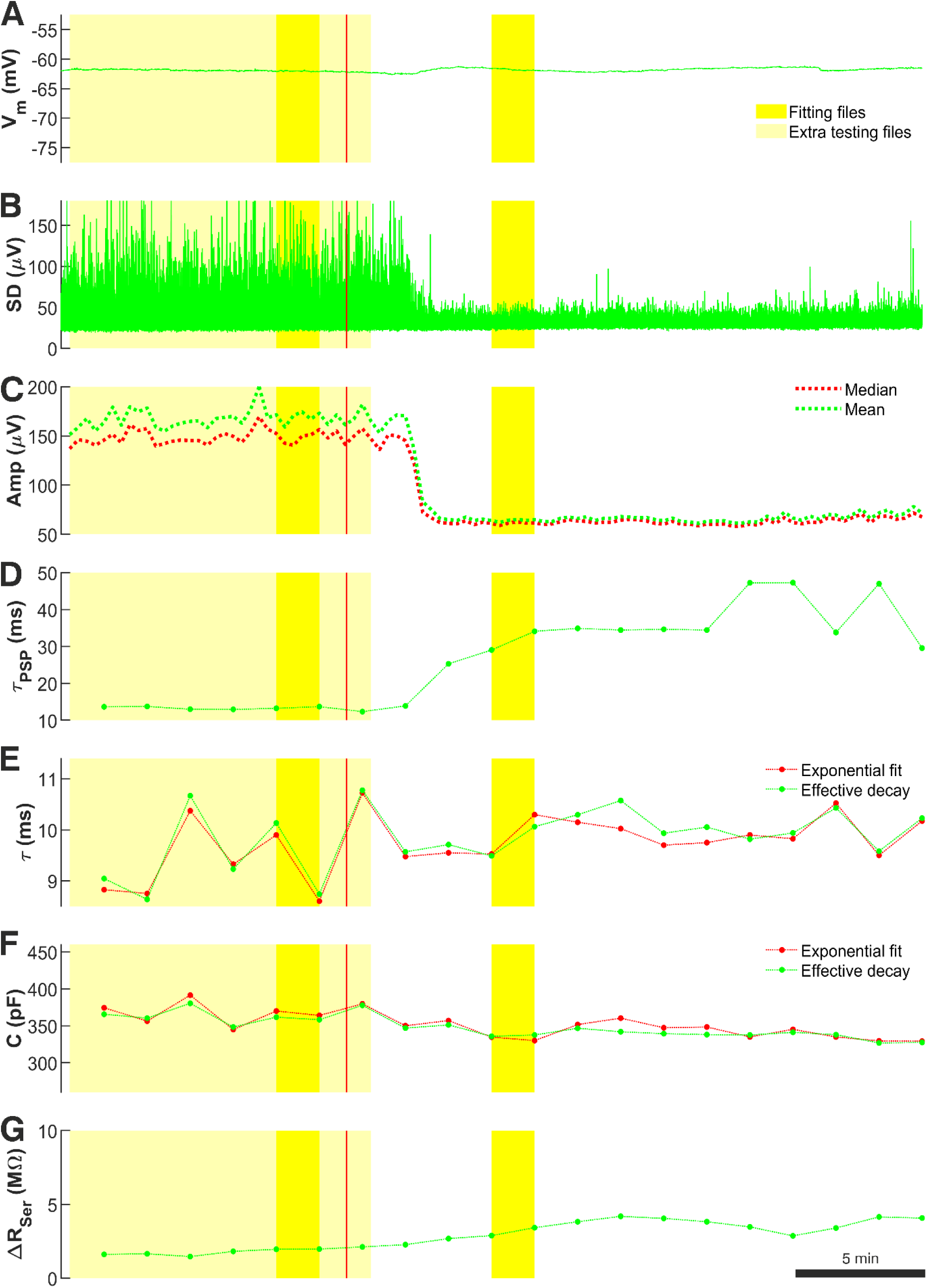
Cell p108c (layer 5). Recording quality measures used to select ’noise with minis’ and ’noise-alone’ sweeps. Panels A-G as for Figure 2 and Supplementary Figure 2. Data were averaged over 100-second-long windows (5 recording sweeps of 20 s each) in panels D-G; duration 2000 s.

**Supplementary Figure 5:**
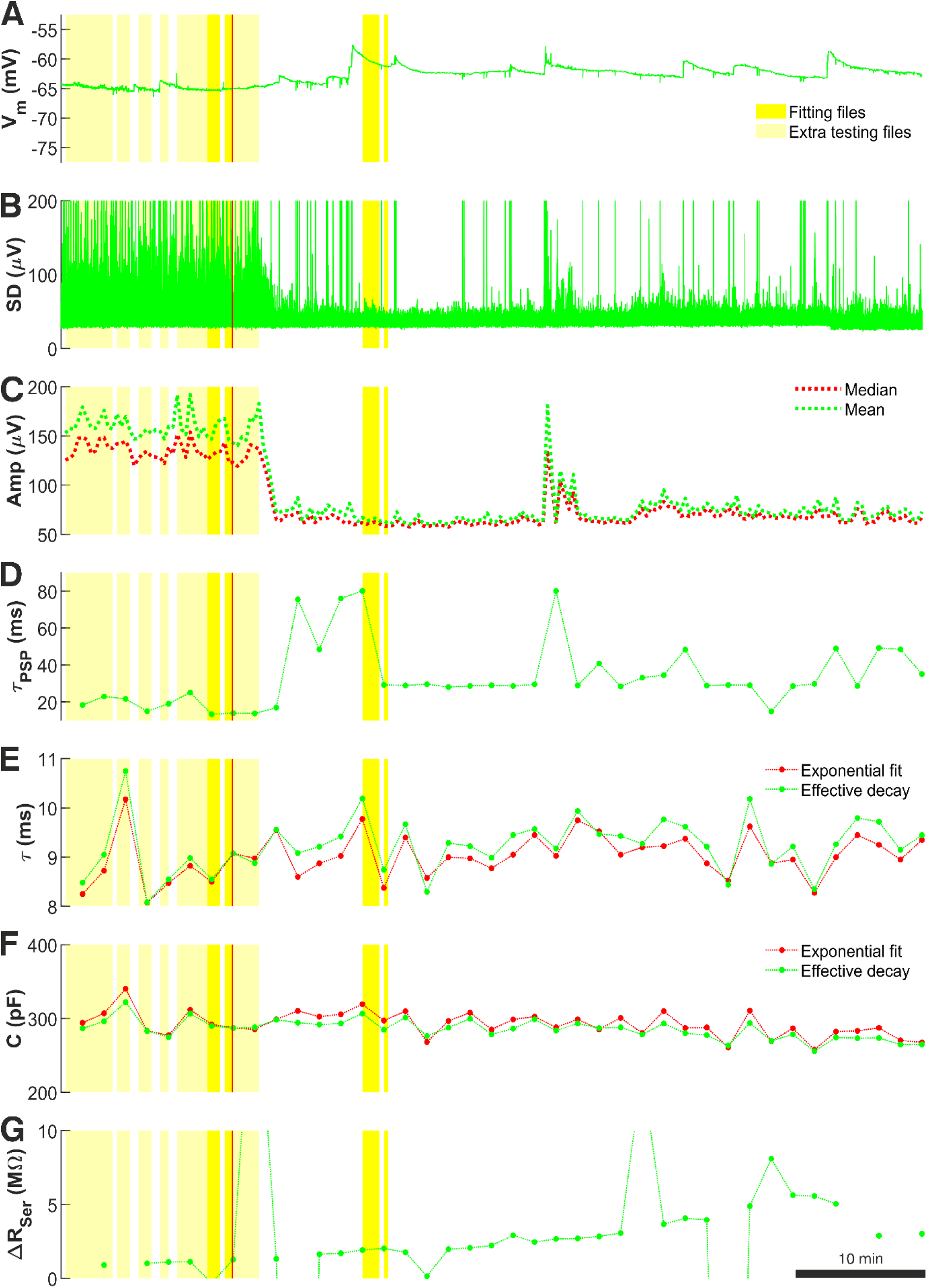
Cell p120b (layer 2/3). Recording quality measures used to select ’noise with minis’ and ’noise-alone’ sweeps. Panels A-G as for Figure 2 and Supplementary Figure 2. Data were averaged over 100-second-long windows (5 recording sweeps of 20 s each) in panels D-G; duration 4000 s. Some transient ‘glitchy’ extra noisy periods (due to transient recording instability) were manually excluded from the ‘noise with minis’ and ‘noise-alone’ epochs (white gaps).

**Supplementary Figure 6:**
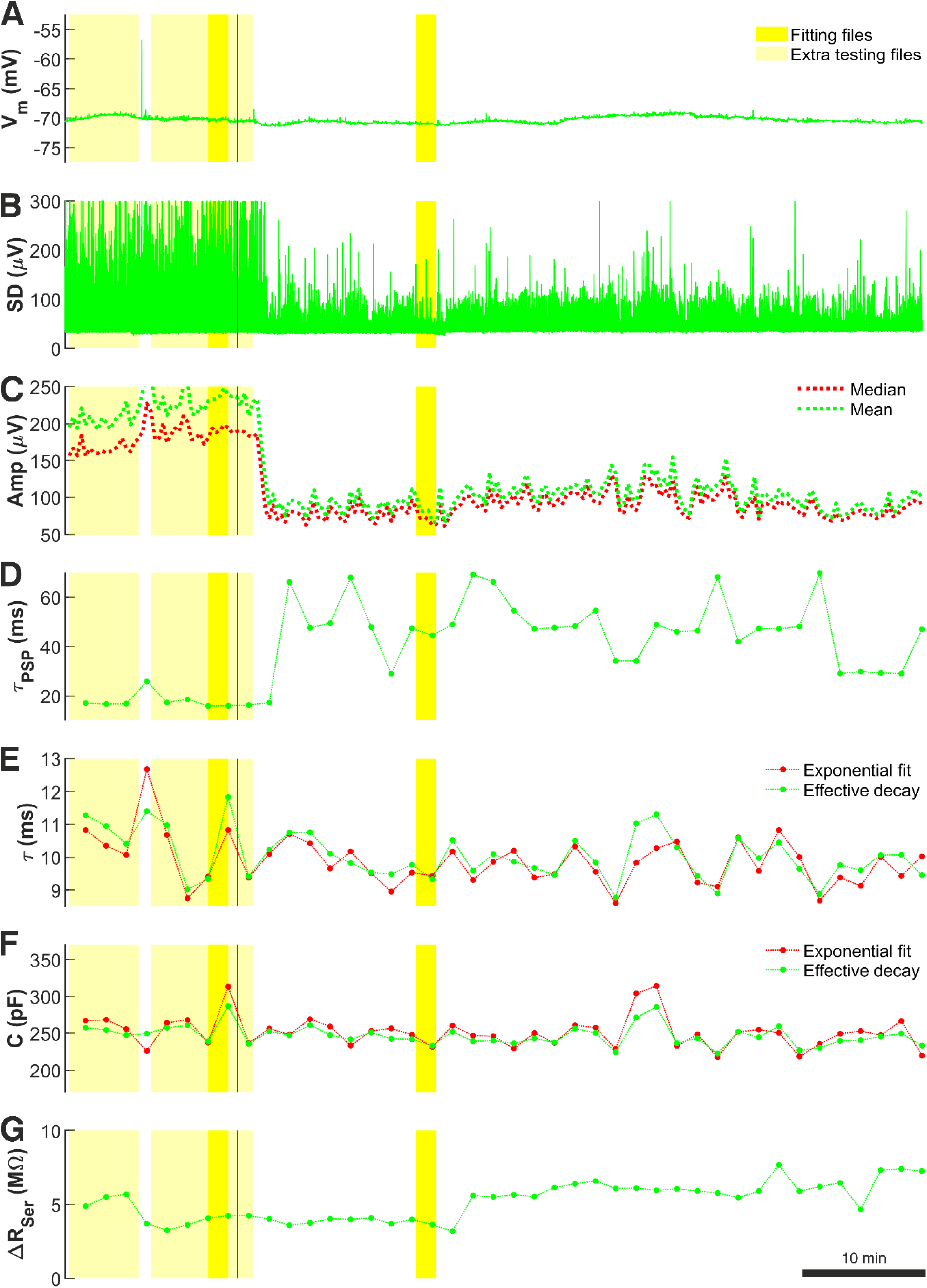
Cell p122a (layer 2/3). Recording quality measures used to select ’noise with minis’ and ’noise-alone’ sweeps. Panels A-G as for Figure 2 and Supplementary Figure 2. Data were averaged over 100-second-long windows (5 recording sweeps of 20 s each) in panels D-G; duration 4200 s.

**Supplementary Figure 7:**
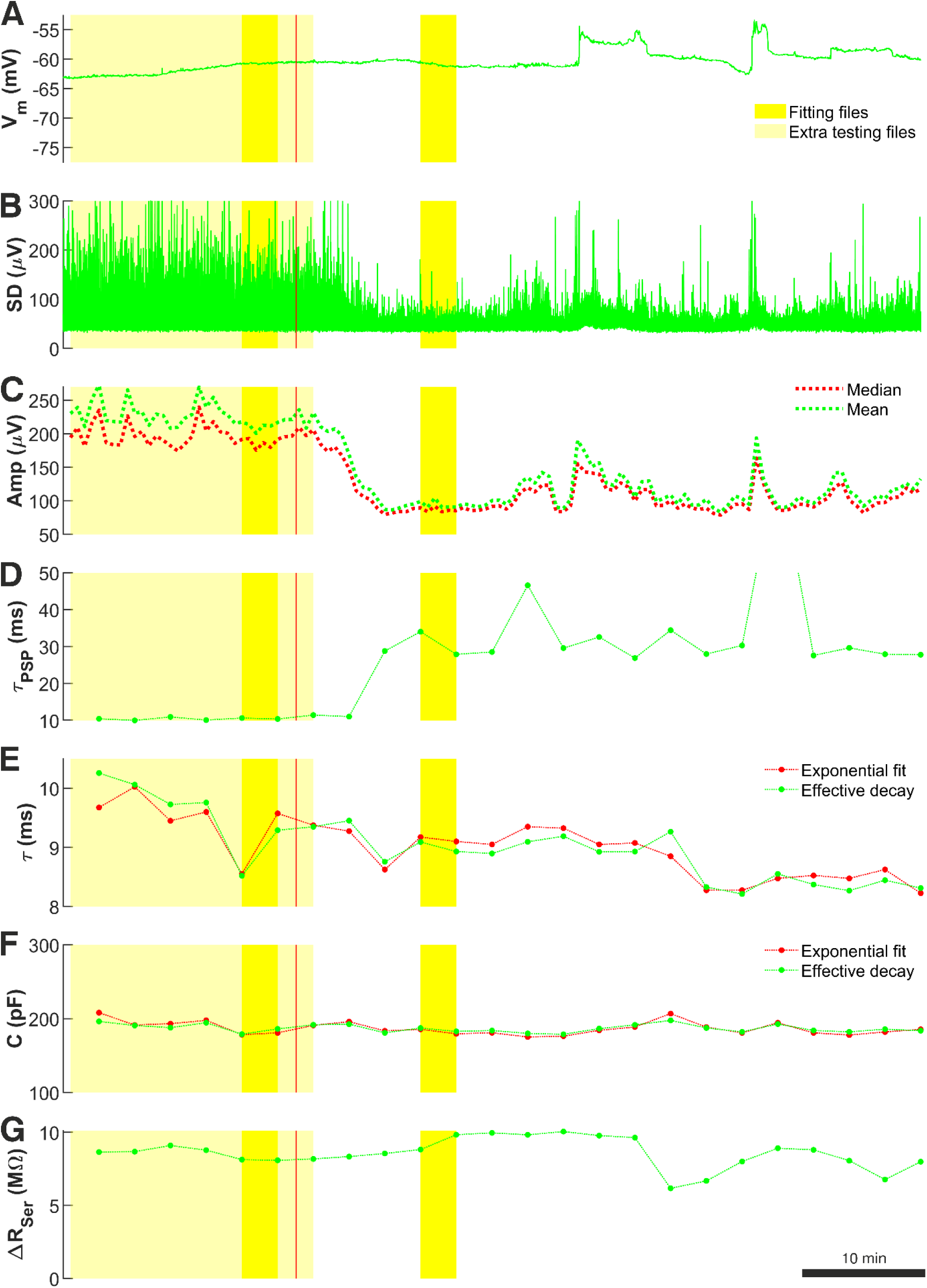
Cell p124b (layer 5). Recording quality measures used to select ’noise with minis’ and ’noise-alone’ sweeps. Panels A-G as for Figure 2 and Supplementary Figure 2. Data were averaged over 100-second-long windows (5 recording sweeps of 20 s each) in panels D-G; duration 2400 s.

**Supplementary Figure 8:**
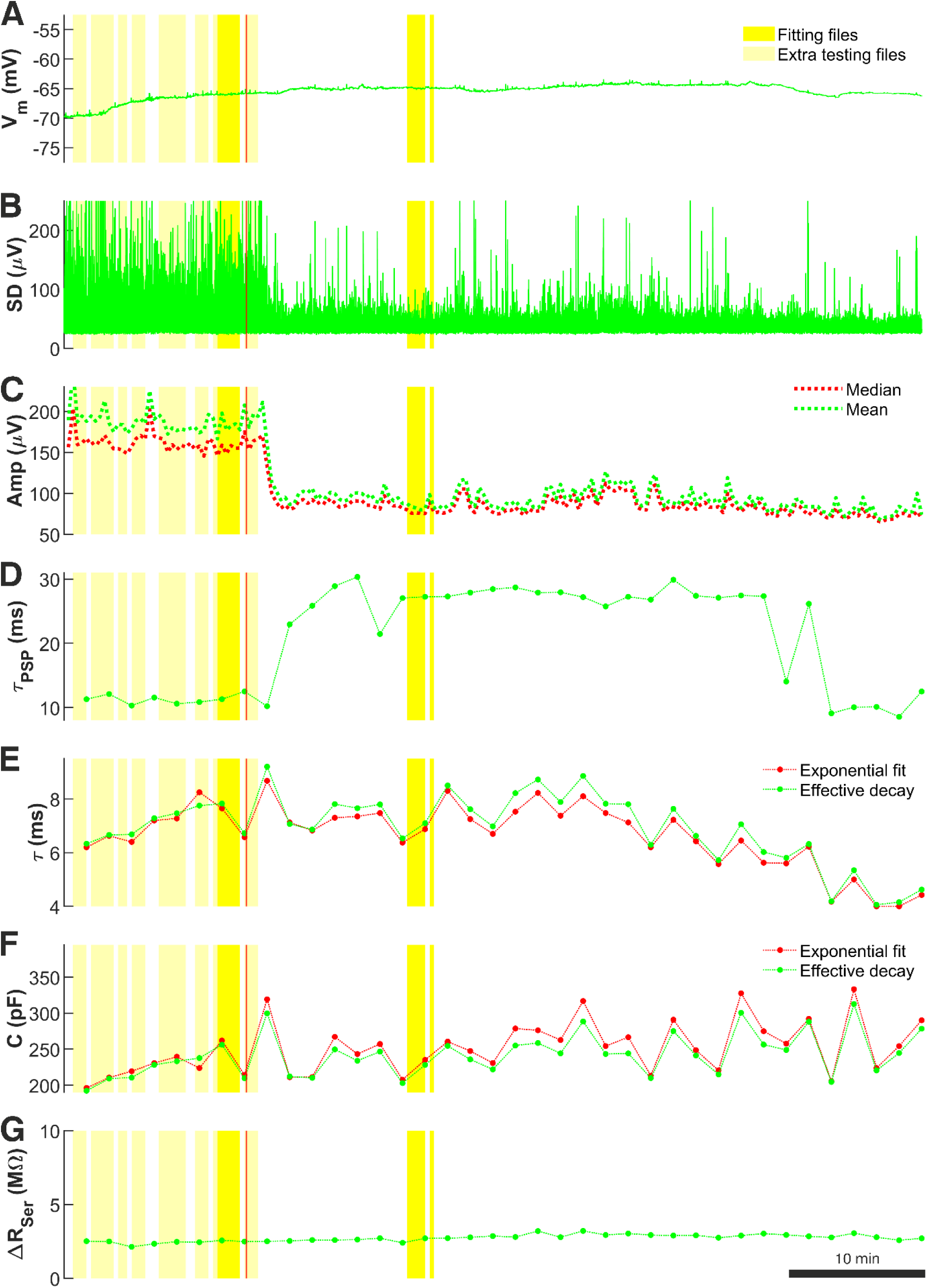
Cell p125a (layer 2/3). Recording quality measures used to select ’noise with minis’ and ’noise-alone’ sweeps. Panels A-G as for Figure 2 and Supplementary Figure 2. Data were averaged over 100-second-long windows (5 recording sweeps of 20 s each) in panels D-G; duration 3800 s. Some transiently extra noisy data due to temporary recording instability were excluded from the ‘noise-alone’ epoch.

**Supplementary Figure 9:**
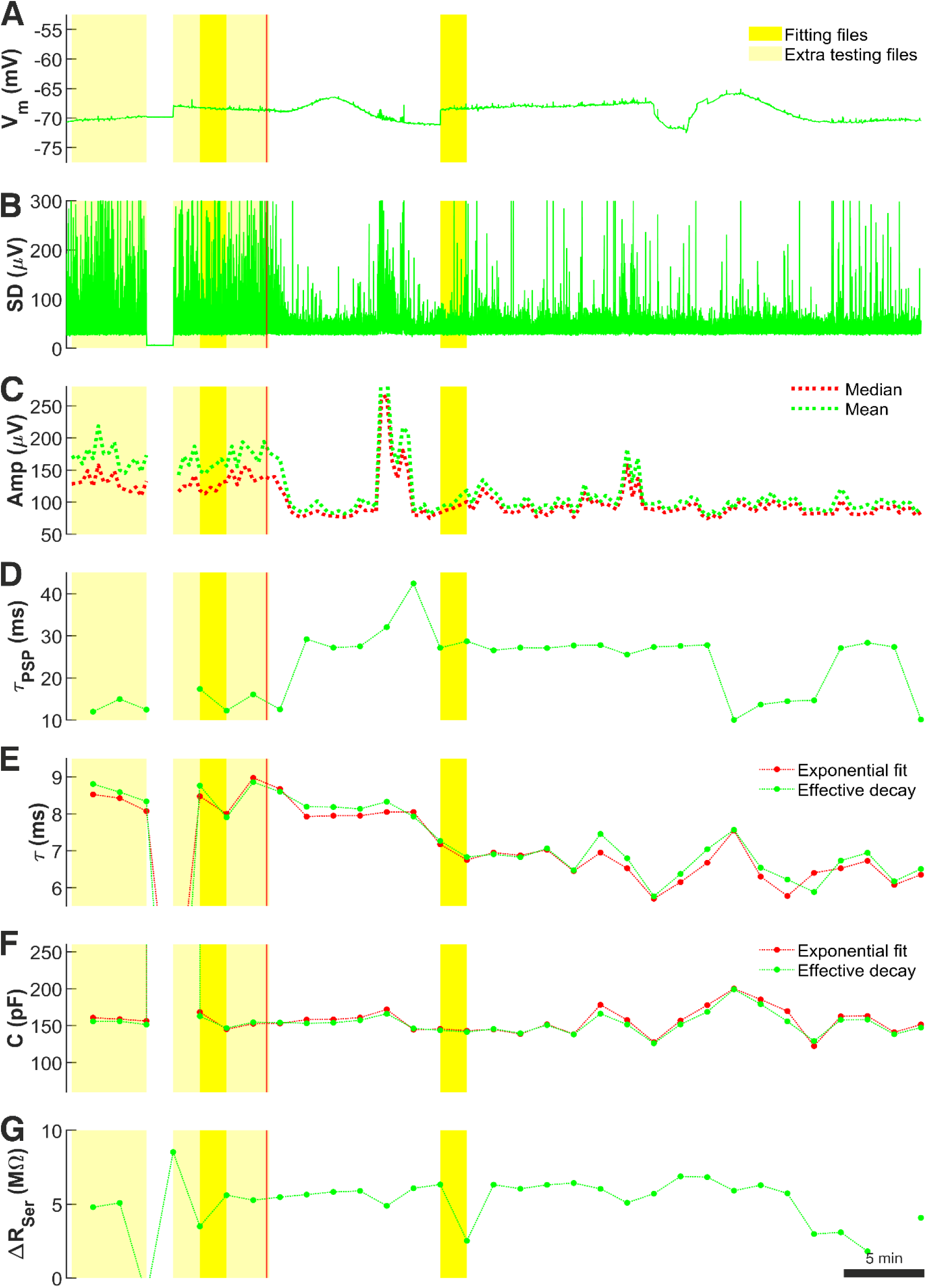
Cell p127c (layer 2/3). Recording quality measures used to select ’noise with minis’ and ’noise-alone’ sweeps. Panels A-G as for Figure 2 and Supplementary Figure 2. Data were averaged over 100-second-long windows (5 recording sweeps of 20 s each) in panels D-G; duration 3200 s.

**Supplementary Figure 10:**
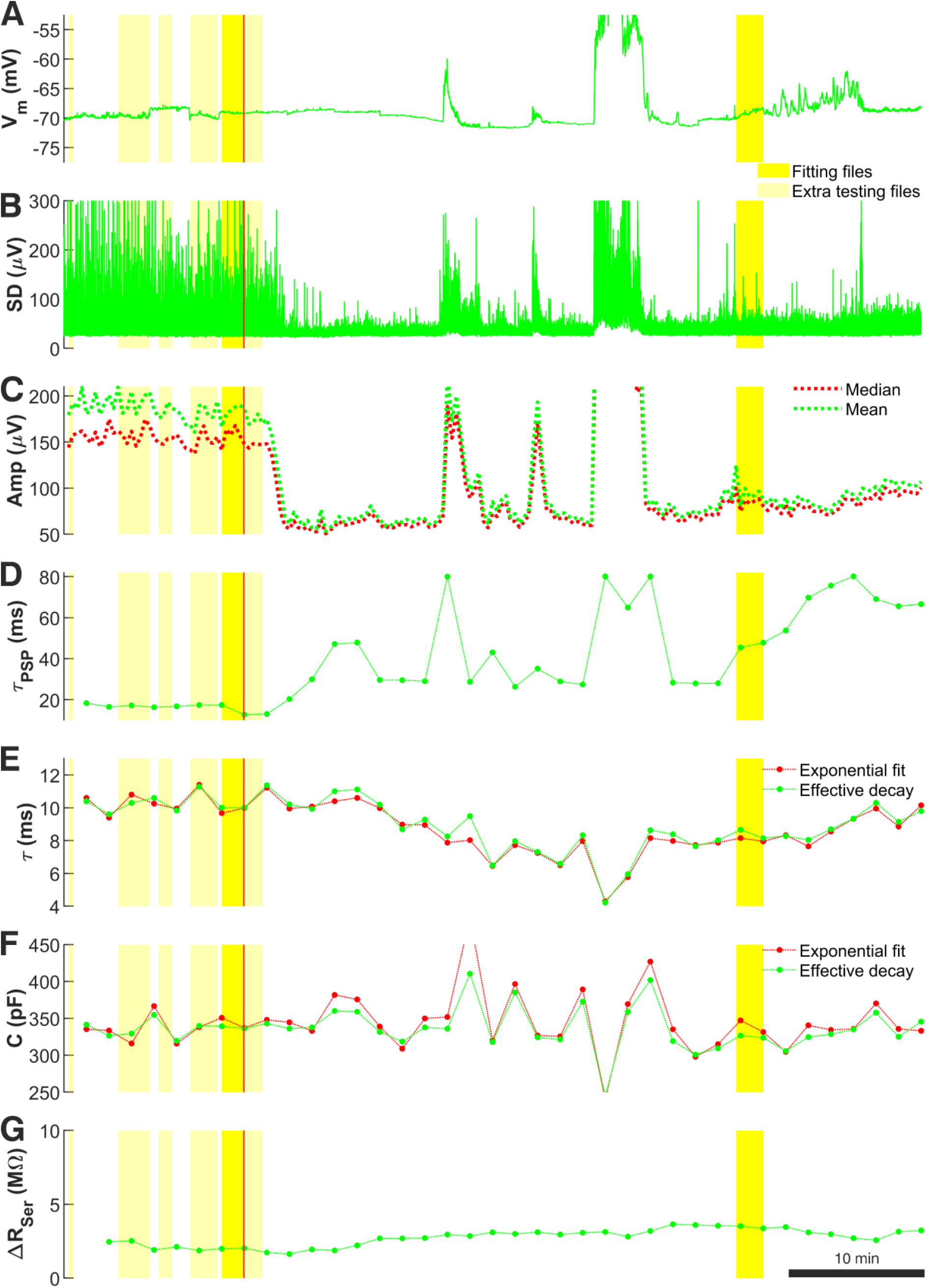
Cell p128c (layer 2/3). Recording quality measures used to select ’noise with minis’ and ’noise-alone’ sweeps. Panels A-G as for Figure 2 and Supplementary Figure 2. Data were averaged over 100-second-long windows (5 recording sweeps of 20 s each) in panels D-G; duration 3800 s.

**Supplementary Figure 11:**
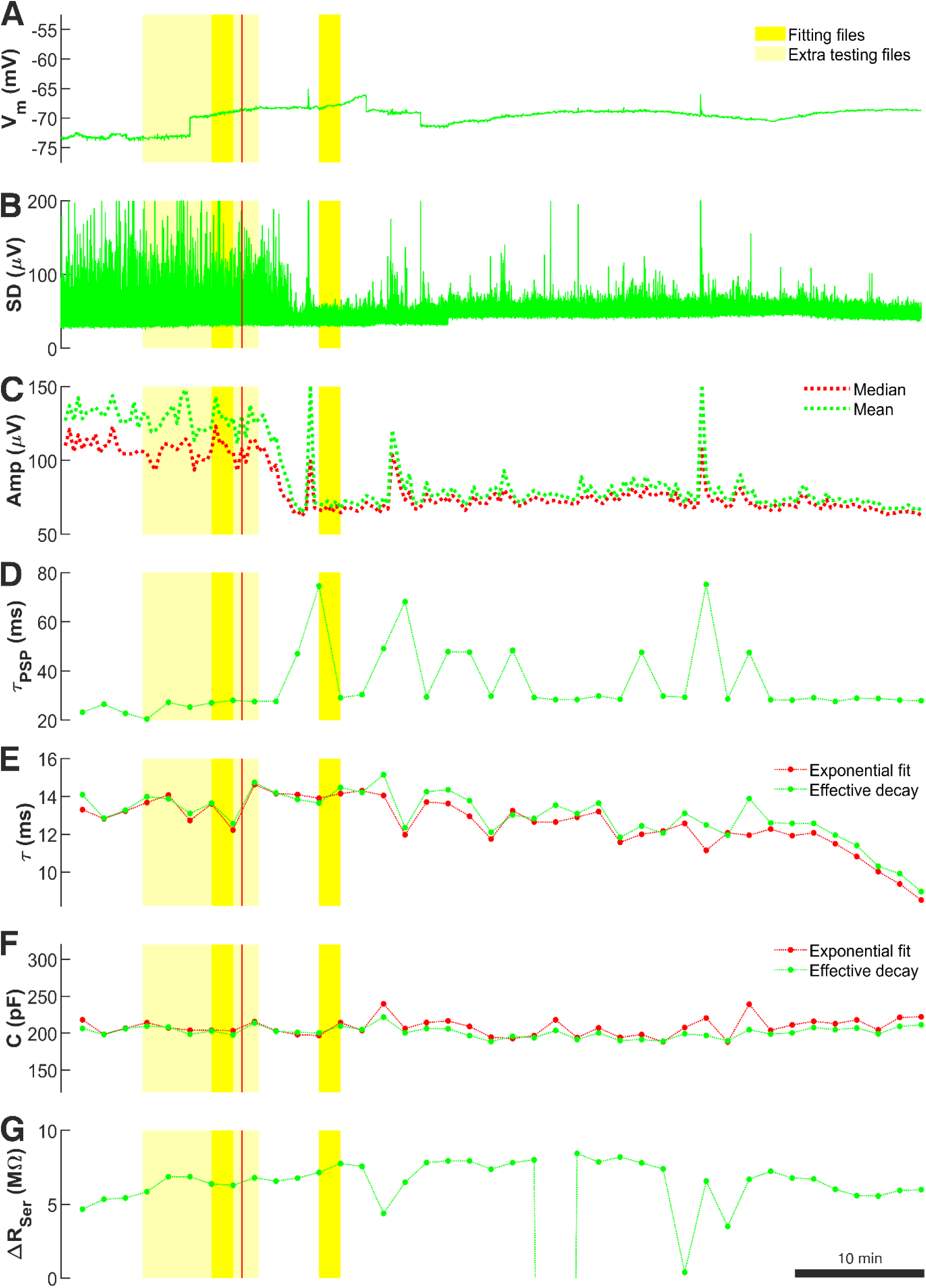
Cell p129a (layer 5). Recording quality measures used to select ’noise with minis’ and ’noise-alone’ sweeps. Panels A-G as for Figure 2 and Supplementary Figure 2. Data were averaged over 100-second-long windows (5 recording sweeps of 20 s each) in panels D-G; duration 4000 s.

**Supplementary Figure 12:**
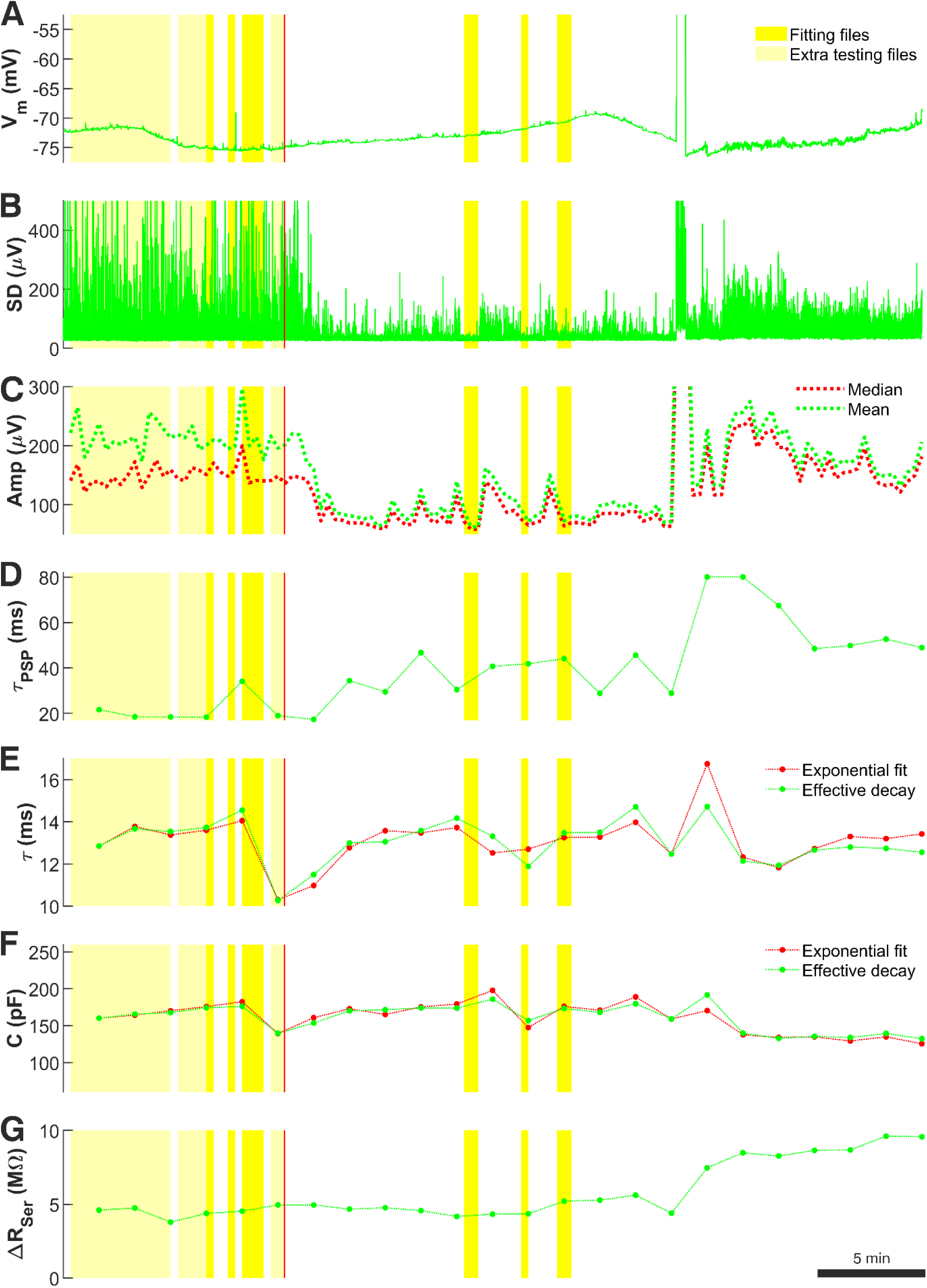
Cell p131a (layer 2/3). Recording quality measures used to select ’noise with minis’ and ’noise-alone’ sweeps. Panels A-G as for Figure 2 and Supplementary Figure 2. Data were averaged over 100-second-long windows (5 recording sweeps of 20 s each) in panels D-G; duration 2400 s. Fit epoch is fragmented due to noisy segments, with transient recording instability, being excluded.

**Supplementary Figure 13:**
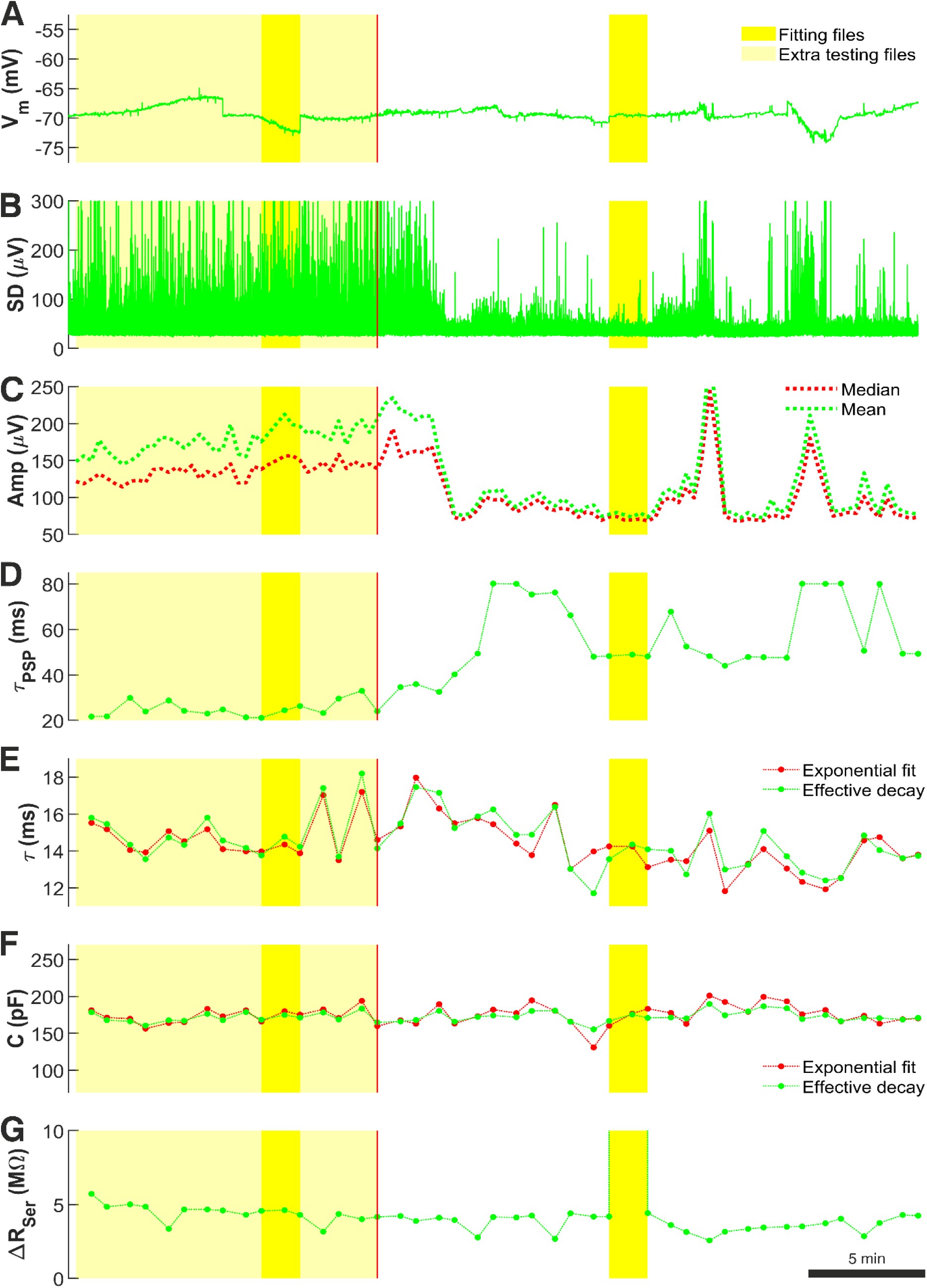
Cell p131c (layer 2/3). Recording quality measures used to select ’noise with minis’ and ’noise-alone’ sweeps. Panels A-G as for Figure 2 and Supplementary Figure 2. Data were averaged over 100-second-long windows (5 recording sweeps of 20 s each) in panels D-G; duration 2200 s.

**Supplementary figure 14:**
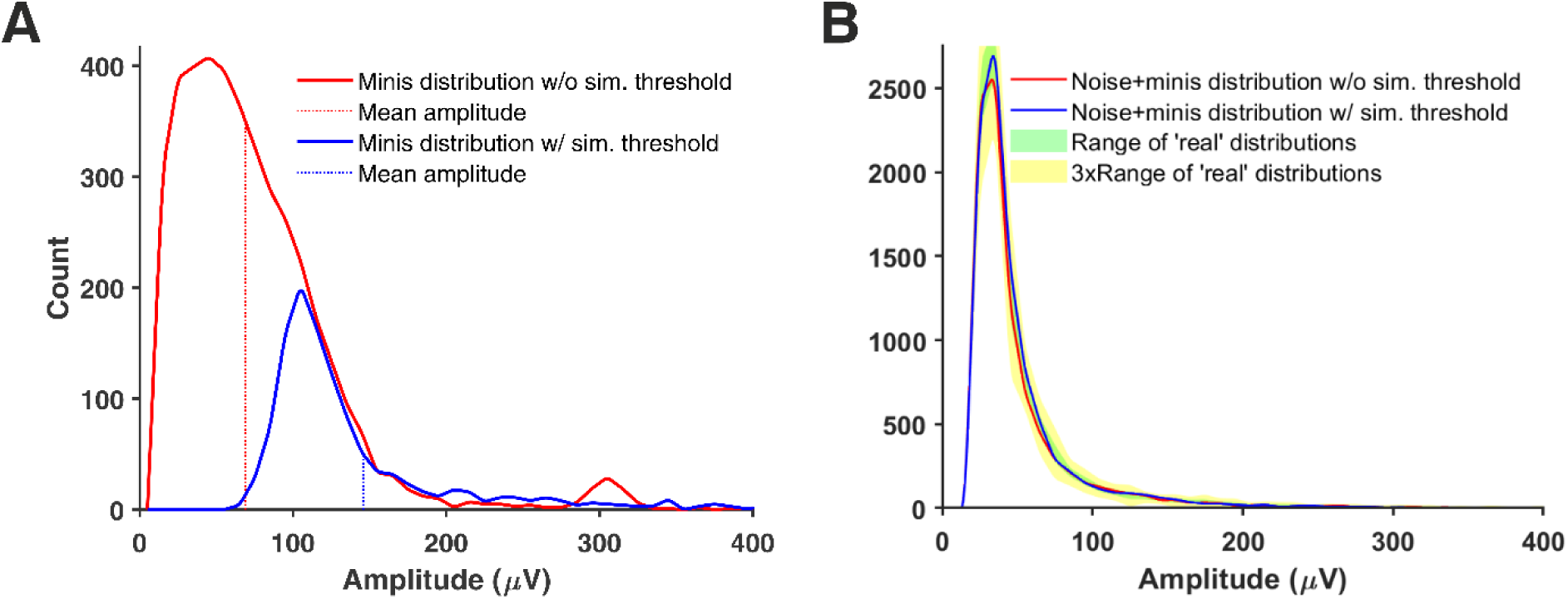
Non-uniqueness of simulated minis distributions. (A) Two amplitude distributions of simulated minis events generated by the GA using a lower limit on simulated amplitudes of 10 µV (red histogram; termed as without simulation threshold) or 60 µV simulation threshold (blue histogram). (B) Distributions of mini-like events detected in a ’noise with simulated minis’ voltage traces corresponding to the two minis source distributions in (A). Event histograms are shown superimposed on the full range of ‘real’ (‘noise with real minis’) distributions. The two distributions clearly fall within the range of ‘real’ distributions albeit being generated by two very different source distributions. Recordings obtained from cell p131c (layer 2/3).

**Supplementary figure 15:**
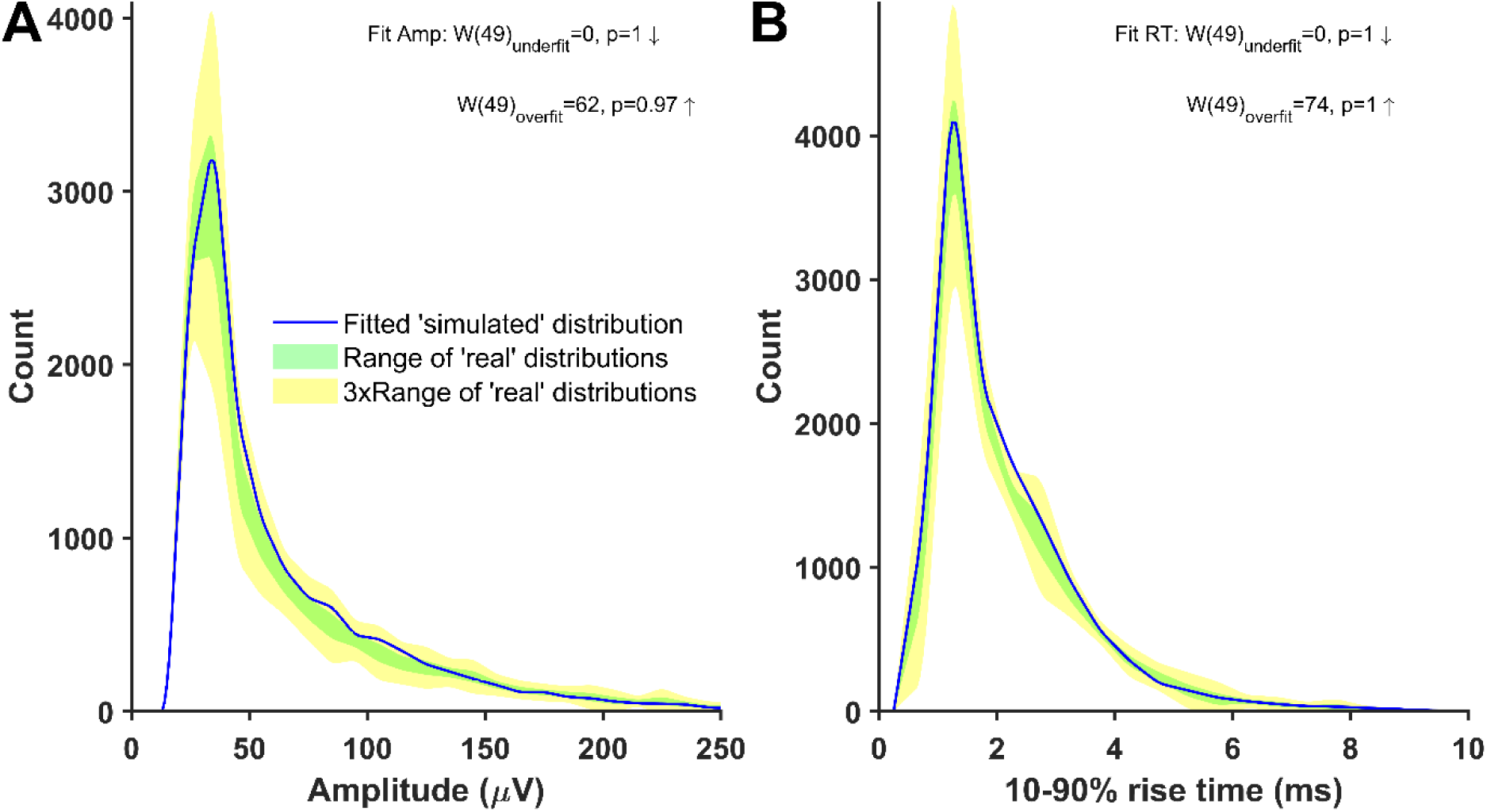
Distributions of mini-like events detected in a ’noise with simulated minis’ voltage trace that closely matched ‘noise with real minis‘ recordings and passed statistical good fit tests, i.e. did not ‘fail’ the poor fit (underfit) unidirectional tests (it could not be ‘picked out’ either objectively or ‘double-blind’ from the ‘noise with real minis’ distributions found with the same detection parameters). Parameters of simulated minis were controlled by the GA; ’noise-alone’ recording obtained from cell p103a (layer 5). Downward arrows indicate ‘simulated’ SAD < ‘real’ SAD. Upward arrows indicate ‘simulated’ SAD > ‘real’ SAD. The ‘underfit’ (poor fit) unidirectional test (top) involves a comparison with the ‘worst-performing’ (highest between-file) ‘real’ SAD file (most discrepant from the others; the null hypothesis is ‘simulated’ SAD does not exceed ‘real’ SAD). The ‘overfit’ unidirectional test (bottom) involves a comparison with the ‘best-performing’ (lowest between-file) ‘real’ SAD (least discrepant from the others; the null hypothesis is ‘simulated’ SAD is not smaller than ‘real’ SAD), but ‘failure’ of this test (p <0.05 Bonferroni-corrected, difference not by chance) does not indicate a fit rejection. (A) Amplitude and (B) 10-90% rise time distributions of mini-like events detected in a ’noise with simulated minis’ voltage trace (blue curve).

**Supplementary Figure 16:**
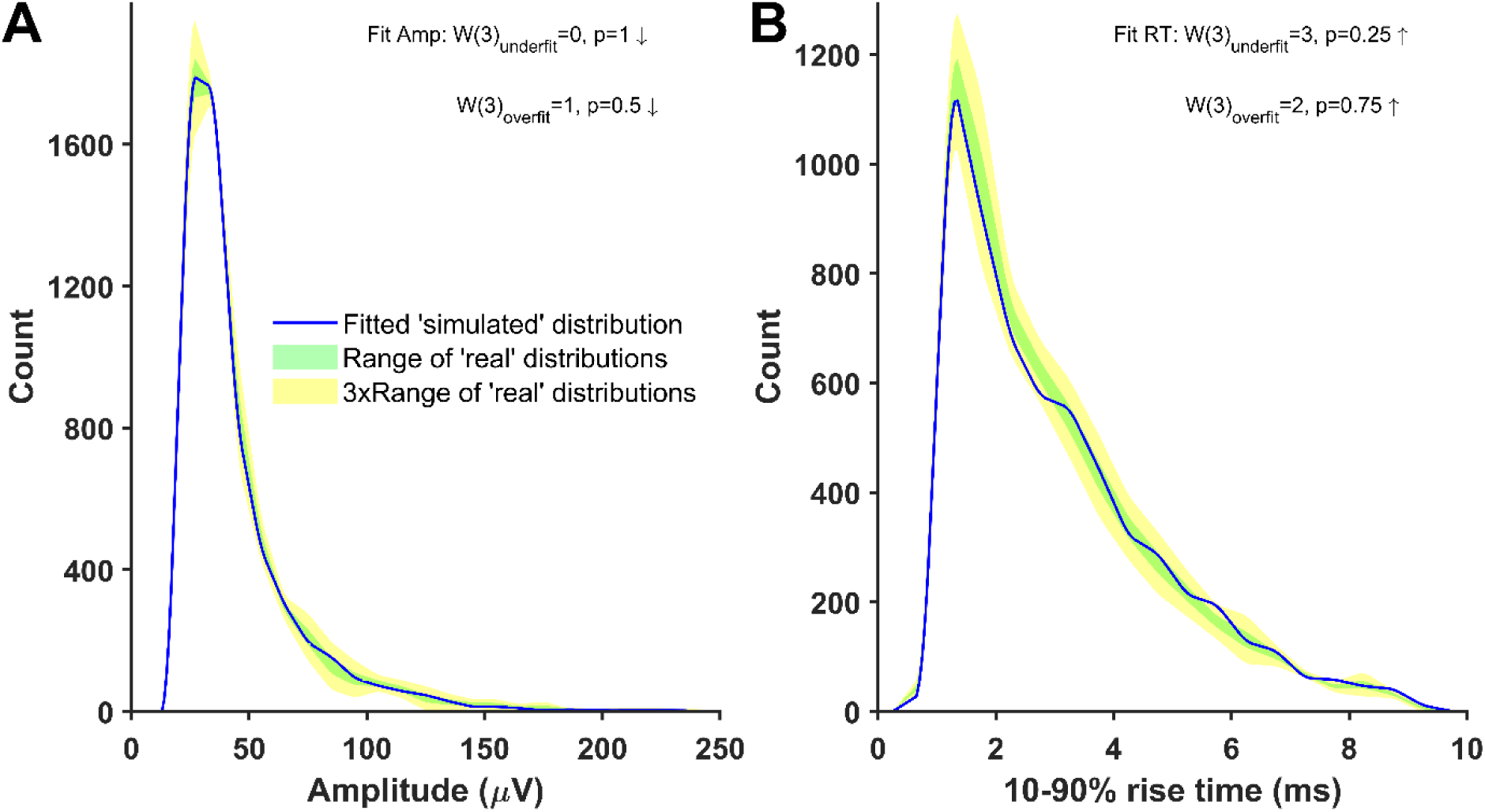
Cell p108a (layer 5), distributions of mini-like events detected in a ’noise with simulated minis’ voltage trace that closely matched ‘noise with real minis‘ recordings. Parameters of simulated minis controlled by GA. Down arrow: ‘simulated’ SAD < ‘real’ SAD (up: ‘simulated’ > ‘real’; Suppl. Fig. 15).

**Supplementary Figure 17:**
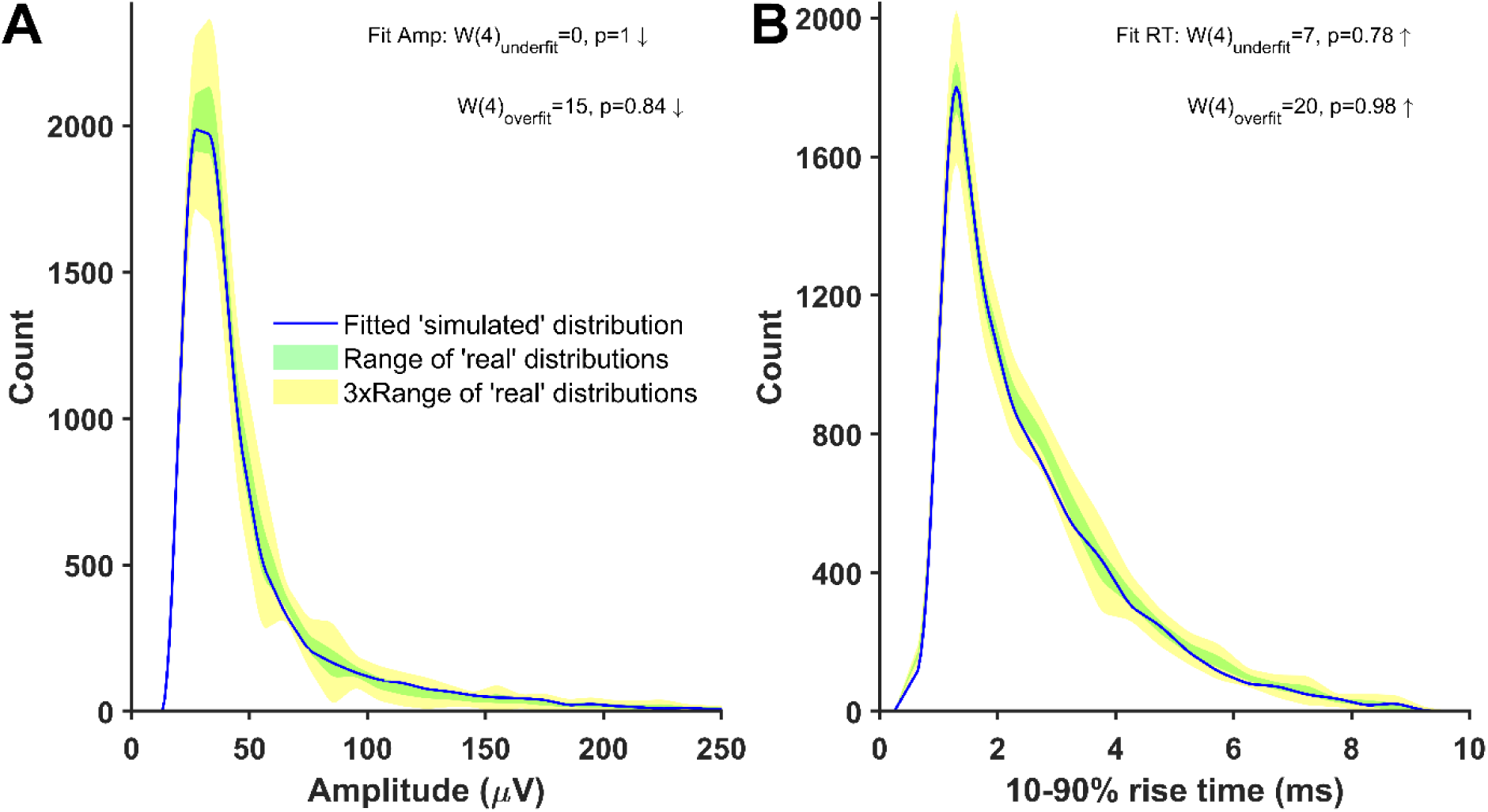
Cell p108b (layer 5), distributions of mini-like events detected in a ’noise with simulated minis’ voltage trace that closely matched ‘noise with real minis‘ recordings. Parameters of simulated minis controlled by GA. Down arrow: ‘simulated’ SAD < ‘real’ SAD (up: ‘simulated’ > ‘real’; Suppl. Fig. 15).

**Supplementary Figure 18:**
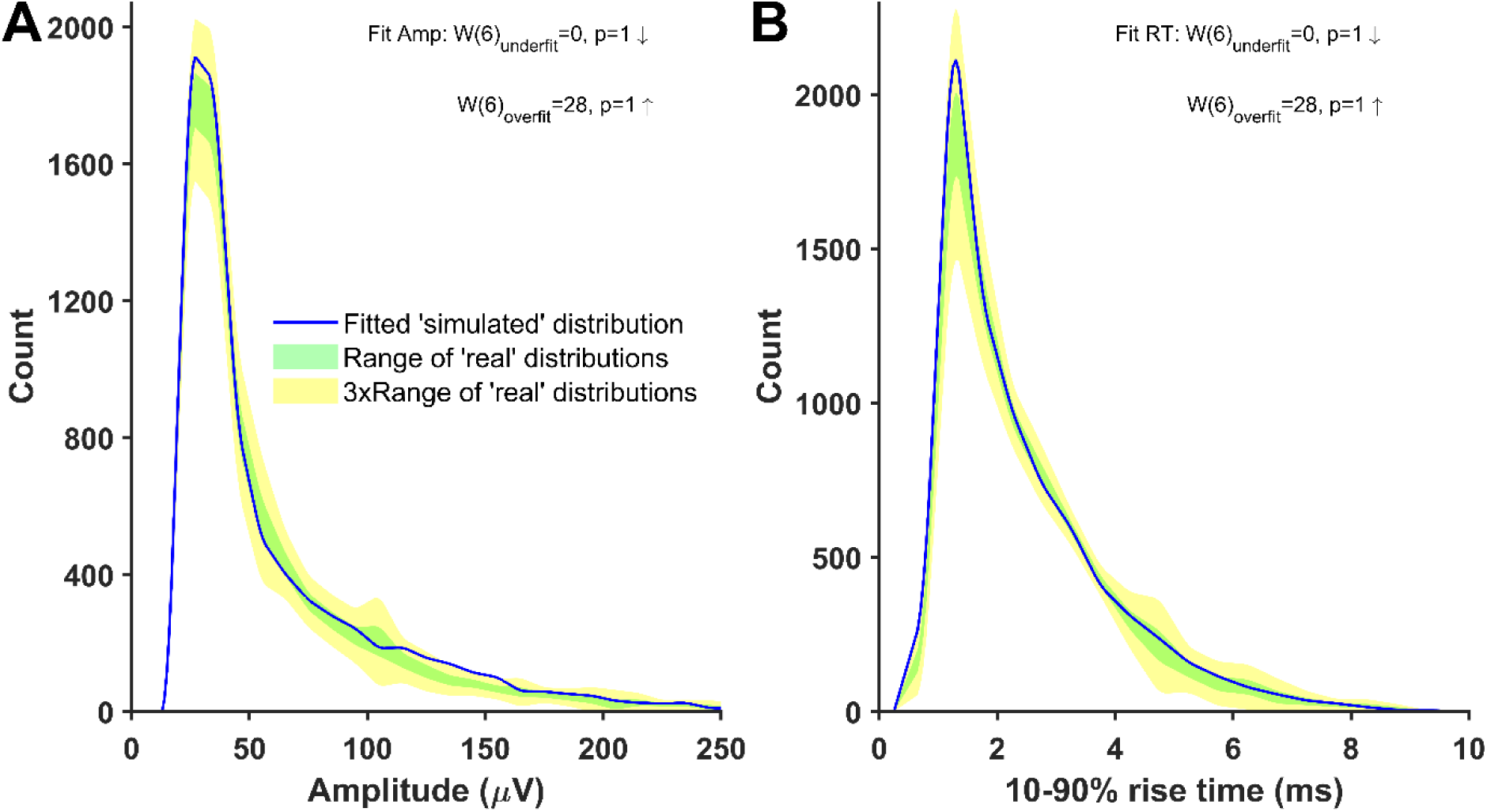
Cell p108c (layer 5), distributions of mini-like events detected in a ’noise with simulated minis’ voltage trace that closely matched ‘noise with real minis‘ recordings. Parameters of simulated minis controlled by GA. Down arrow: ‘simulated’ SAD < ‘real’ SAD (up: ‘simulated’ > ‘real’; Suppl. Fig. 15).

**Supplementary Figure 19:**
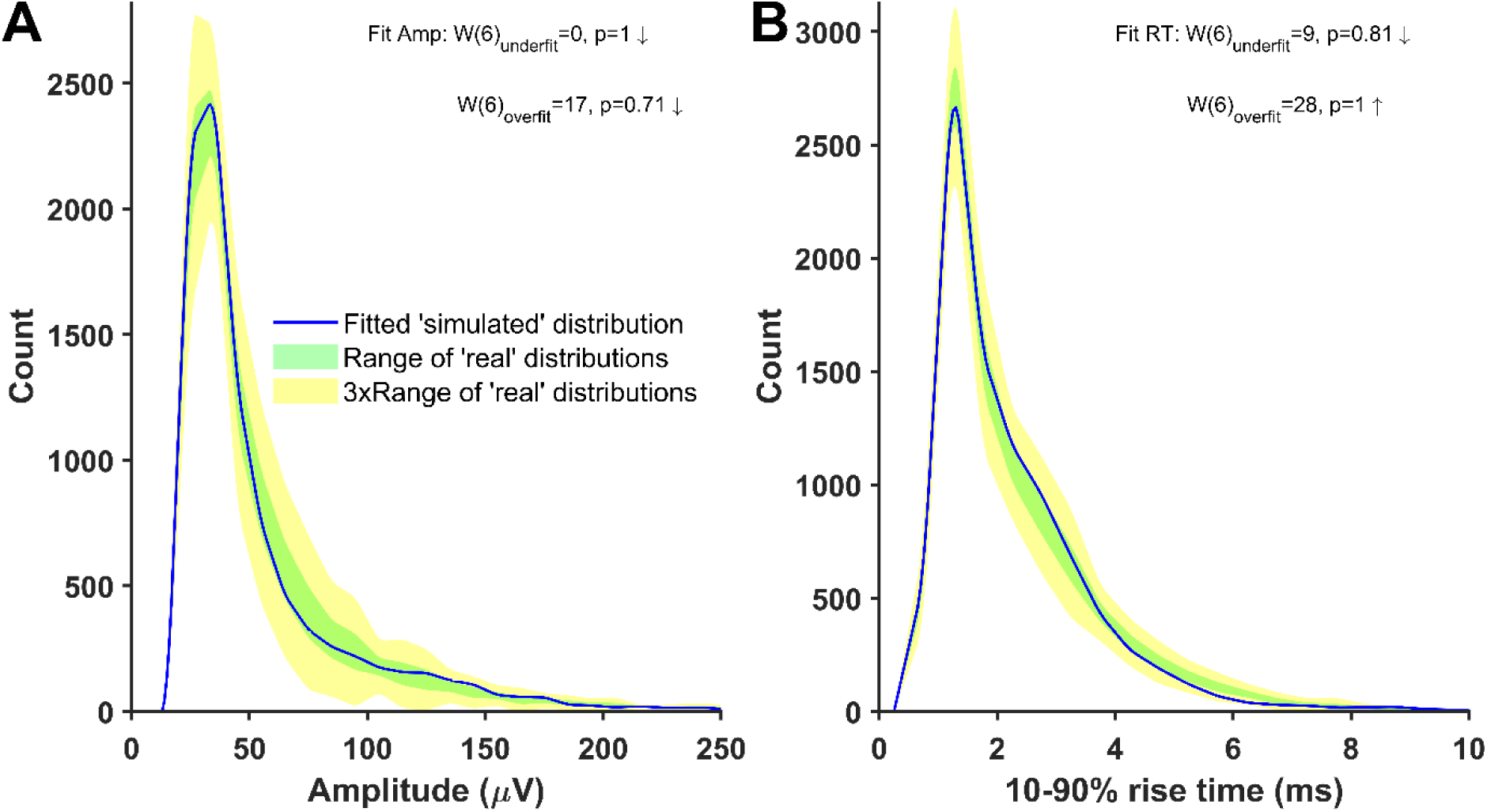
Cell p120b (layer 2/3), distributions of mini-like events detected in a ’noise with simulated minis’ voltage trace that closely matched ‘noise with real minis‘ recordings. Parameters of simulated minis controlled by GA. Down arrow: ‘simulated’ SAD < ‘real’ SAD (up: ‘simulated’ > ‘real’; Suppl. Fig. 15).

**Supplementary Figure 20:**
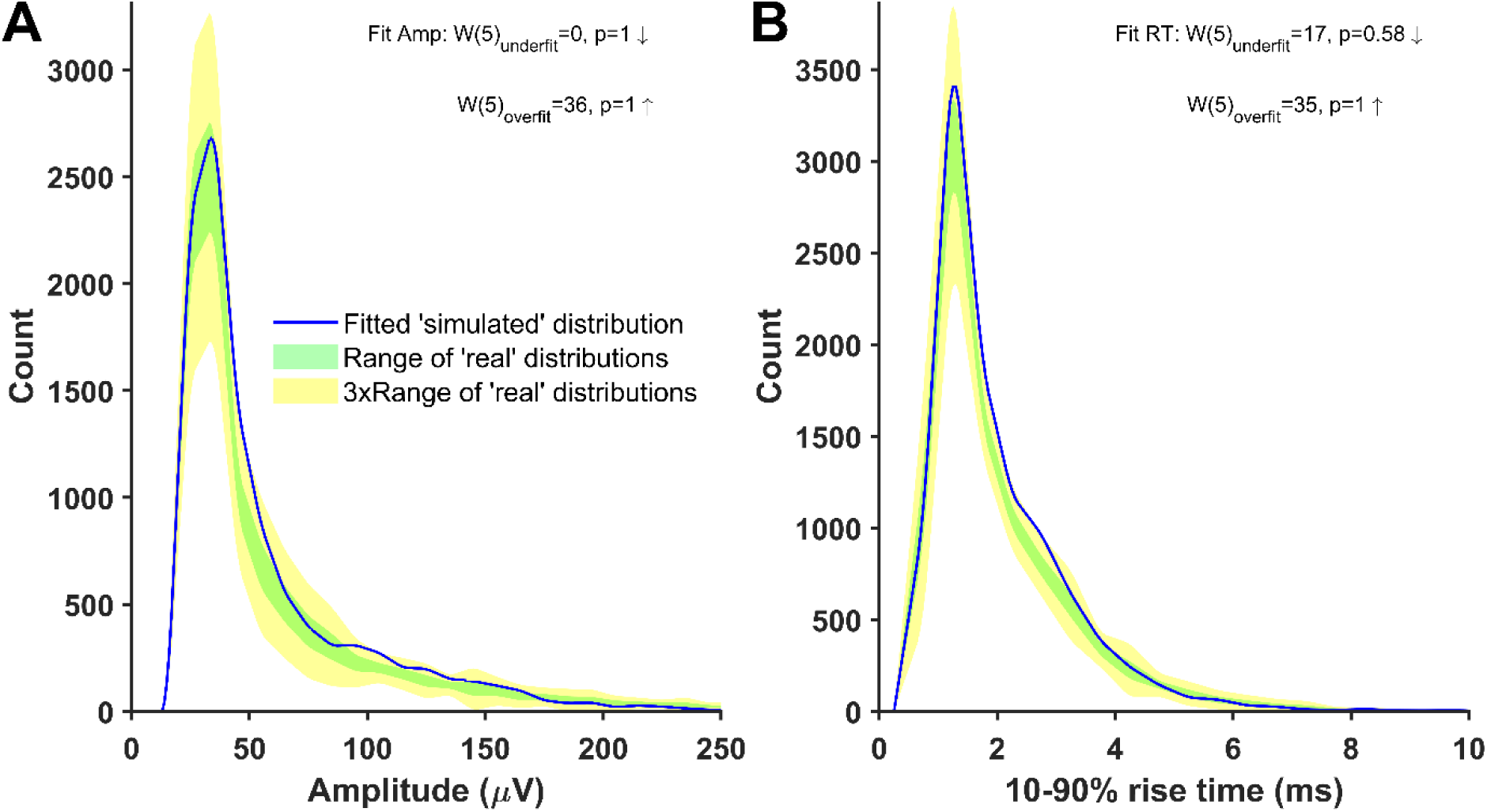
Cell p122a (layer 2/3), distributions of mini-like events detected in a ’noise with simulated minis’ voltage trace that closely matched ‘noise with real minis‘ recordings. Parameters of simulated minis controlled by GA. Down arrow: ‘simulated’ SAD < ‘real’ SAD (up: ‘simulated’ > ‘real’; Suppl. Fig. 15).

**Supplementary Figure 21:**
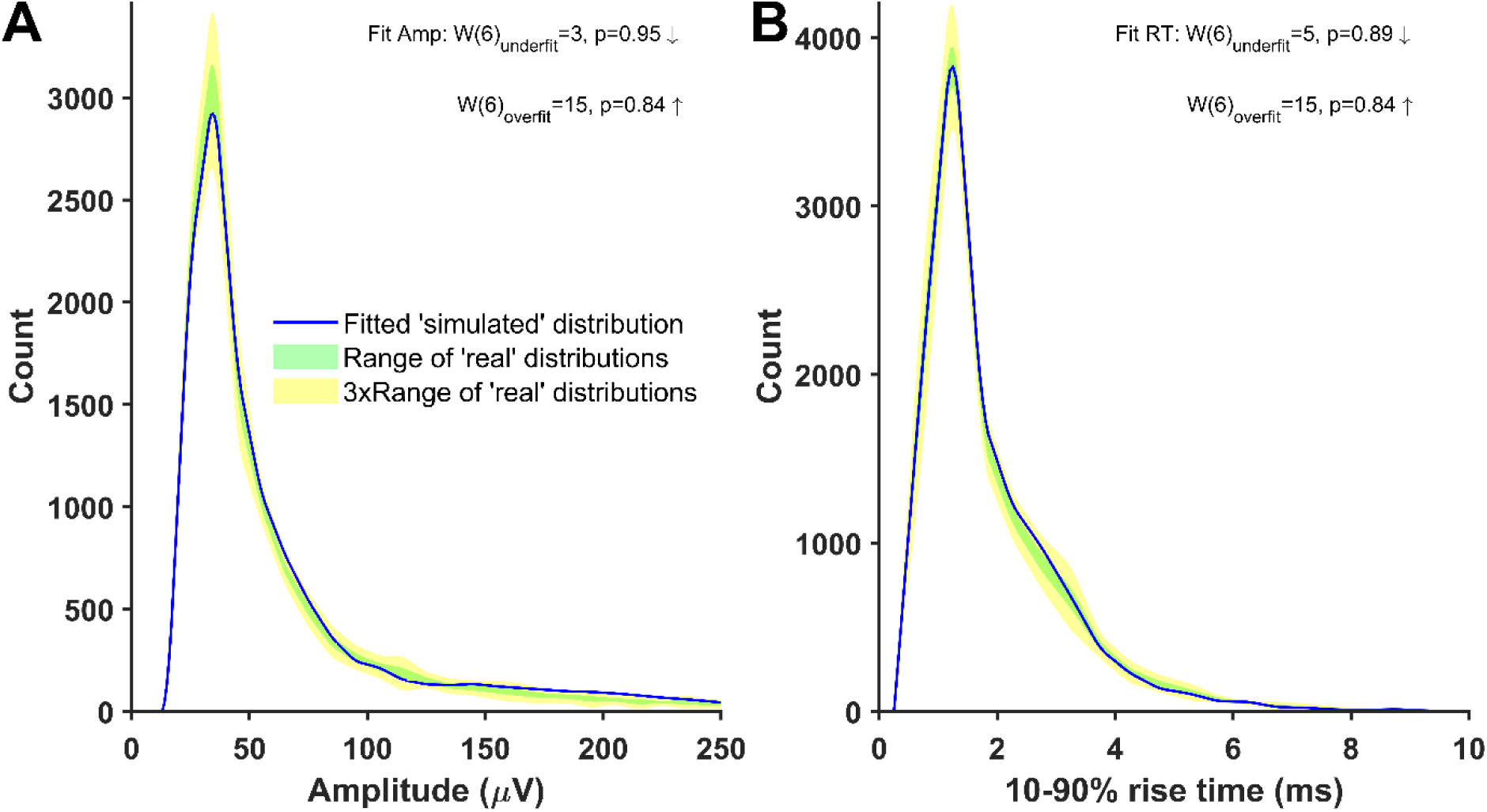
Cell p124b (layer 5), distributions of mini-like events detected in a ’noise with simulated minis’ voltage trace that closely matched ‘noise with real minis‘ recordings. Parameters of simulated minis controlled by GA. Down arrow: ‘simulated’ SAD < ‘real’ SAD (up: ‘simulated’ > ‘real’; Suppl. Fig. 15).

**Supplementary Figure 22:**
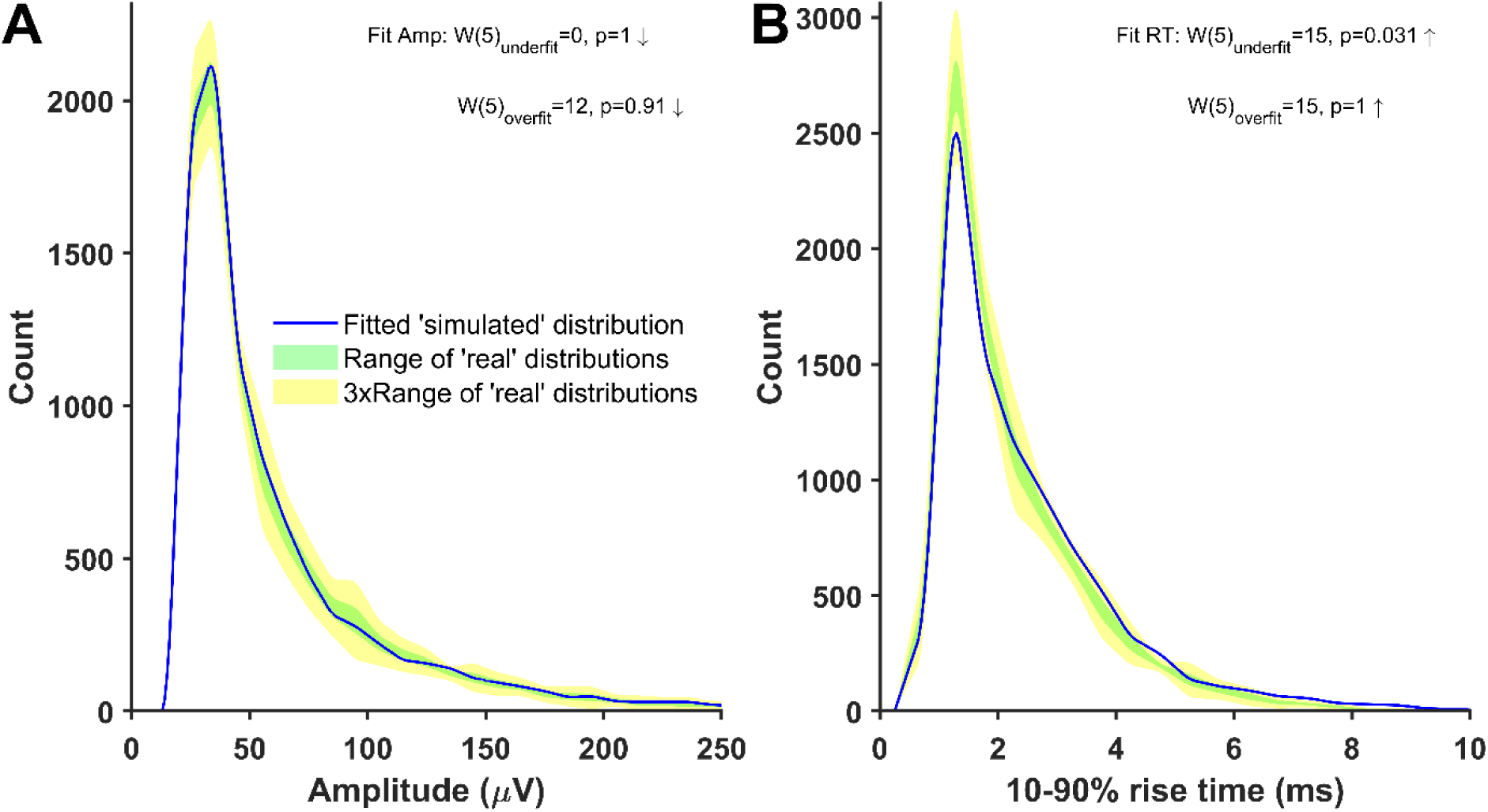
Cell p125a (layer 2/3), distributions of mini-like events detected in a ’noise with simulated minis’ voltage trace that closely matched ‘noise with real minis‘ recordings. Parameters of simulated minis controlled by GA. Down arrow: ‘simulated’ SAD < ‘real’ SAD (up: ‘simulated’ > ‘real’; Suppl. Fig. 15).

**Supplementary Figure 23:**
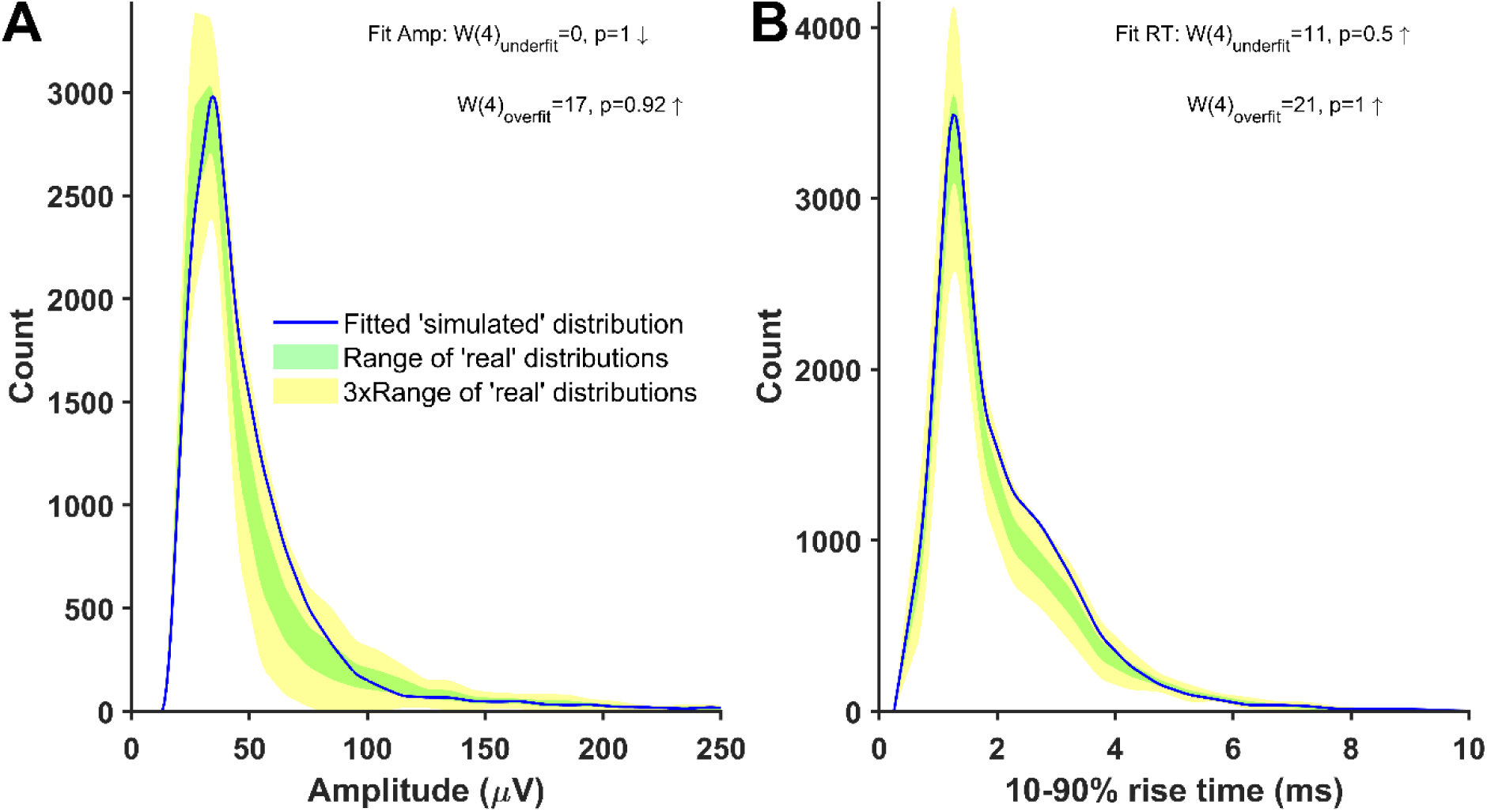
Cell p127c (layer 2/3), distributions of mini-like events detected in a ’noise with simulated minis’ voltage trace that closely matched ‘noise with real minis‘ recordings. Parameters of simulated minis controlled by GA. Down arrow: ‘simulated’ SAD < ‘real’ SAD (up: ‘simulated’ > ‘real’; Suppl. Fig. 15).

**Supplementary Figure 24:**
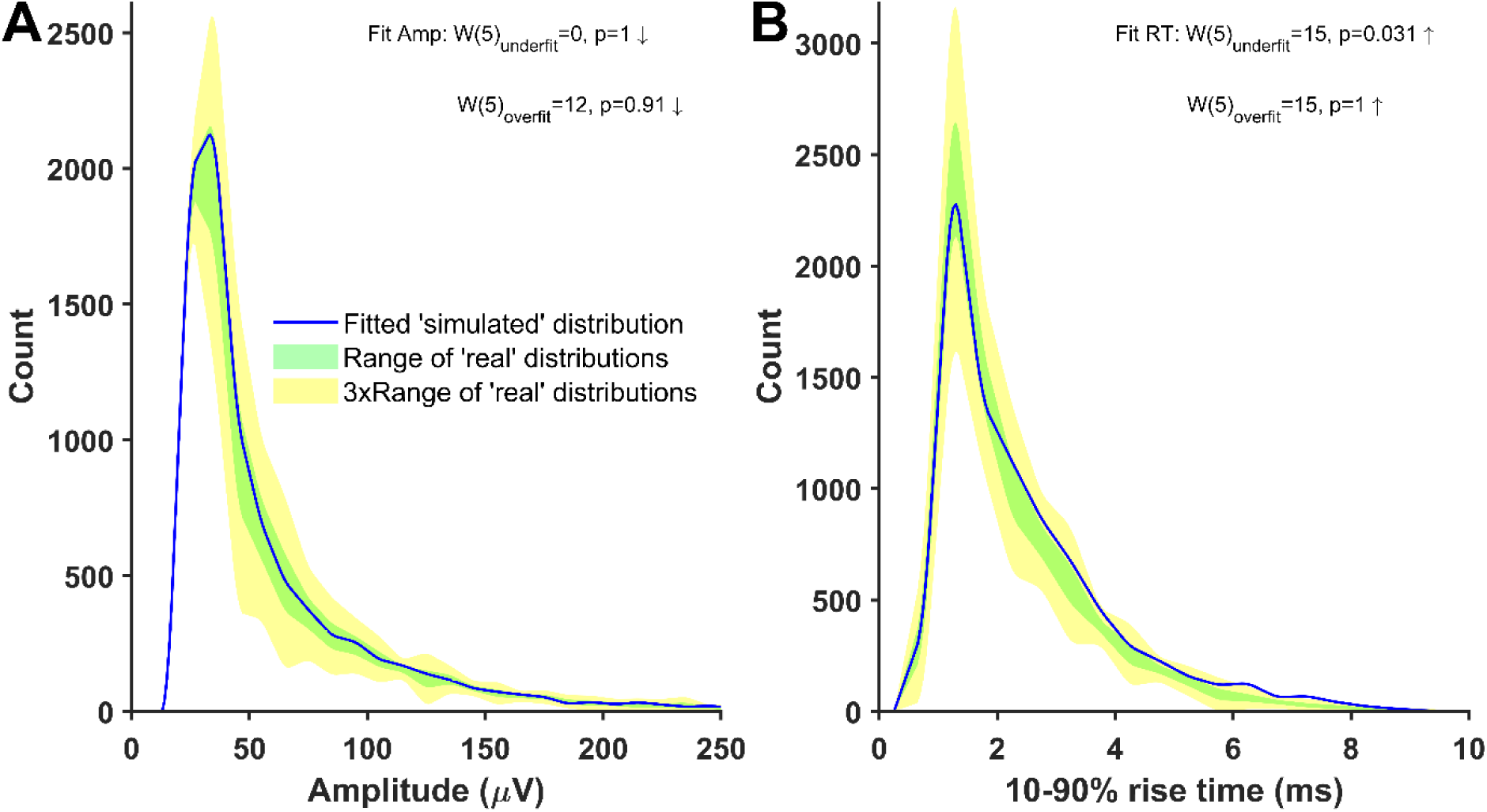
Cell p128c (layer 2/3), distributions of mini-like events detected in a ’noise with simulated minis’ voltage trace that closely matched ‘noise with real minis‘ recordings. Parameters of simulated minis controlled by GA. Down arrow: ‘simulated’ SAD < ‘real’ SAD (up: ‘simulated’ > ‘real’; Suppl. Fig. 15).

**Supplementary Figure 25:**
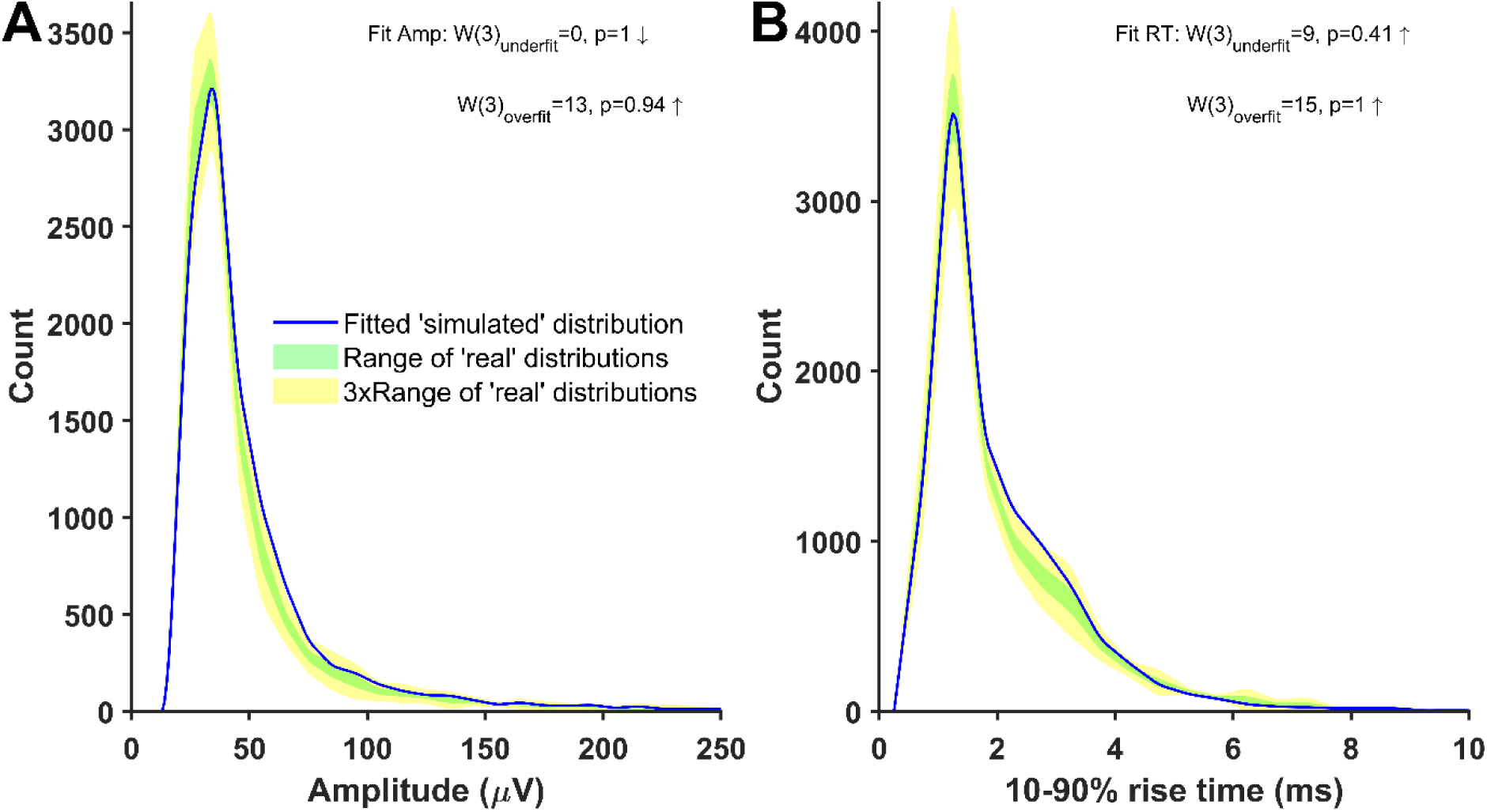
Cell p129a (layer 5), distributions of mini-like events detected in a ’noise with simulated minis’ voltage trace that closely matched ‘noise with real minis‘ recordings. Parameters of simulated minis controlled by GA. Down arrow: ‘simulated’ SAD < ‘real’ SAD (up: ‘simulated’ > ‘real’; Suppl. Fig. 15).

**Supplementary Figure 26:**
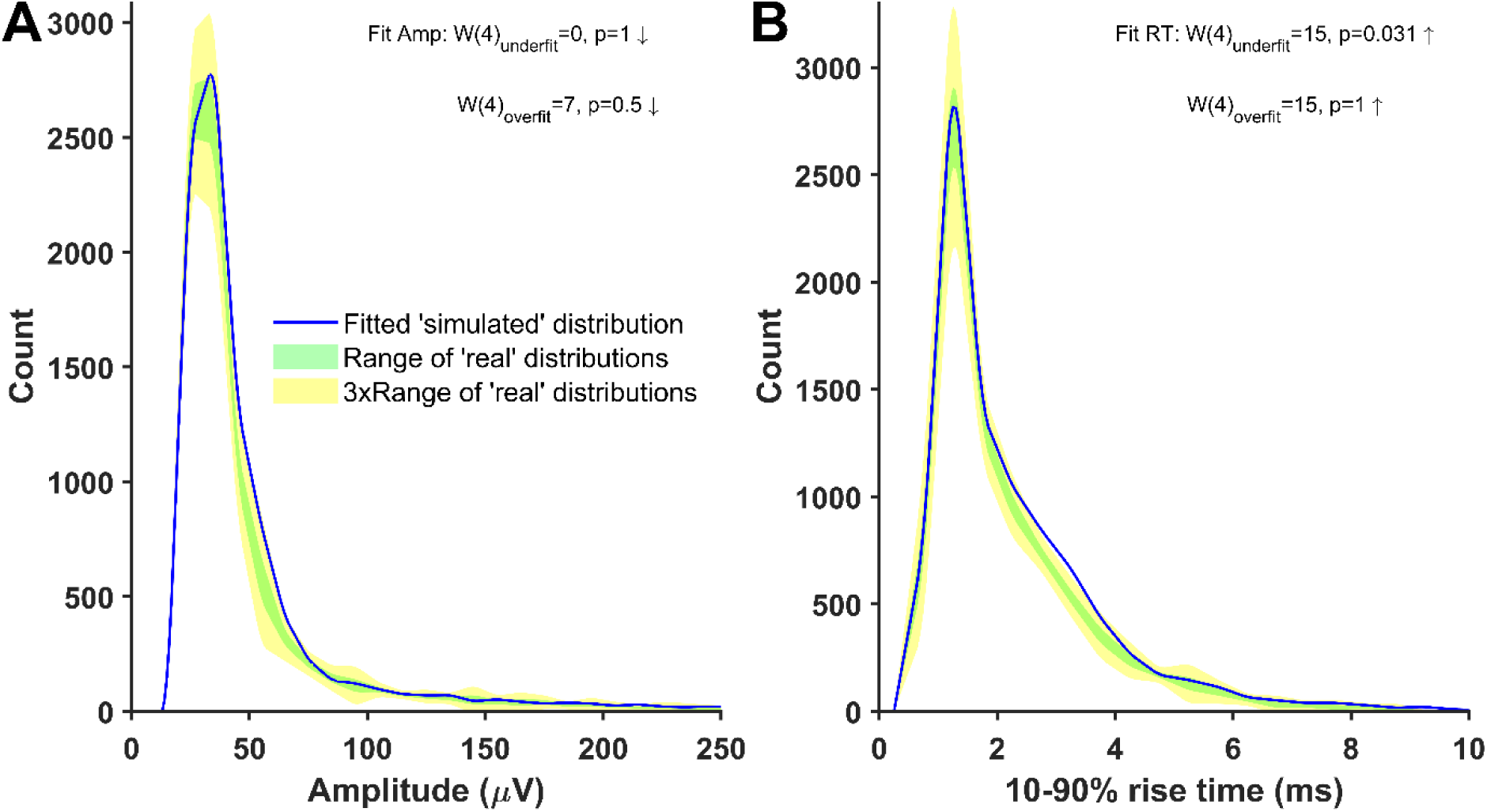
Cell p131a (layer 2/3), distributions of mini-like events detected in a ’noise with simulated minis’ voltage trace that closely matched ‘noise with real minis‘ recordings. Parameters of simulated minis controlled by GA. Down arrow: ‘simulated’ SAD < ‘real’ SAD (up: ‘simulated’ > ‘real’; Suppl. Fig. 15).

**Supplementary Figure 27:**
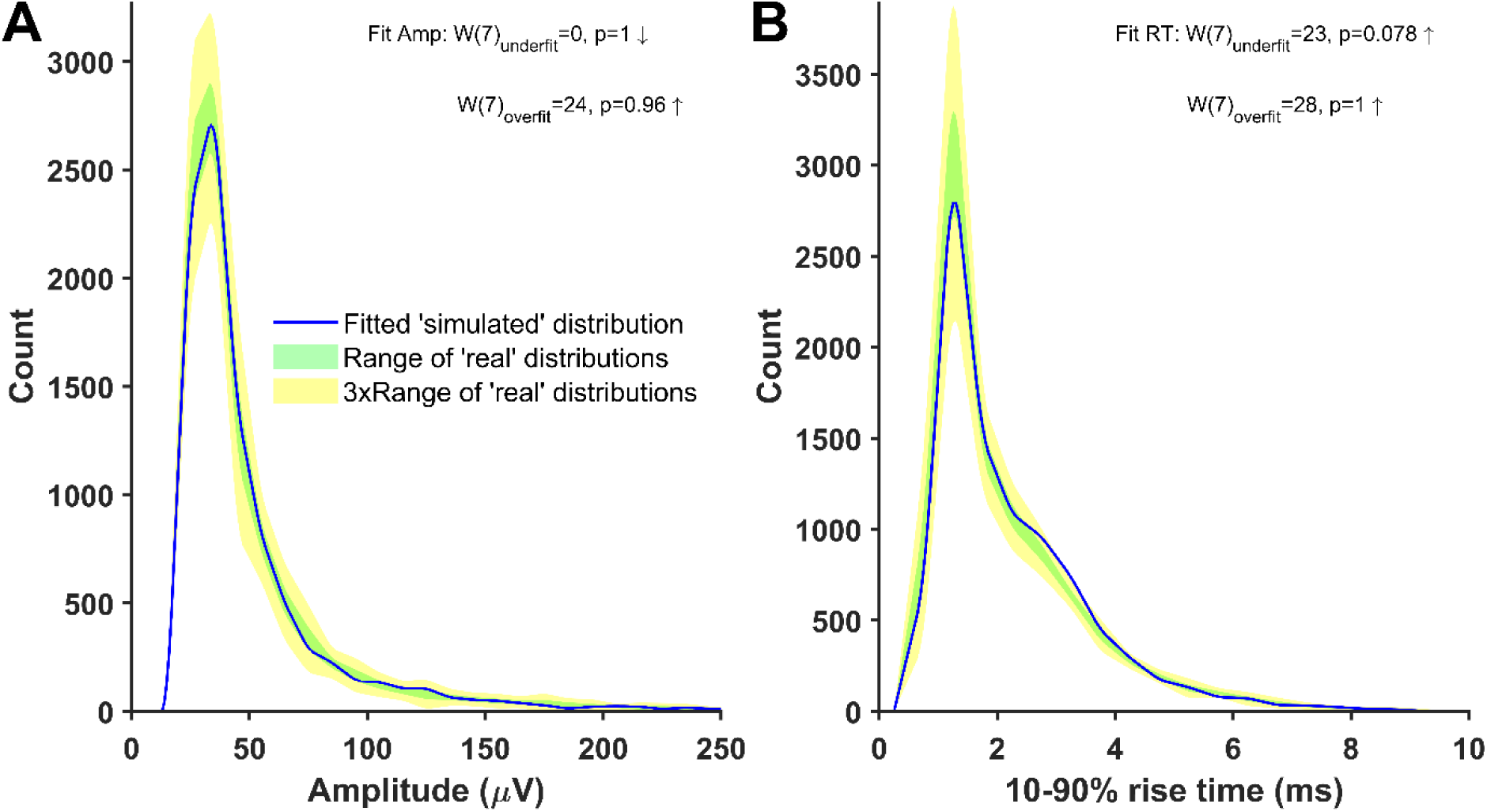
Cell p131c (layer 2/3), distributions of mini-like events detected in a ’noise with simulated minis’ voltage trace that closely matched ‘noise with real minis‘ recordings. Parameters of simulated minis controlled by GA. Down arrow: ‘simulated’ SAD < ‘real’ SAD (up: ‘simulated’ > ‘real’; Suppl. Fig. 15).

**Supplementary Figure 28:**
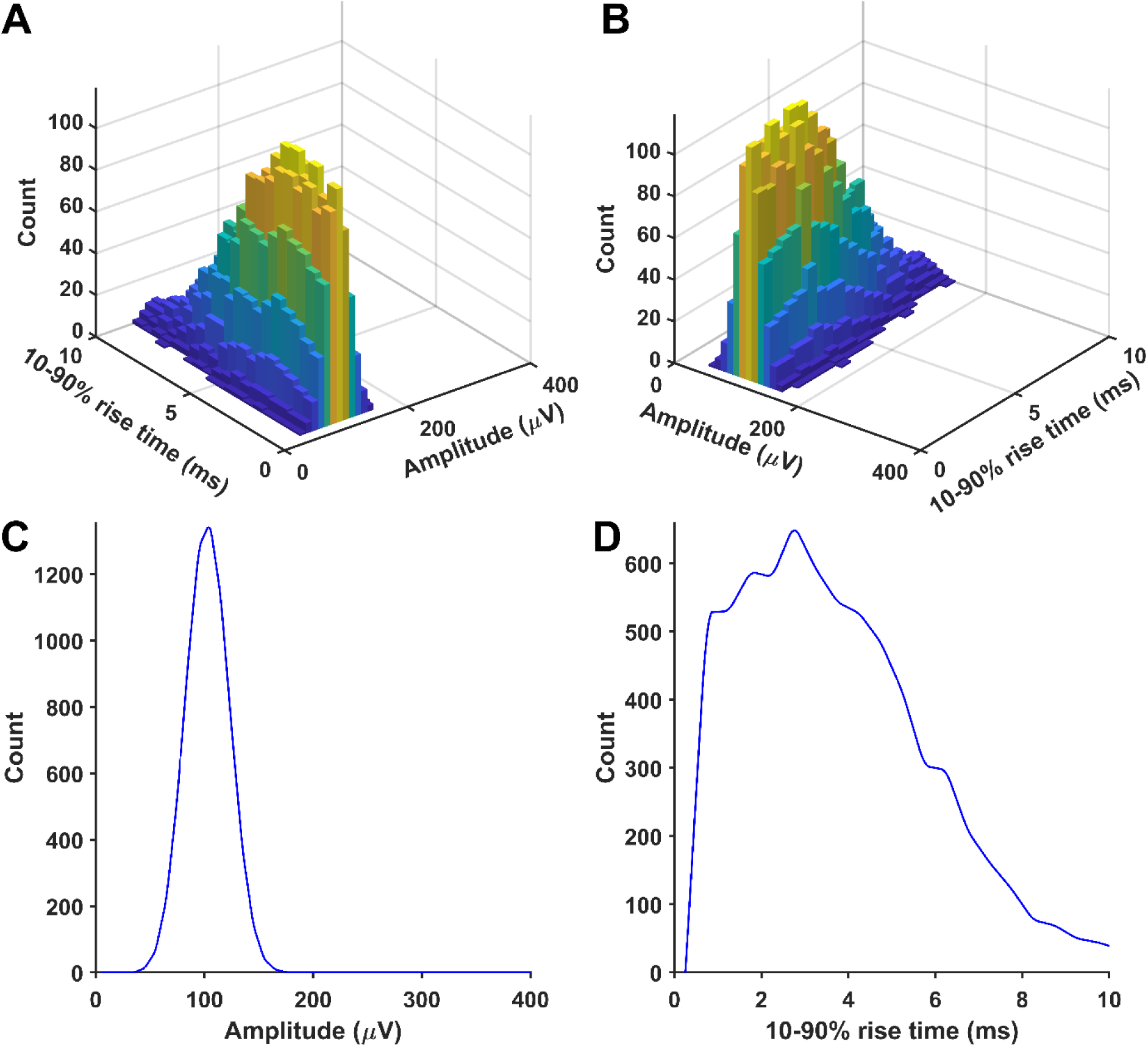
An example of a distribution of randomly selected simulated minis that was used to produce a close match between distributions of mini-like events detected in a ’noise with simulated minis’ voltage trace (40 µV lower limit on simulated amplitudes) and a ’noise with real minis’ recording for cell p103a (layer 5). (A) and (B) Joint amplitude and 10-90% rise time distribution of simulated minis (two views). (C) ‘Marginal’ amplitude and (D) 10-90% rise time distributions (all 2-D bins projected onto that axis).

**Supplementary Figure 29:**
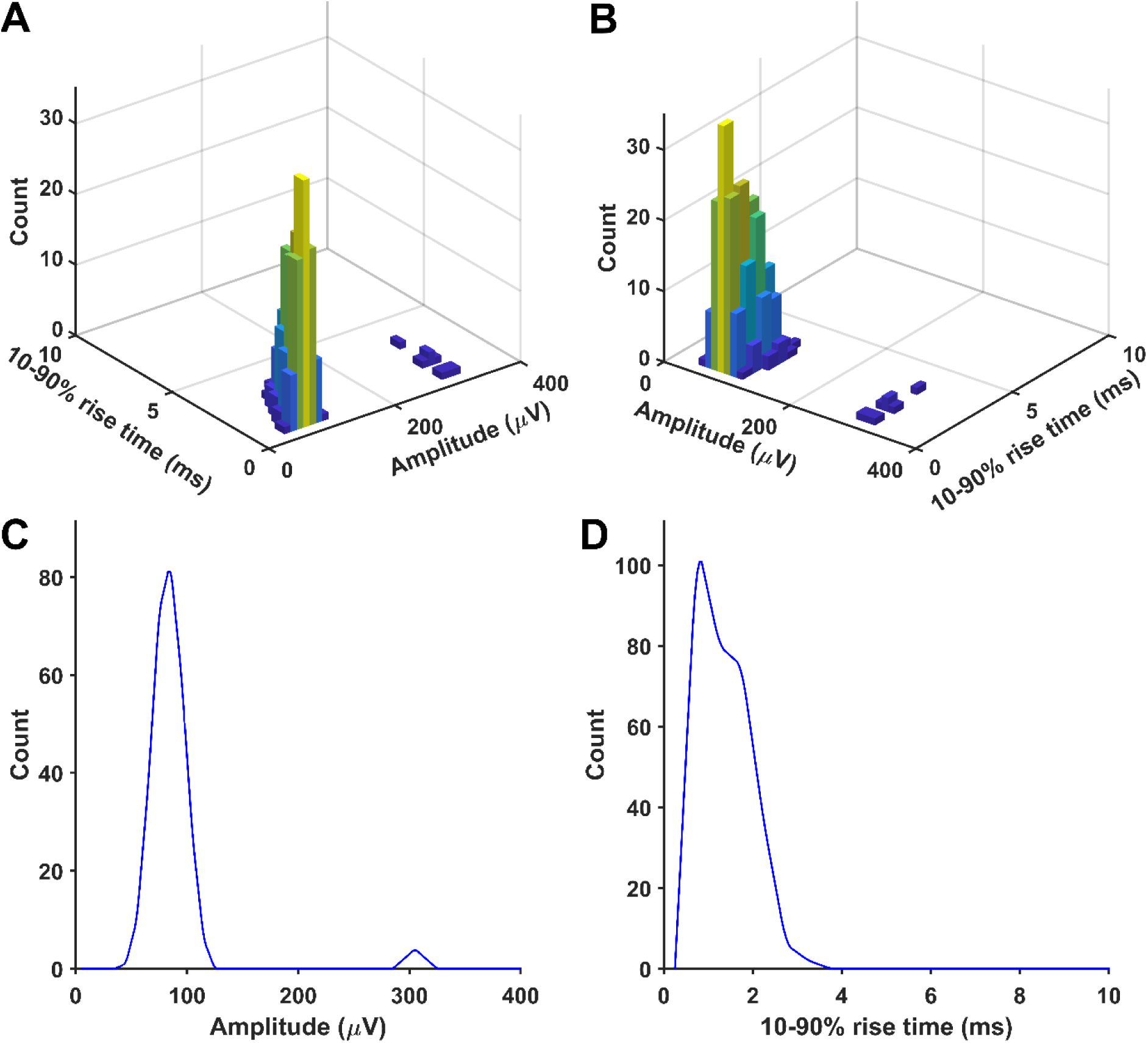
An example of a distribution of randomly selected simulated minis that was used to produce a close match between distributions of mini-like events detected in a ’noise with simulated minis’ voltage trace (40 µV lower limit on simulated amplitudes) and a ’noise with real minis’ recording for cell p106b (layer 5). Panels as for Supplementary Figure 28.

**Supplementary Figure 30:**
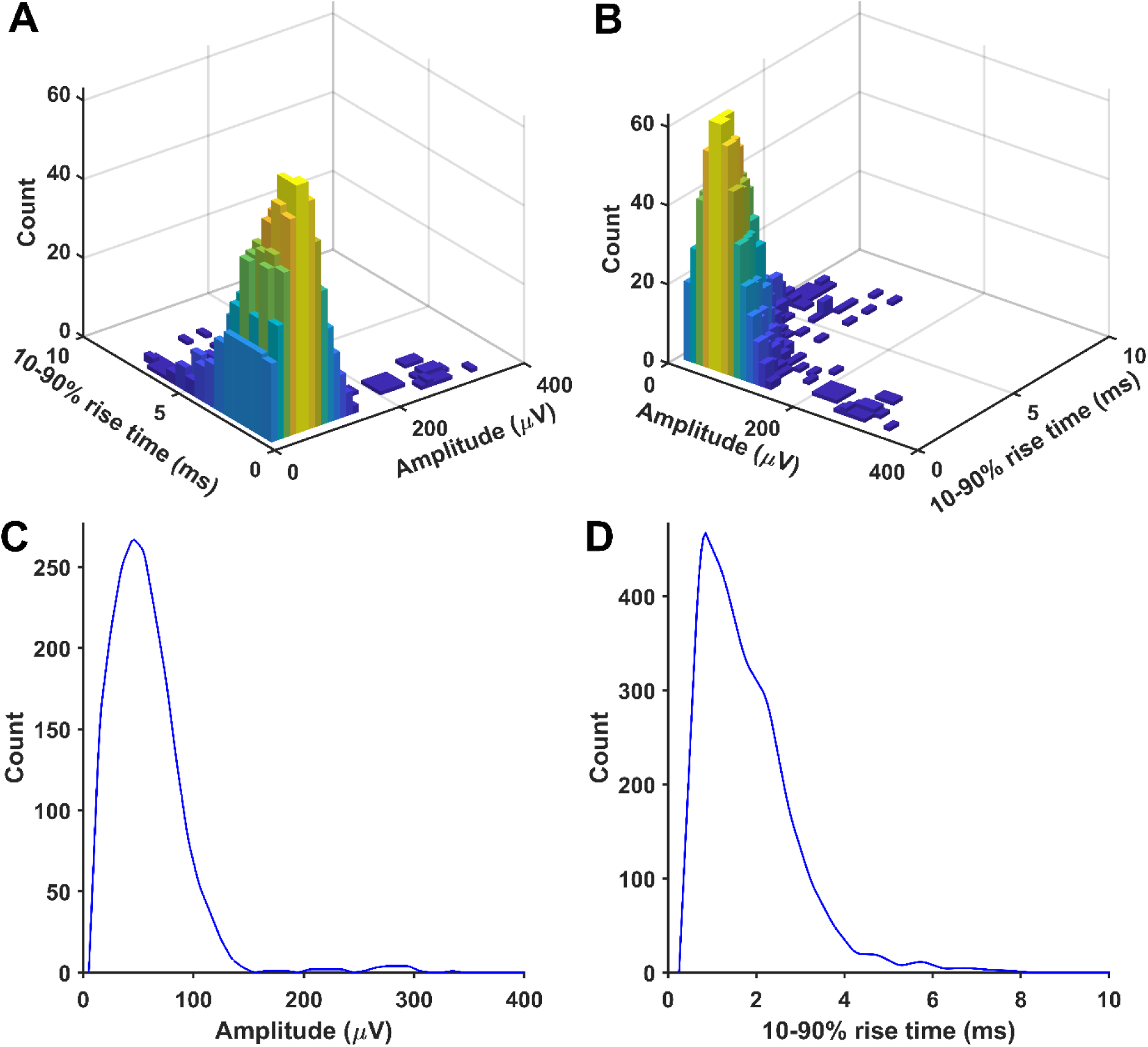
An example of a distribution of randomly selected simulated minis that was used to produce a close match between distributions of mini-like events detected in a ’noise with simulated minis’ voltage trace (10 µV lower limit on simulated amplitudes) and a ’noise with real minis’ recording for cell p108a (layer 5). Panels as for Supplementary Figure 28.

**Supplementary Figure 31:**
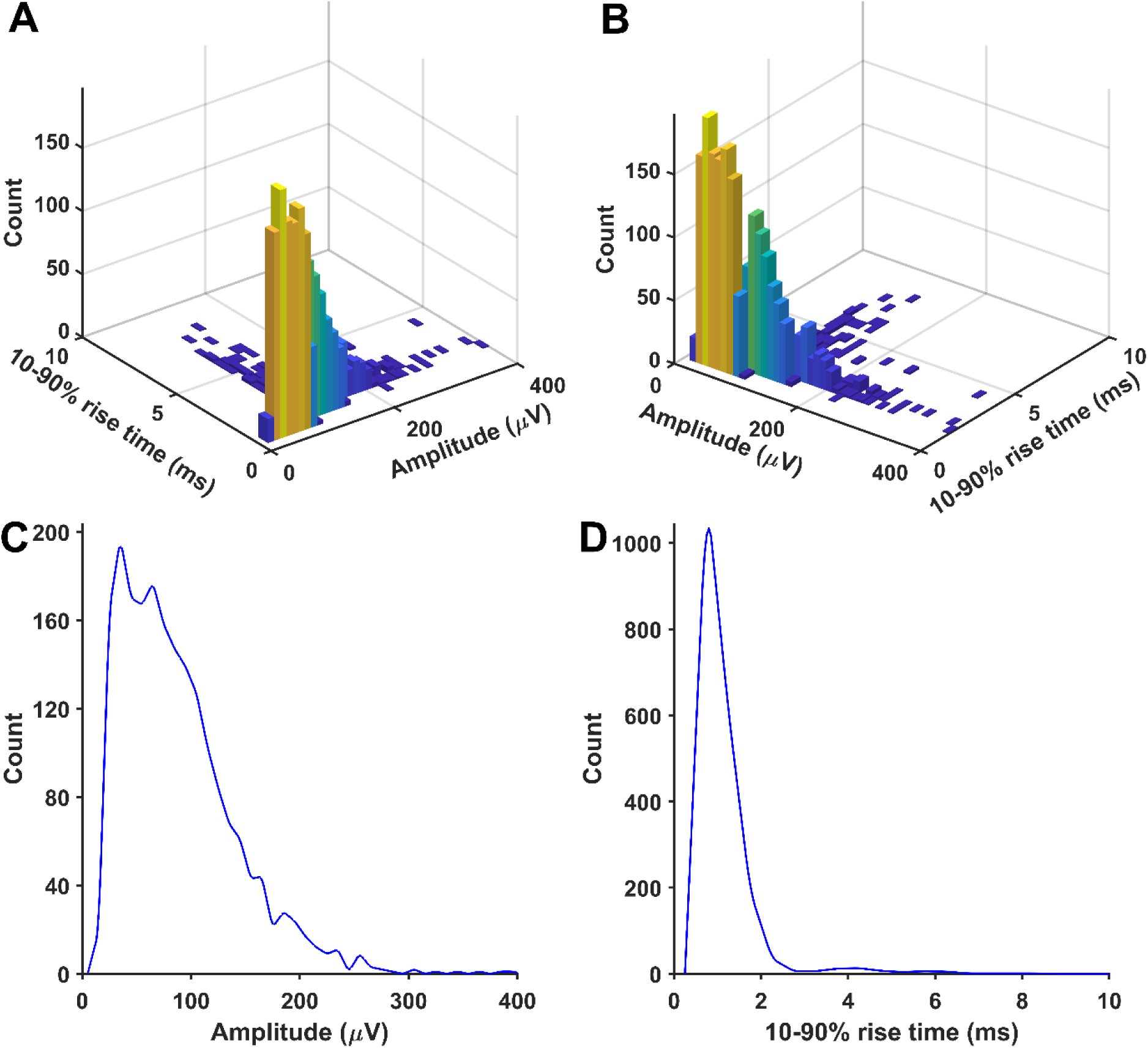
An example of a distribution of randomly selected simulated minis that was used to produce a close match between distributions of mini-like events detected in a ’noise with simulated minis’ voltage trace (10 µV lower limit on simulated amplitudes) and a ’noise with real minis’ recording for cell p108b (layer 5). Panels as for Supplementary Figure 28.

**Supplementary Figure 32:**
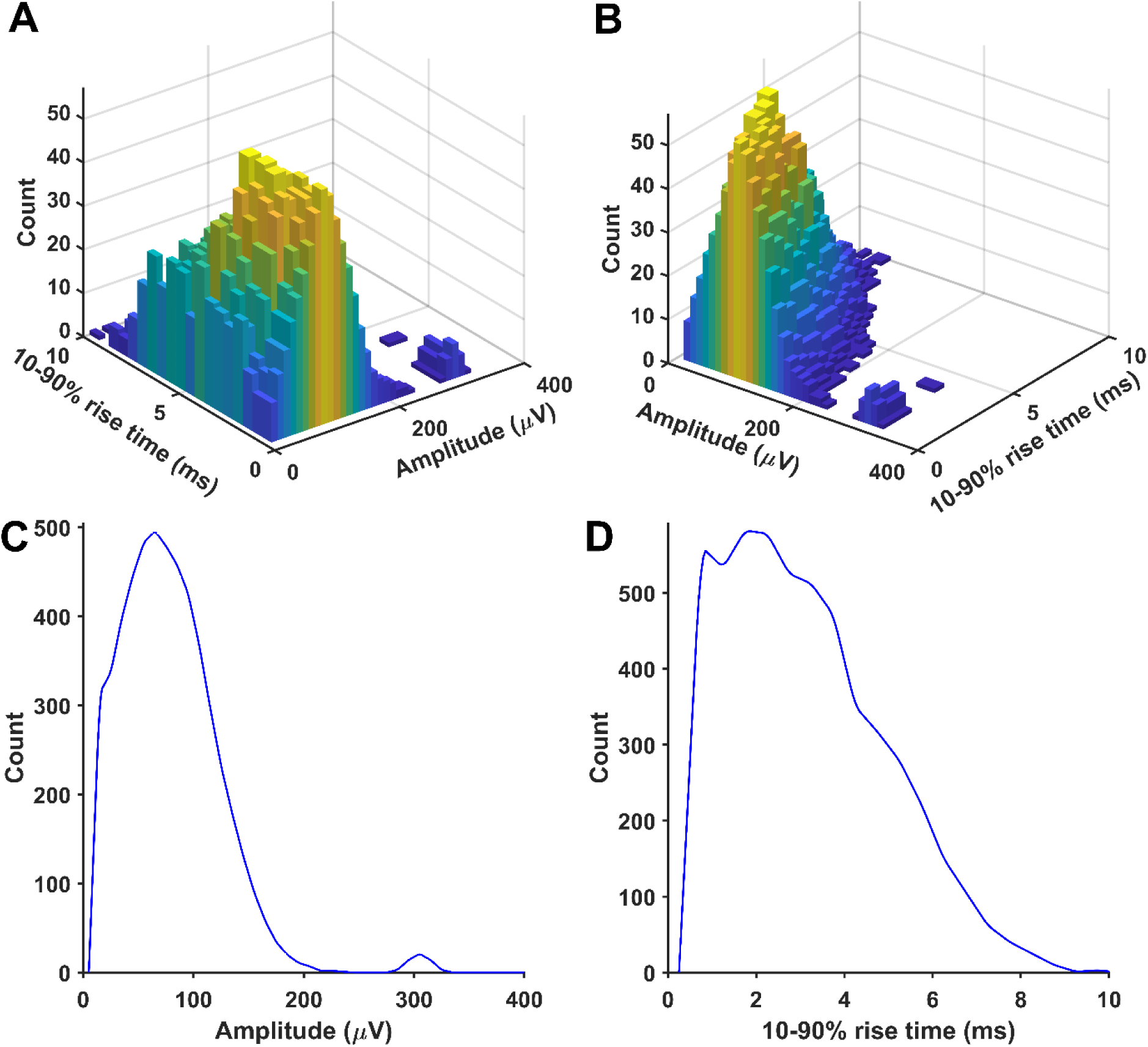
An example of a distribution of randomly selected simulated minis that was used to produce a close match between distributions of mini-like events detected in a ’noise with simulated minis’ voltage trace (10 µV lower limit on simulated amplitudes) and a ’noise with real minis’ recording for cell p108c (layer 5). Panels as for Supplementary Figure 28.

**Supplementary Figure 33:**
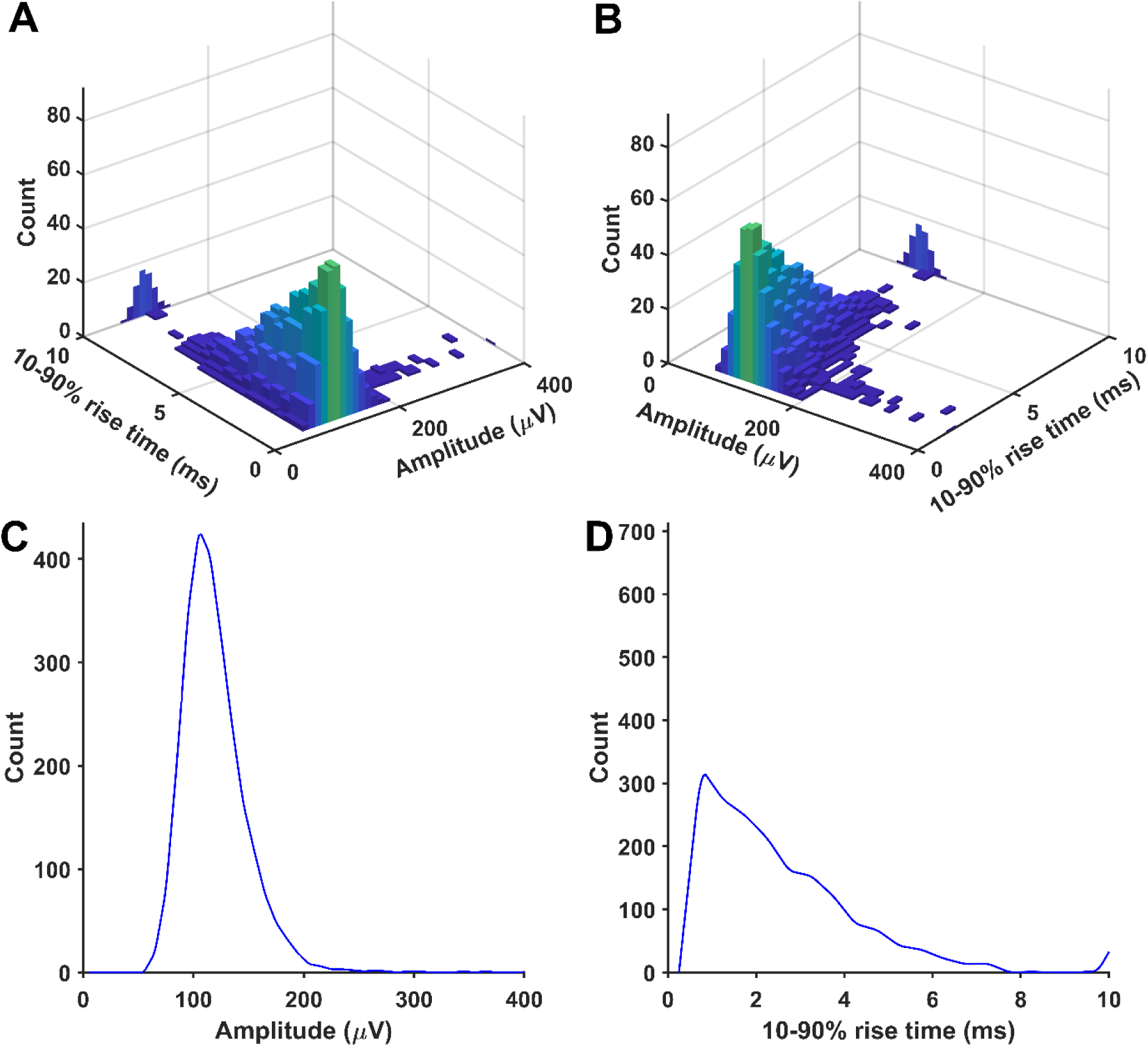
An example of a distribution of randomly selected simulated minis that was used to produce a close match between distributions of mini-like events detected in a ’noise with simulated minis’ voltage trace (60 µV lower limit on simulated amplitudes) and a ’noise with real minis’ recording for cell p120b (layer 2/3). Panels as for Supplementary Figure 28.

**Supplementary Figure 34:**
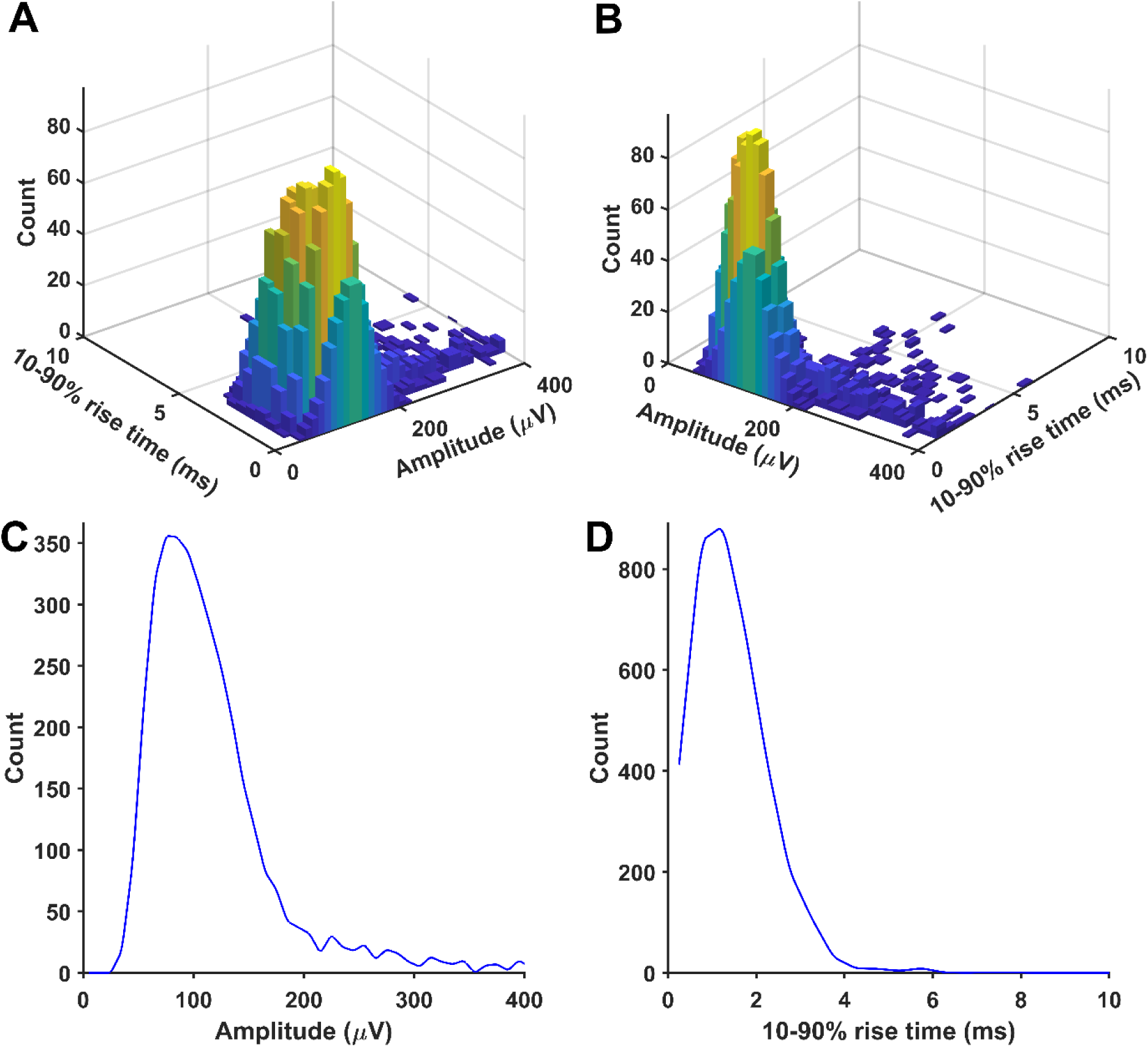
An example of a distribution of randomly selected simulated minis that was used to produce a close match between distributions of mini-like events detected in a ’noise with simulated minis’ voltage trace (30 µV lower limit on simulated amplitudes) and a ’noise with real minis’ recording for cell p122a (layer 2/3). Panels as for Supplementary Figure 28.

**Supplementary Figure 35:**
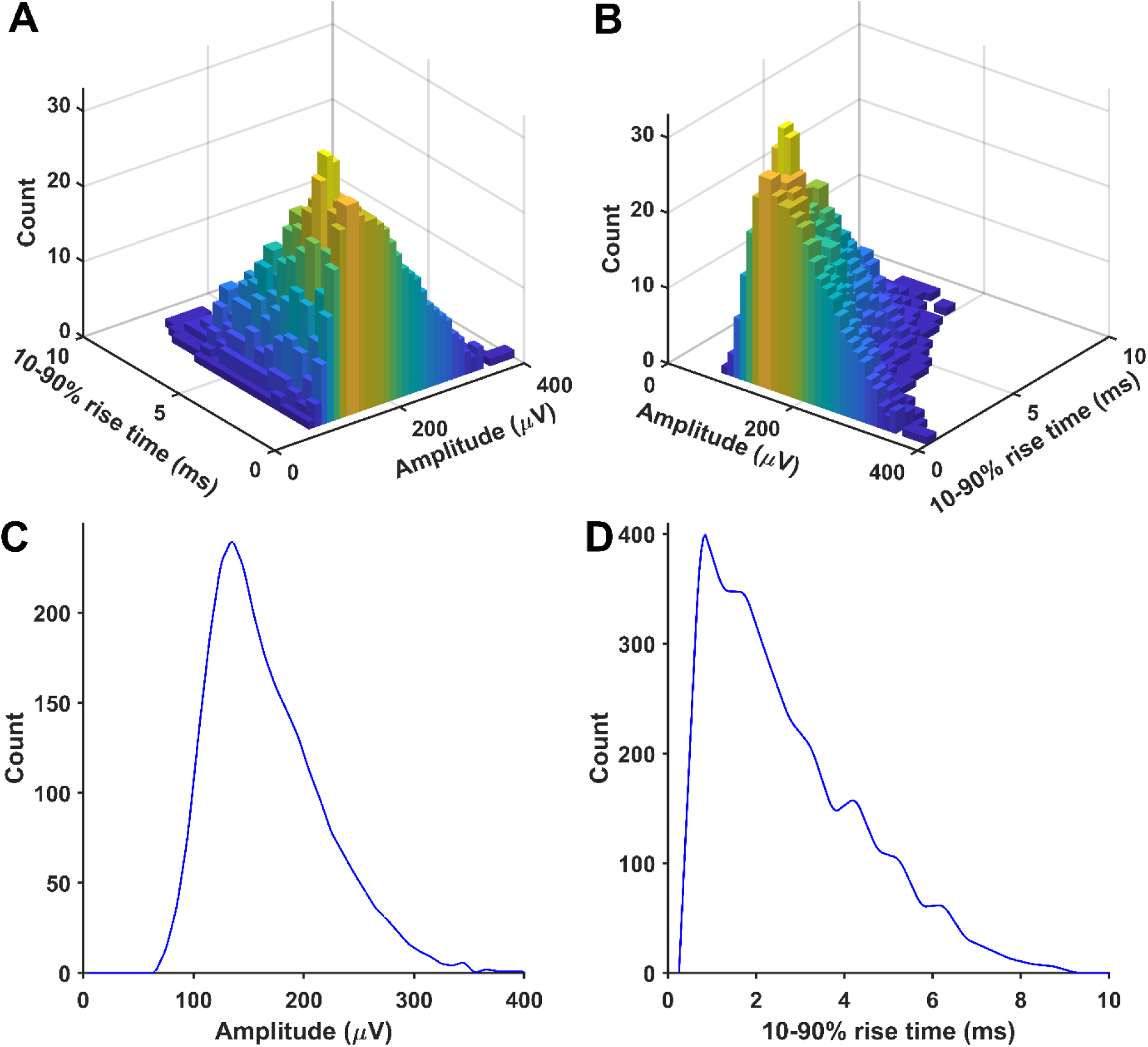
An example of a distribution of randomly selected simulated minis that was used to produce a close match between distributions of mini-like events detected in a ’noise with simulated minis’ voltage trace (40 µV lower limit on simulated amplitudes) and a ’noise with real minis’ recording for cell p124b (layer 5). Panels as for Supplementary Figure 28.

**Supplementary Figure 36:**
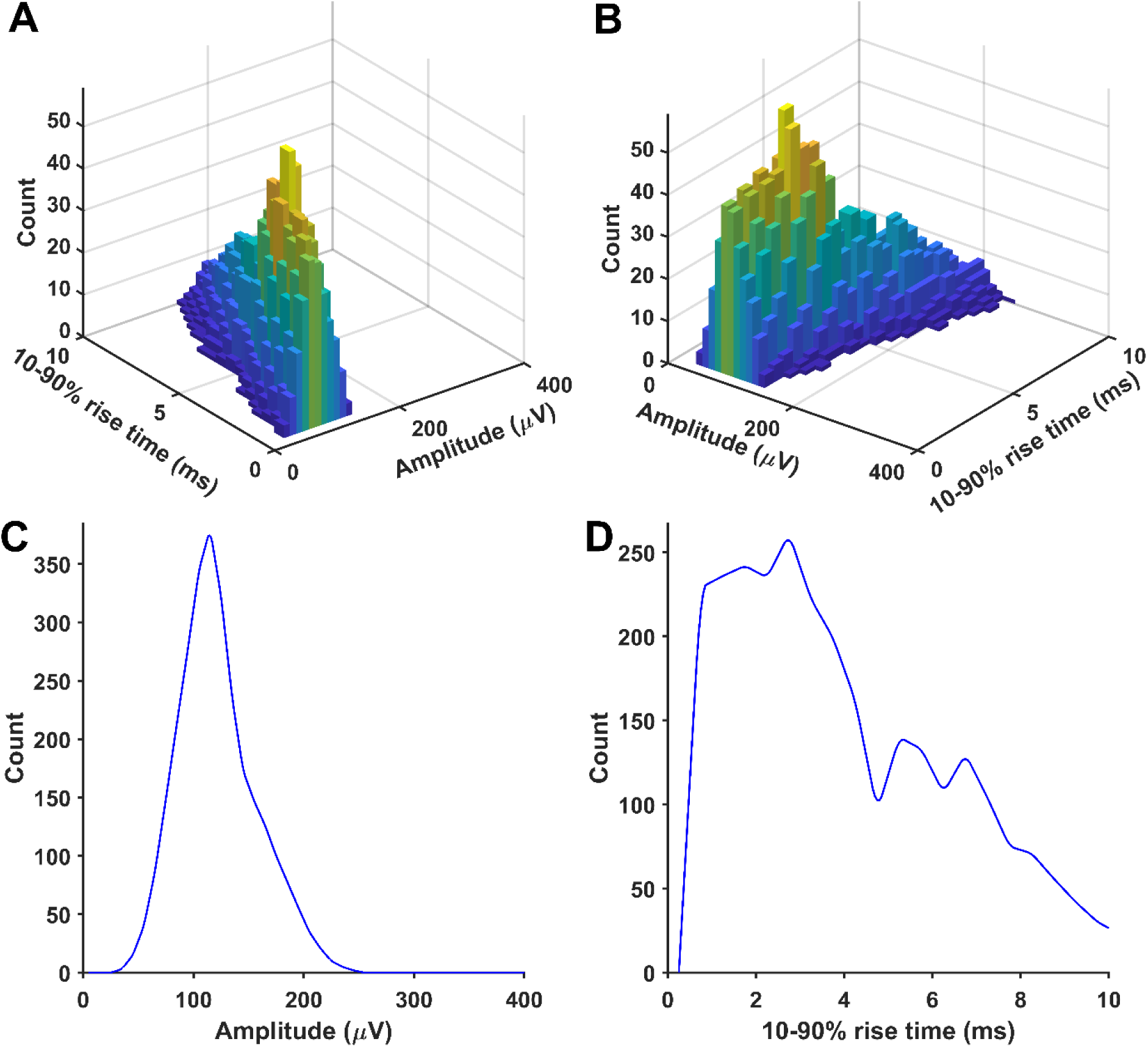
An example of a distribution of randomly selected simulated minis that was used to produce a close match between distributions of mini-like events detected in a ’noise with simulated minis’ voltage trace (30 µV lower limit on simulated amplitudes) and a ’noise with real minis’ recording for cell p125a (layer 2/3). Panels as for Supplementary Figure 28.

**Supplementary Figure 37:**
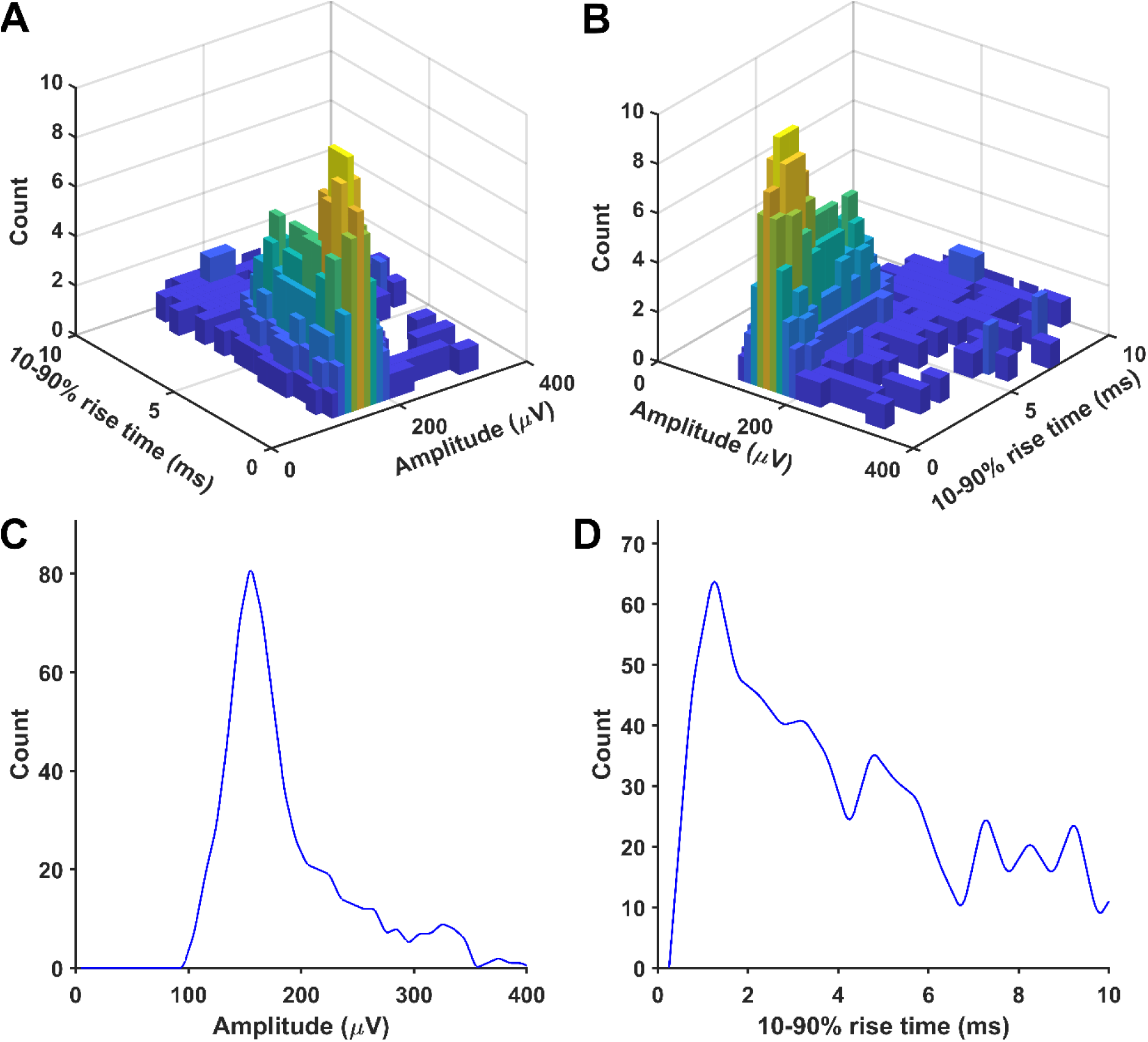
An example of a distribution of randomly selected simulated minis that was used to produce a close match between distributions of mini-like events detected in a ’noise with simulated minis’ voltage trace (100 µV lower limit on simulated amplitudes) and a ’noise with real minis’ recording for cell p127c (layer 2/3). Panels as for Supplementary Figure 28.

**Supplementary Figure 38:**
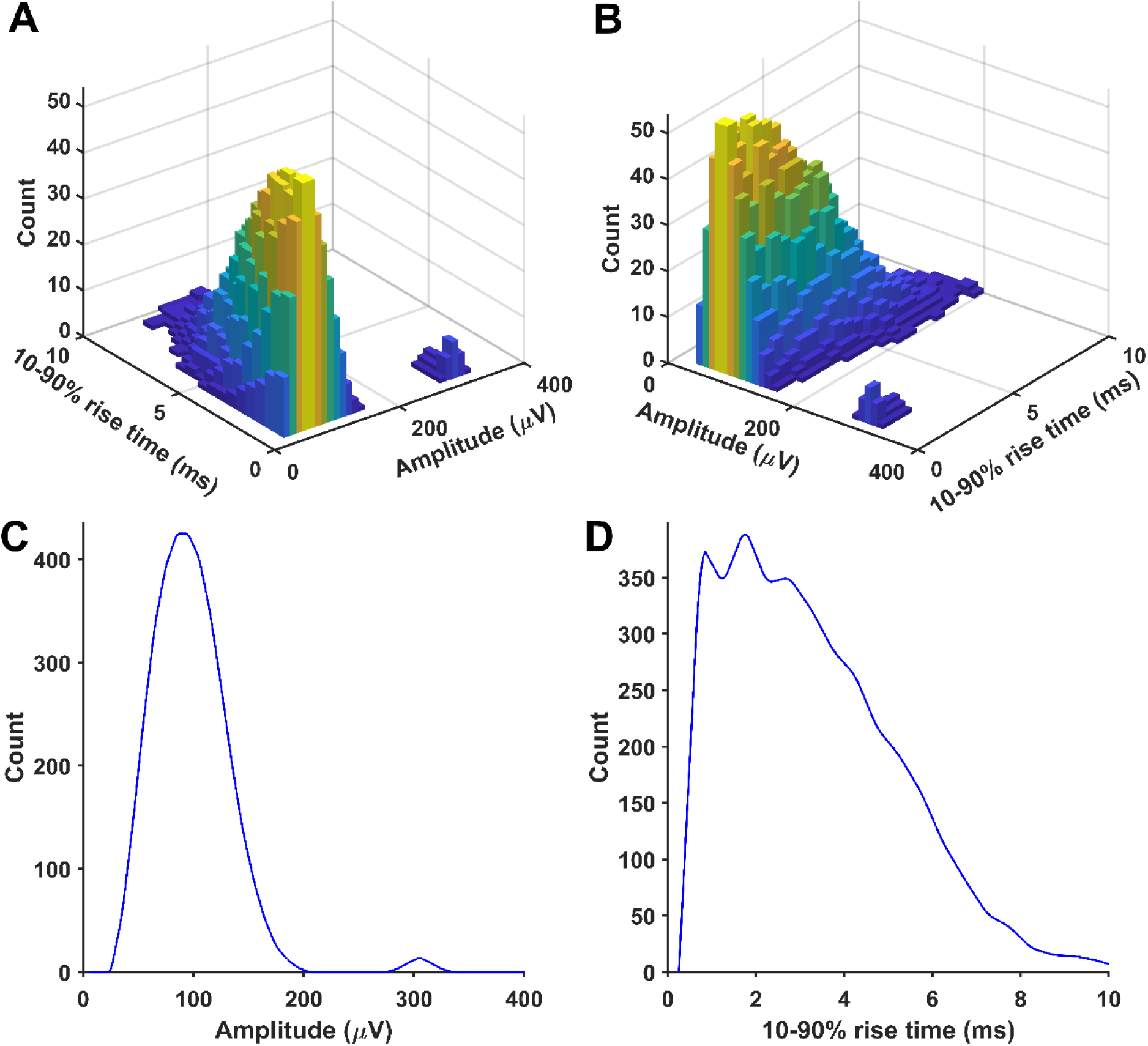
An example of a distribution of randomly selected simulated minis that was used to produce a close match between distributions of mini-like events detected in a ’noise with simulated minis’ voltage trace (30 µV lower limit on simulated amplitudes) and a ’noise with real minis’ recording for cell p128c (layer 2/3). Panels as for Supplementary Figure 28.

**Supplementary Figure 39:**
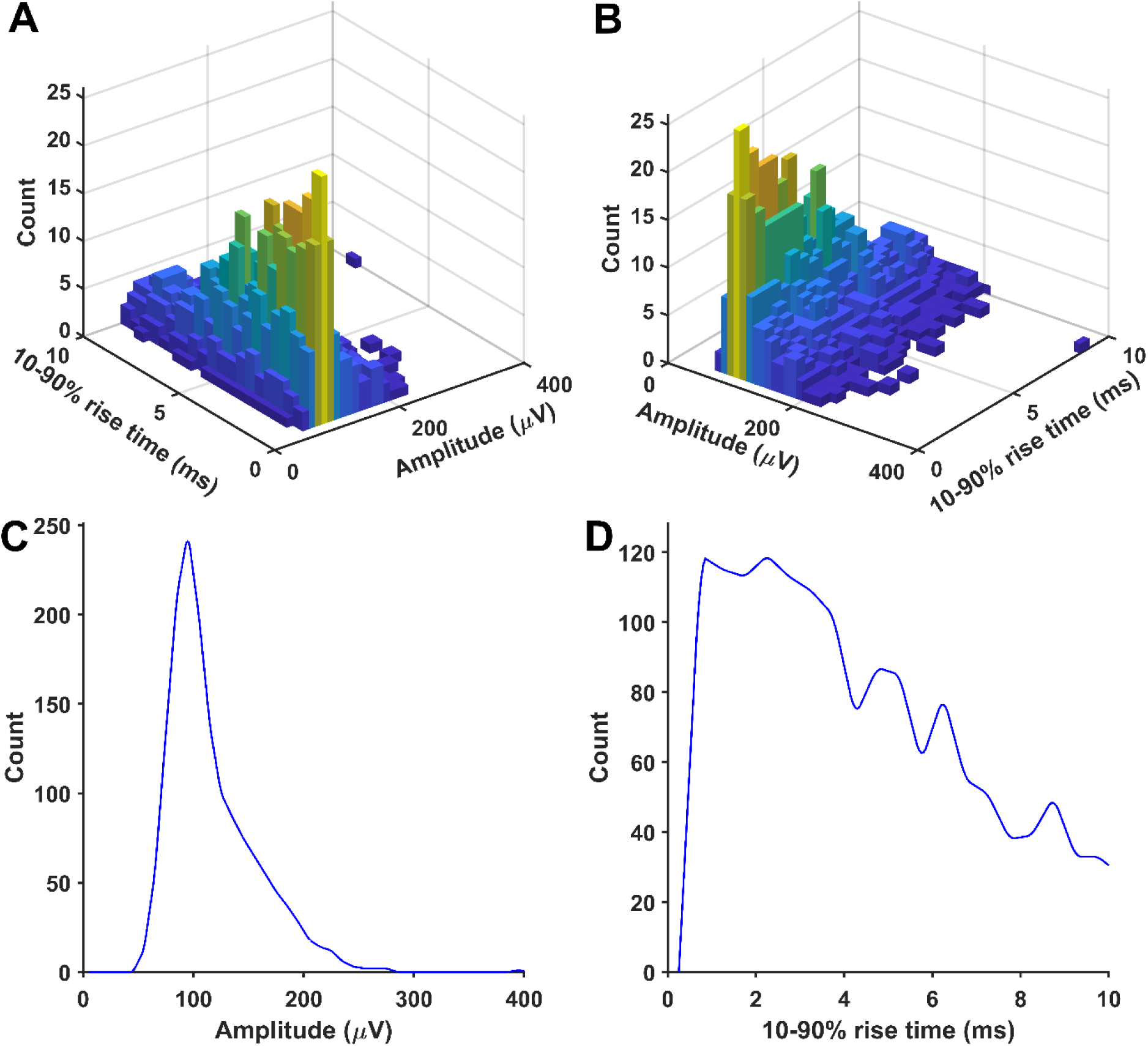
An example of a distribution of randomly selected simulated minis that was used to produce a close match between distributions of mini-like events detected in a ’noise with simulated minis’ voltage trace (50 µV lower limit on simulated amplitudes) and a ’noise with real minis’ recording for cell p129a (layer 5). Panels as for Supplementary Figure 28.

**Supplementary Figure 40:**
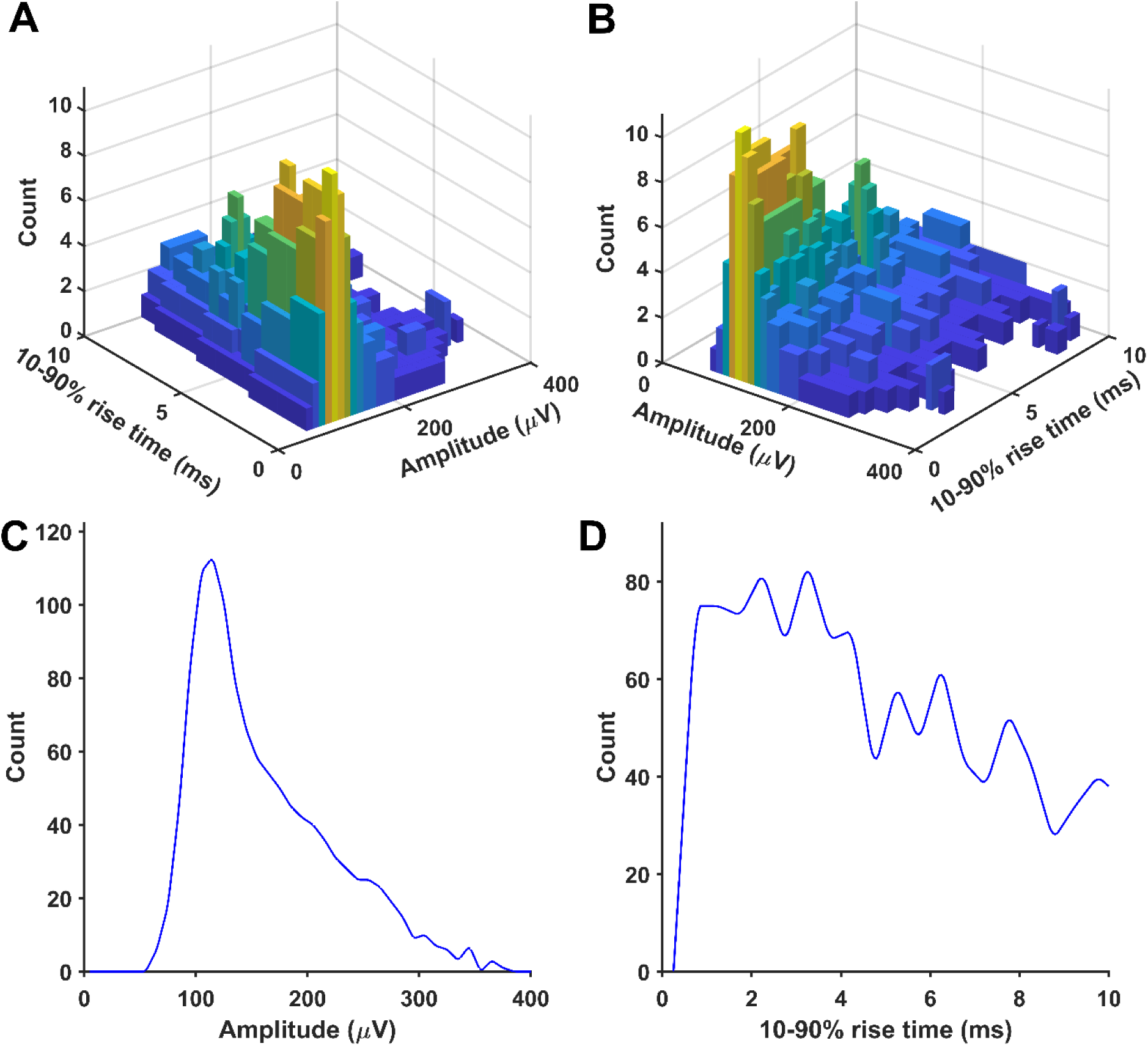
An example of a distribution of randomly selected simulated minis that was used to produce a close match between distributions of mini-like events detected in a ’noise with simulated minis’ voltage trace (60 µV lower limit on simulated amplitudes) and a ’noise with real minis’ recording for cell p131a (layer 2/3). Panels as for Supplementary Figure 28.

